# Mathematical Modelling Indicates Th-cell Targeted Antibody-Dependent Cellular Cytotoxic Is a Crucial Obstacle Hurdling HIV Vaccine Development

**DOI:** 10.1101/2024.02.07.579394

**Authors:** Zhaobin Xu, Qiangcheng Zeng, Dongying Yang, Xiaoguang Sun, Dongqing Wei, Jacques Demongeot, Zanxia Cao

**Affiliations:** School of Life Science, Dezhou University, Dezhou 253023, China; School of Medicine, Dezhou University, Dezhou 253023, China; People hospital of Yucheng City, Dezhou 251299, China; School of Life Sciences and Biotechnology, Shanghai Jiao Tong University, Shanghai 200030, China; Laboratory AGEIS EA 7407, Team Tools for e-Gnosis Medical, Faculty of Medicine, University Grenoble Alpes (UGA), 38700 La Tronche, France; Shandong Provincial Key Laboratory of Biophysics, Institute of Biophysics, Dezhou University, Dezhou 253023, China

**Keywords:** HIV, ADCC effect, Th-cell, mathematical modelling, vaccine development, bispecific antibody

## Abstract

HIV poses a significant threat to human health. Although some progress has been made in the development of an HIV vaccine, there is currently no reported success in achieving an effective and fully functional vaccine for HIV. This highlights the challenges involved in HIV vaccine development. Through mathematical modeling, we have conducted a systematic study on the impact of antibody-dependent cellular cytotoxicity (ADCC) on HIV-specific immune responses. Unlike other viral infections, the ADCC effect following HIV infection may cause significant damage to the follicular center Th cells, leading to apoptosis of follicular center cells and rapid death of effector Th cells. This impedes the generation of neutralizing antibodies and creates barriers to viral clearance, thereby contributing to long-term infection. Another challenge posed by this effect is the substantial reduction in vaccine effectiveness, as effective antigenic substances such as gp120 bind to Th cell surfaces, resulting in the apoptosis of follicular center Th cells due to ADCC, hindering antibody regeneration. To address this issue, we propose the concept of using bispecific antibodies. By genetically editing B cells to insert the bispecific antibody gene, which consists of two parts-one targeting the CD4 binding site of HIV, such as the broadly neutralizing antibody 3BNC117, and the other targeting antibodies against other viruses, such as the spike protein of SARS-CoV-2-we can simultaneously enhance the levels of two pathogen-specific antibodies through stimulation with non-HIV-antigens corresponding to the other part of the chimeric antibody, such as the spike protein. This study contributes to the elucidation of the pathophysiology of HIV, while also providing a theoretical framework for the successful development of an HIV vaccine.

## 1 Introduction

The HIV virus was discovered as early as the 1980s, yet an effective vaccine remains elusive to this day. The challenges in developing an AIDS vaccine are multifaceted. For instance, HIV exhibits rapid mutation, with over 60 main HIV-1 subtypes identified thus far [1–4]. However, progress in research has revealed several conserved regions within the AIDS virus, and antibodies targeting these regions have shown broad protective effects. Interestingly, these conserved regions often concentrate on the receptor binding domain of gp140, which interacts with CD4 [5–10]. Fortunately, these antibodies have been identified in some HIV-infected individuals [11–13], bolstering the confidence of vaccine developers.

However, disappointingly, nearly all human clinical trials have faced failures. Even the most promising RV144 trial only achieved partial protection in the short term [14–17]. Yet, these vaccines often succeed in eliciting antibodies in animal models [18].

The objective of this study is to uncover the reasons behind this phenomenon and propose feasible solutions from a theoretical perspective. Using computational immunology, we simulated the specific immune response process following HIV infection and further developed our previous antibody dynamics model [19]. We focused on investigating the impact of antibody-dependent cellular cytotoxicity (ADCC) on germinal center B cells and T-helper cells. In the context of HIV infection, ADCC could significantly impair Th cells in the germinal center due to high peripheral antibody concentrations caused by proximity. This leads to rapid death of infected Th cells, hindering antibody regeneration. We constructed two models: one coarse-grain model that did not consider cell differentiation. With this model, we elucidated how antigen-specific Th cell immunogenicity affects the host’s specific immune response. We explained how the immune system recognizes foreign substances, triggers the production of high concentrations of specific antibodies, and simultaneously accomplishes the clonal deletion of self-reactive antibodies, as well as the clonal redemption of self-reactive antibodies. Furthermore, using this coarse-grain model, we analyzed the impact of ADCC on effector B cells and Th cells, along with the possible physical mechanisms underlying this effect. To better simulate this phenomenon, we developed a complex model that considers cell differentiation and the specific processes involved in immune responses. We explicitly incorporated B cells and Th cells into this model. With this approach, we explained the following phenomena:

I: Why it is challenging to elicit the production of specific neutralizing antibodies in most infected individuals.
II: Why antibodies targeting the receptor binding domain exhibit better viral inhibition than ordinary antibodies.
III: Why antibody therapy shows promising results.

Finally, we discussed several feasible strategies to overcome the challenges posed by ADCC in HIV vaccine development, with a particular focus on the potential application of bispecific antibodies.

## 2 Methods

### 2.1 A coarse-grained modelling of adaptive immune response (Model 1)

Before delving into the model, it is necessary to introduce the core concepts of adaptive immunity. Although this process is well understood by many immunologists, computational modelers often overlook basic immunological processes, leading to absurd models. While increasing the number of parameters can improve fitting performance, an erroneous model can result in incorrect predictions.

Biological processes are often complex, and it is difficult to mathematically describe each process in detail. Even if we could do so, the parameters of these complex processes are often unobtainable. It is therefore impractical to use mathematical fitting to obtain the parameters of a complex system. We attempt to use the simplest model possible to describe this process. Firstly, antibodies are generated by B cells, and a portion of these antibodies are embedded in the B cell membrane to form membrane proteins, while another portion is secreted into body fluids as free antibodies. These free antibodies display Fc regions that are crucial because when they bind to antigens, the resulting antigen-antibody complexes’ Fc regions guide immune cells such as NK cells and Tc cells to phagocytose and remove them rapidly. Moreover, when these antibodies come into contact with infected cells, the binding of the antibodies to antigen-like substances on the infected cell surface exposes many exposed Fc domains, which guide immune cell-mediated cytotoxicity and quickly identify and lyse the infected cells.

The process of antibody proliferation is as follows: When antibodies embedded in the B cell membrane recognize antigens, their variable regions bind to the antigen, inducing phagocytosis by the B cell. The antigen-antibody complex will eventually be lysed into peptide fragments, presented to Th cells by MHC complexes on the cell surface via presenting proteins, thereby inducing changes in Th cell signaling pathways, resulting in the secretion of various cytokines promoting their own proliferation and that of neighboring B cells, thereby achieving antibody proliferation.

In modern immunology textbooks, the antigen-presenting process is often referred to as APC (antigen-presenting cell) presentation. This definition is understandable because antigen presentation is not primarily accomplished by B cells but by dendritic cells. Still, dendritic cells themselves digest presented antigens via phagocytosis to stimulate specific Th cell proliferation. However, this description can mislead mathematical modelers into ignoring B cells’ antigen-presenting cell properties and treating APC as a unique type of cell. This severs the correct interaction between Th cells and B cells, and they typically believe that, with the help of APC cells, antigen promotes specific Th cell proliferation, which in turn promotes B cell proliferation. In effect, they believe that the antigen promotes antibody production through subsequent linear signaling chains, which is clearly incorrect. The correct process is that only after antibodies bind to antigens are they presented to the corresponding Th cell by B cells. The stronger the binding force between antibodies and antigens, the more peptide molecules that have been lysed will be presented on the B cell surface, and the stronger the B cell’s ability to induce Th cell activation. Since spatial proximity effects occur between B cells and Th cells during the binding process, the cytokines secreted by Th cells will preferentially act on the proliferation of B cells with which they are bound, leading to the massive proliferation of corresponding high-binding activity antibodies. Therefore, the antigen itself does not directly induce antibody proliferation; the antigen-antibody complex induces the massive regeneration of antibodies. In recent years, many studies have also shown that B cells’ antigen-presenting role plays a dominant role in virus-induced specific immunity [20–22]. We define the binding complex formed by B cells and Th cells as a germinal center because it is the engine that promotes antibody and corresponding B cell proliferation. This process is represented by a simple schematic diagram.

Here x denotes the amount of antibody-antigen (virus) complexes, y denotes the total number of antibodies, and z denotes the number of viruses. Six processes are displayed in our model. The first reaction represents the proliferation or the replication of virus with a reaction constant k_l_. The second reaction represents the binding reaction between virus and antibody, with a forward reaction constant k_2_ and reverse constant k_-2_. The third reaction represents the removal of antibody-virus complex with a reaction constant k_3_ with the help of Natural Killer (NK) cells [23]. The fourth reaction represents the induction of new antibody by the antibody-virus complex with a kinetic constant k_4_. In immunology, those virus-antibody complexes are on the surface of B-cells since the antibody are initially produced by B-cells and will attach to the plasma membrane of B-cells. Those complexes would further bind to the helper cells because the antibody has another structure binding region toward those receptors. Those helper cells will present the antigen part, which is a virus in this case, to the T-cells. The physical placement should be B-cells bind to those helper cells and further present themselves close to T-cells. The T-cells will handle those antigen substances; if those substances are not self-originated, they would secrete signal molecules to promote the proliferation or the division of B-cells who attached on them. Therefore, B-cells finally get proliferated, so are the antibodies generated by their B-cells. The fifth reaction represents the degradation of virus with a constant k_5_. The sixth reaction represents the degradation of antibody with a rate constant k_6_.

Here are a few parameters that require special explanation. The parameter k1 represents the viral proliferation capacity, which is related to the properties of the virus. Acute infections often exhibit strong replication ability, as do viruses with smaller genomes, resulting in larger values for the coefficient k1. k2 represents the forward binding coefficient between antigens and antibodies. Antibodies with higher binding activity typically have larger values for k2. On the other hand, k-2 represents the dissociation coefficient. For antibodies with strong binding affinity, this coefficient is often small, and hence this process can be neglected. k3 represents the clearance rate of antigen-antibody complexes. After binding, these complexes present Fc domains on their surface, qualitatively inducing clearance by immune cells. Therefore, the value of k3 is significantly greater than the natural degradation rate of the antigen, k5. k4 represents the induction effect of antibody-antigen complexes on antibody regeneration, which can be influenced by various factors. A larger k4 value indicates a stronger induction effect, promoting robust antibody proliferation. The Th cell immunogenicity of the antigen itself, specifically the primary sequence of the antigen, directly affects the value of k4 [24–25]. If the Th cell immunogenicity of the antigen is extremely low, such as in the case of self-proteins or highly homologous proteins, the k4 value will be small. This prevents B cells with strong binding affinity to this antigen from proliferating extensively. However, antibody-antigen binding accelerates antibody degradation, ultimately leading to clone deletion. Apart from the primary sequence of the antigen, the molecular weight of the antigen also contributes to differences in k4 values. When the primary sequences of antigens are the same, antigens with larger molecular weights often induce higher concentrations of B cells that present antigen peptides, resulting in larger k4 values and a more pronounced antibody proliferation effect. It should be noted that various other factors, such as antigen structure and Th cell susceptibility to the antigen, significantly influence the values of k4 and k3. We will discuss this in detail in the Results section. Specific model description and parameter set can be referred to supplementary materials (model 1.1 to model 1.4).

### 2.2 Modeling HIV Infection with a Cell-Compartment Model (Model 2)

To better describe the process of HIV viral infection, we constructed a complex model that includes multiple compartments, explicitly representing the main cell types involved in the infection process. The model includes several key compartments: susceptible effector Th cells (S-Th), infected effector Th cells (I-Th), virus V, antibodies (A), antibody-virus complexes (A-V), effector B cells (B), germinal center complexes formed by effector B cells presenting virus to susceptible effector Th cells (B-S-Th), germinal center complexes formed by effector B cells presenting virus to infected effector Th cells (B-I-Th), susceptible Th cells (S), and infected Th cells (I).

The difference of this model from the previous simple model is that we assume that antibodies are generated by B cells, and the proliferation of B cells requires the formation of germinal center through the combination of virus-specific B cells and effector Th cells. For HIV infection, its susceptible cells are CD4+ cells, i.e. Th cells. For other virus infections, their susceptible cells are other types of cells, such as SARS-CoV-2 mainly infects epithelial cells with abundant ACE2 receptors. Generally speaking, most viruses do not infect Th cells or effector Th cells. By effector Th cells, we mean those that can produce an immune response to the primary sequence of a specific antigen. Because Th cells are similar to B cells, there is a phenomenon of genetic immune rearrangement in vivo, so not all Th cells are sensitive to short peptide sequences presented after processing a certain antigen. For a specific antigen, there are effector Th cells that are sensitive to it and can form germinal centers with B cells, which we can call antigen-sensitive effector Th cells. As shown in Figure 2, when a virus invades a susceptible cell, the infected cell will be quickly cleared by ADCC with the presence of specific antibodies. The virus will further proliferate in the uncleared infected cells, and ultimately achieve the lysis of infected cells, releasing a large number of virus particles into the body fluid environment. Meanwhile, when B cells expressing certain specific antibodies come into contact with the virus, they will phagocytose and process the virus, and present it to effector Th cells, forming a germinal center. The effector Th cells will release cytokines to promote the proliferation of the germinal center effector Th cells and the B cells bound to them, thereby achieving the mass proliferation of specific antibodies [26–28]. The specific process is represented by Table 1, and the rate of each reaction step is represented by Table 2. infection rate is α(S-Th)V/(V+ k_m_), which differs from the classic expression α(S-Th)V significantly, because the process of virus infecting cells is not one-to-one correspondence, multiple viruses can infect one cell simultaneously. Assuming the probability of a single virus infecting a cell is α, the average number of infected cells after V viruses infect S cells is 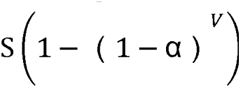, which differs significantly from the αSV model, especially when V is large. Using the Michaelis-Menten equation to represent the rate of virus infecting susceptible cells as 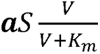 can avoid the phenomenon of target-cell depletion. The second reaction represents the ADCC clearance effect of infected cells in the presence of antibody A, and its clearance rate is also represented by a formula similar to the Michalis-Menton equation, which is 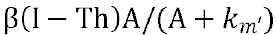, where the higher the concentration of antibody A, the faster the clearance rate, and the maximum clearance rate is 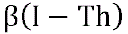. The third reaction represents the rate of germinal center formation between B cells and healthy effector Th cells after binding to the virus, which is 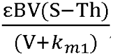, and the germinal center can also dissociate, with a dissociation rate of 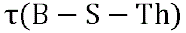. The fourth reaction represents the formation of infectious germinal centers of B cells with infected effector Th cells after binding to the virus, whose forward reaction rate is 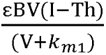, and the reverse dissociation rate is 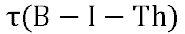.

**Figure 1:**
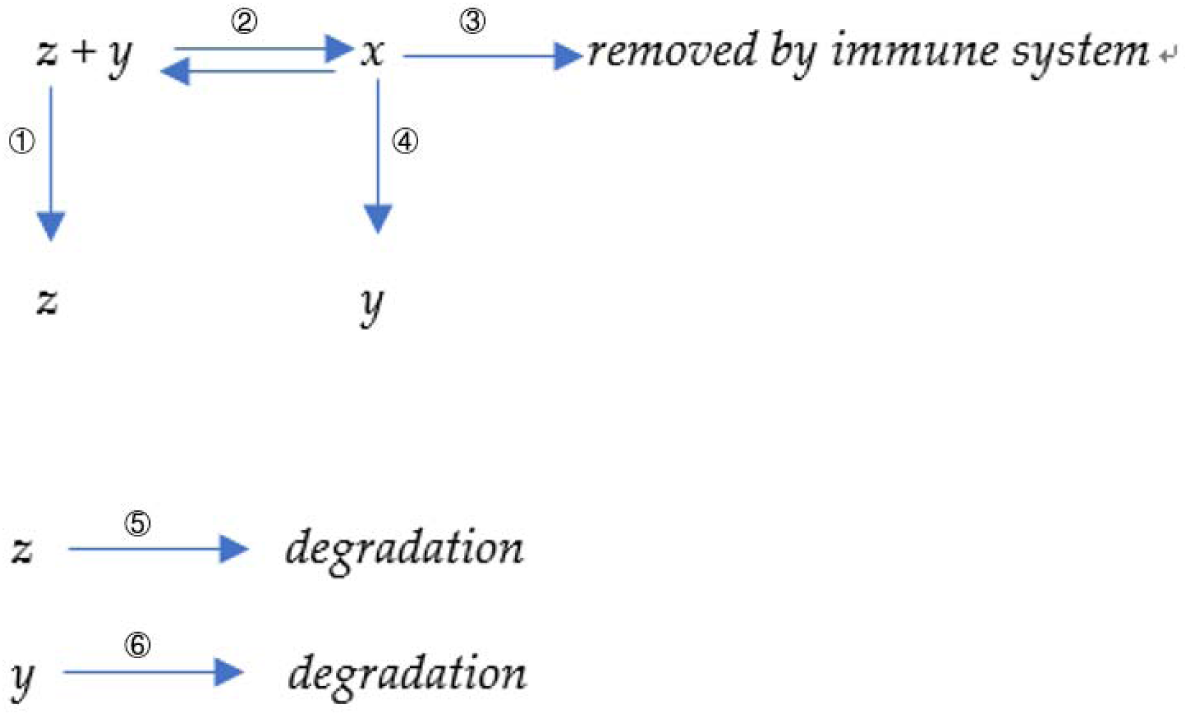
A diagram of simple model of virus-antibody interaction

**Figure 2:**
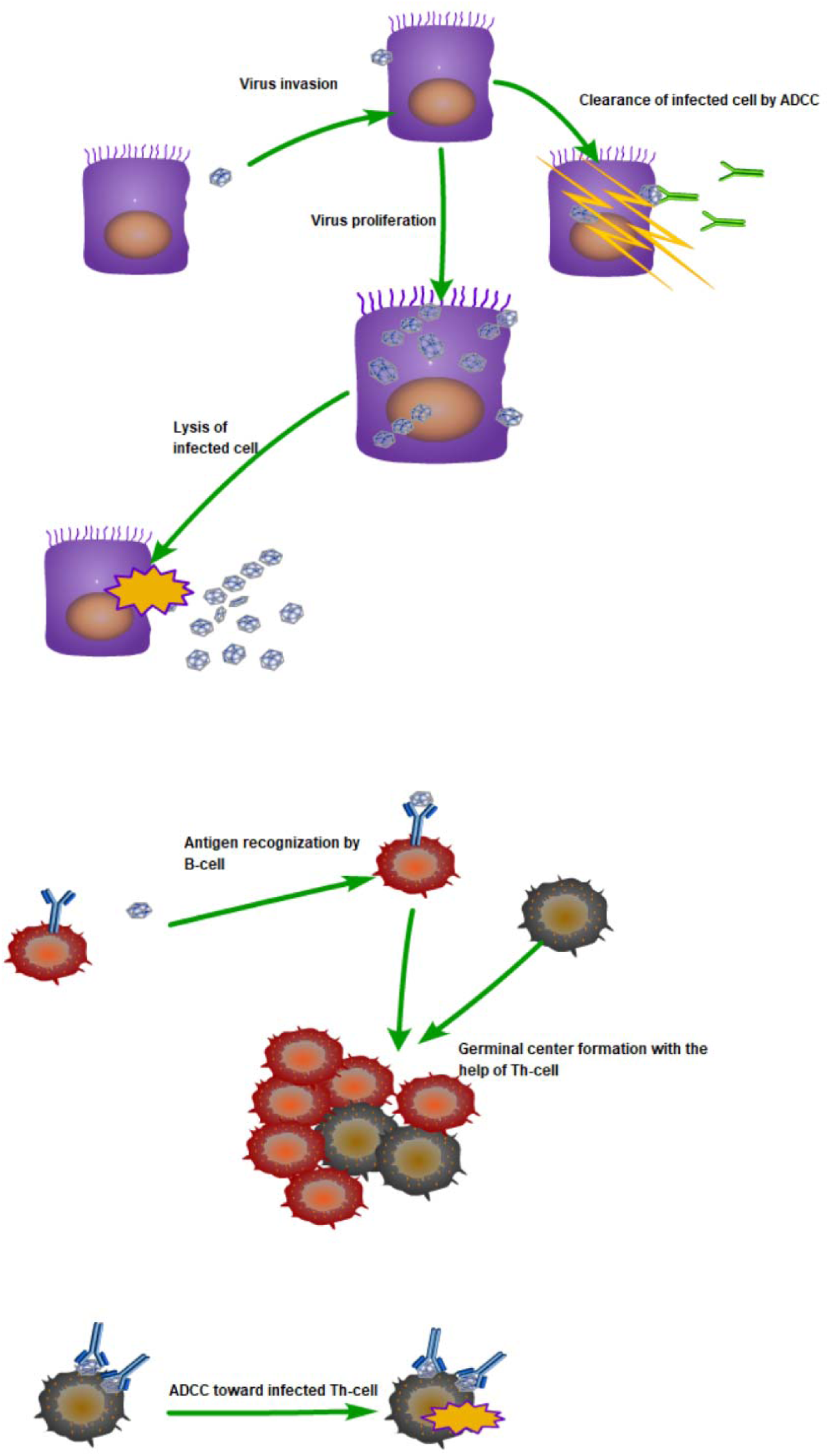
A simple diagram of compartment model

**Table 1:**
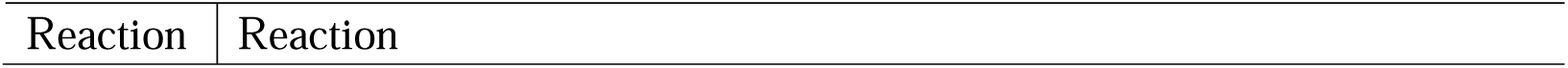
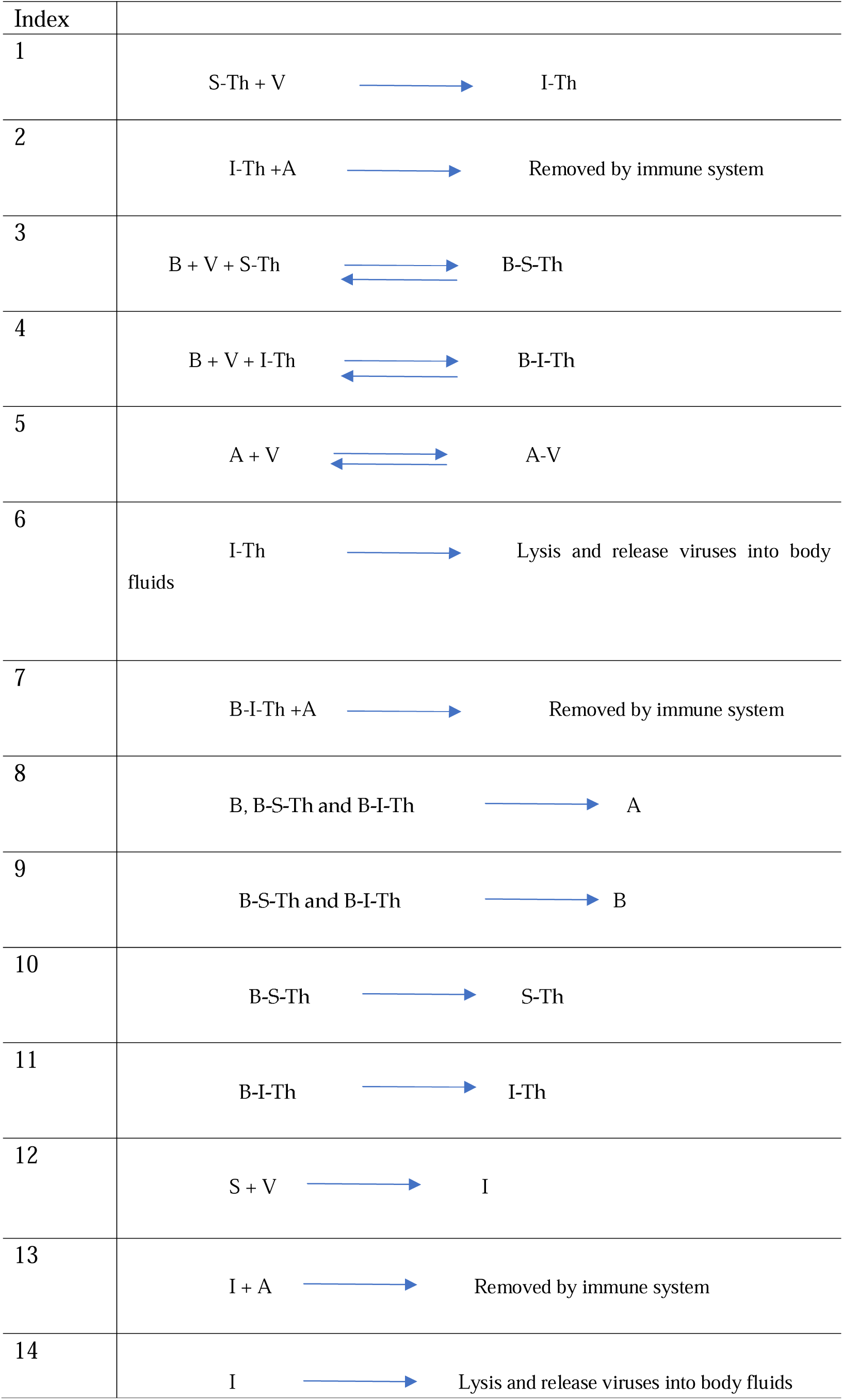
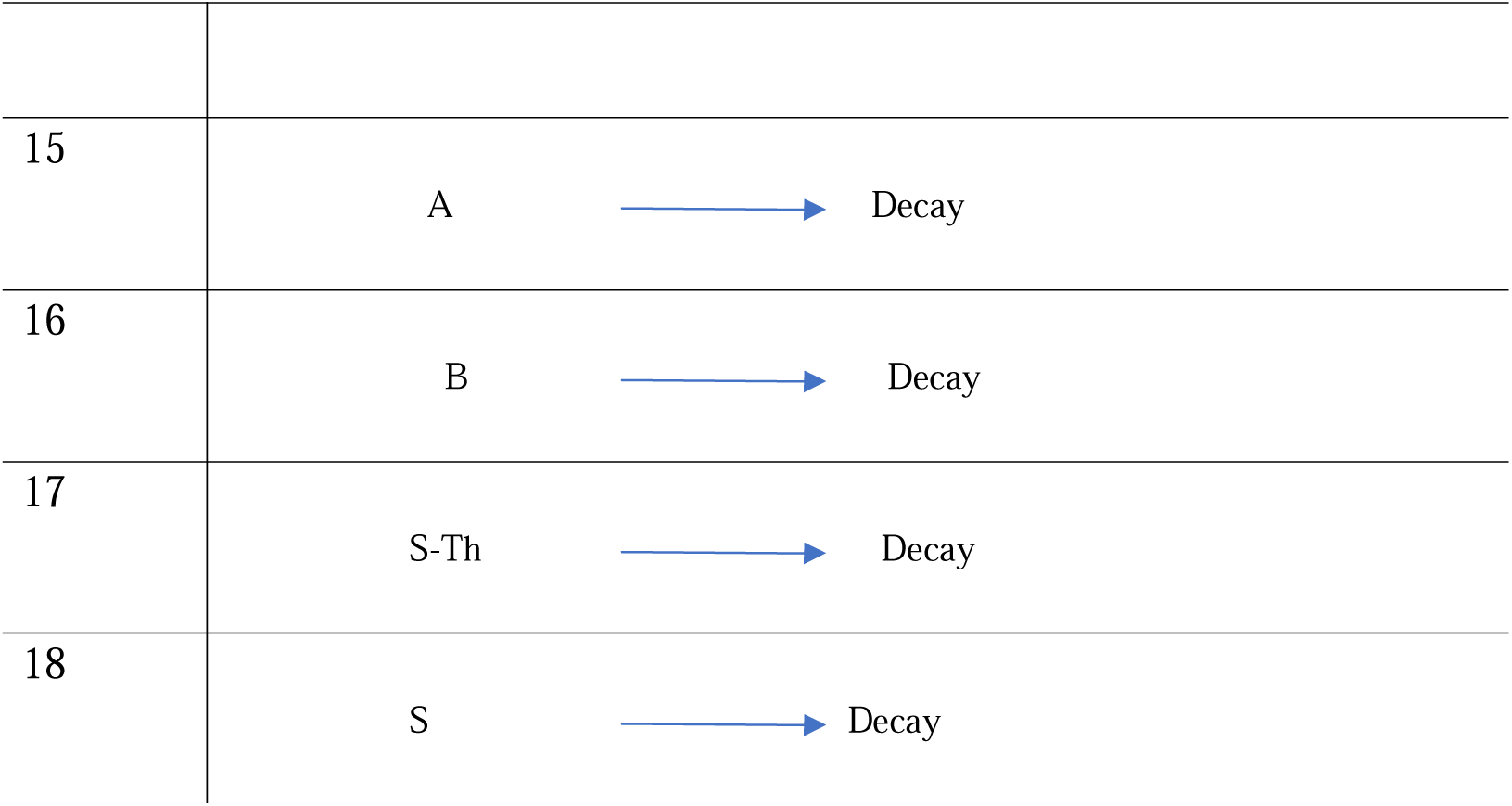
Reaction index and the name of each reaction in our cell-compartment model.

**Table 2:**
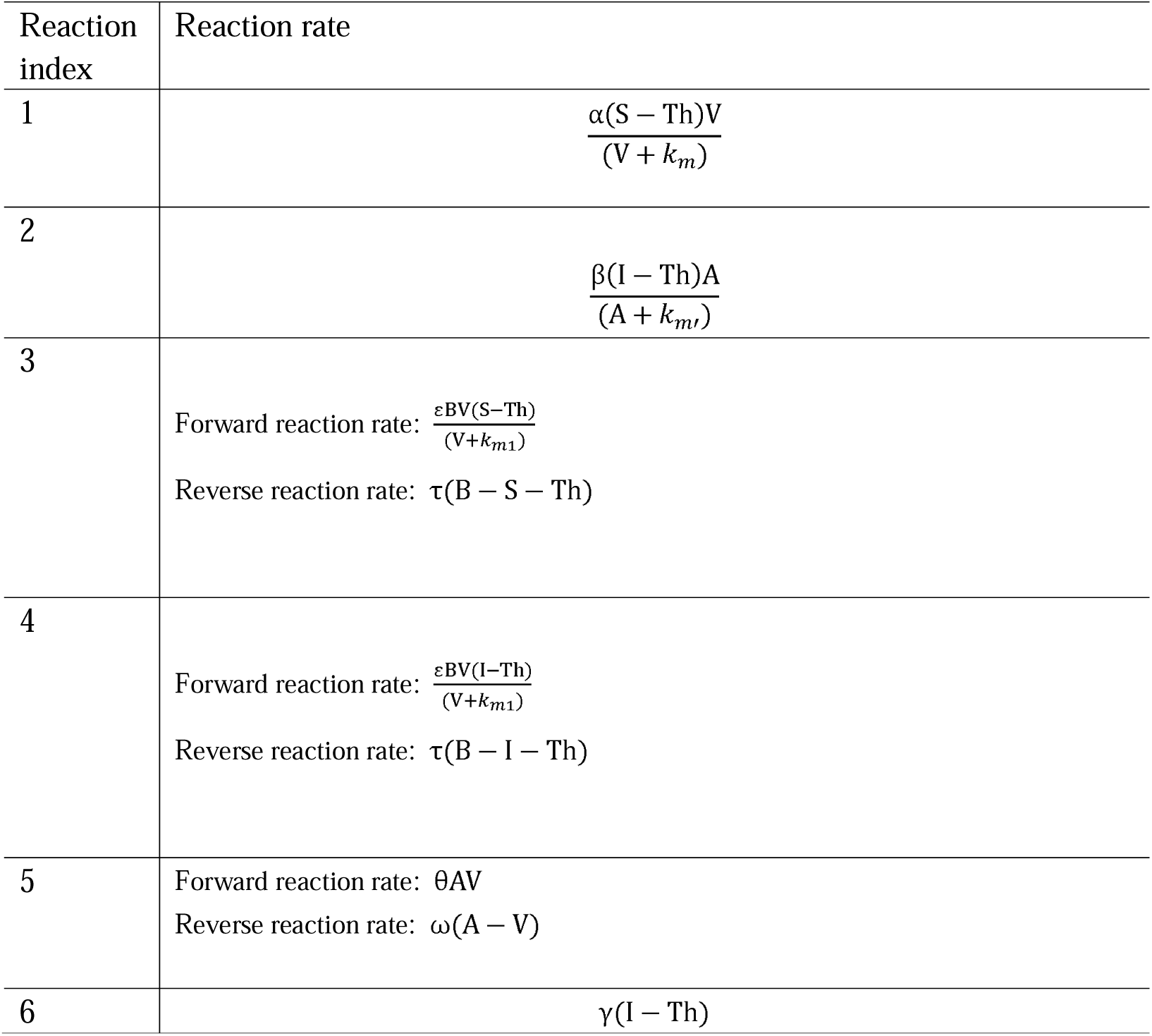
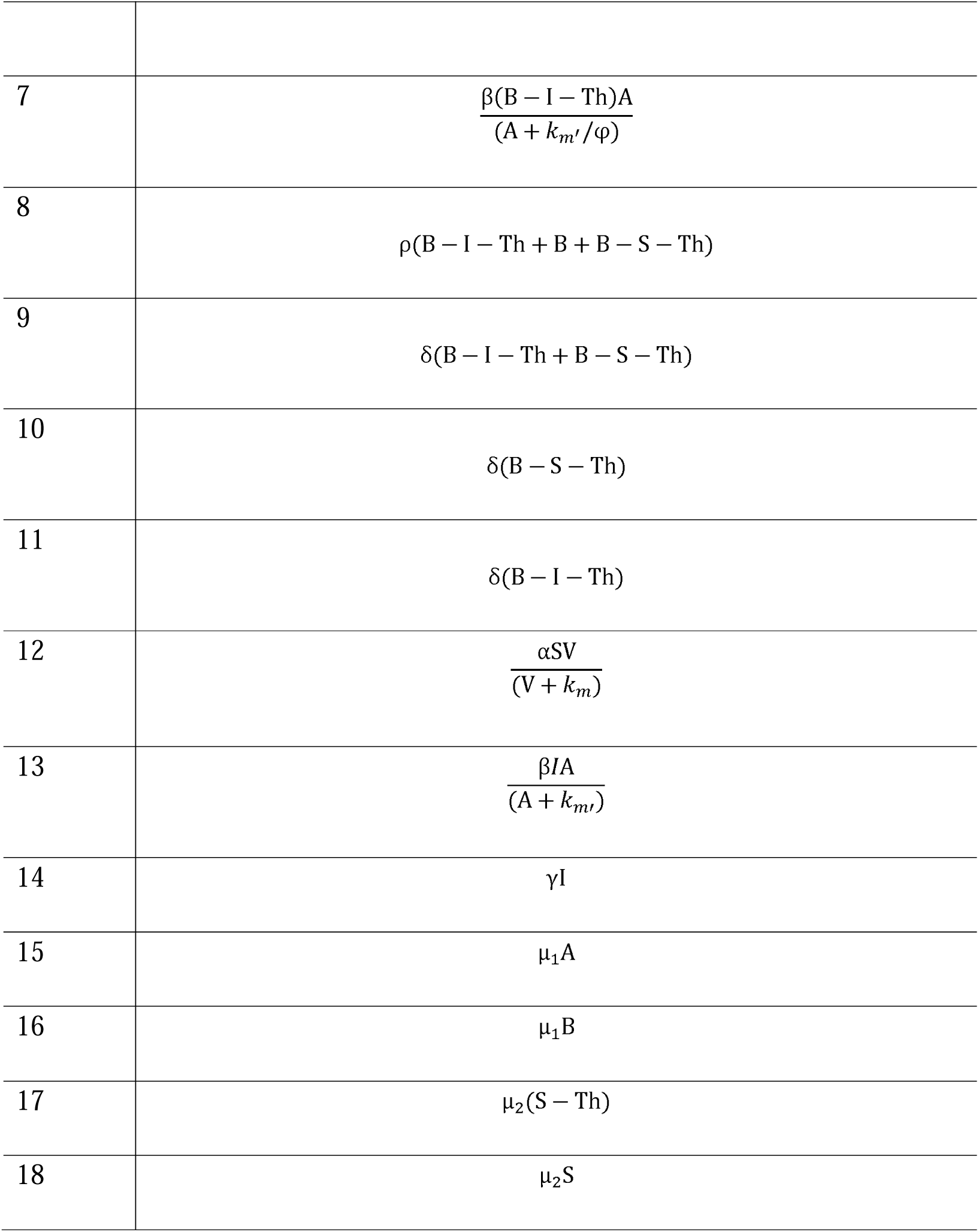
Reaction rate of each reaction in our cell-compartment model.

The fifth reaction is the binding reaction between antibody and virus, whose forward binding rate is θAV, and the reverse dissociation rate is ω(A-V). The sixth reaction represents the lysis of infected effector Th cells under the action of the virus, and its lysis rate is γ(I-Th). Assuming that each lysed cell contains θ virus particles, the rate of release of virus particles into the environment is θγ(I-Th). The seventh reaction is the ADCC clearance effect of germinal center effector Th cells infected with the virus, and its clearance rate is 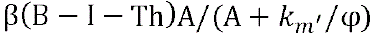, where φ is added because due to the spatial proximity effect, the concentration of antibodies faced by the core effector Th cells in the germinal center is often φ times the concentration of antibodies in the body fluid. Therefore, the ADCC effect of effector Th cells in the germinal center is often more significant and stronger. The eighth reaction is the process of antibody production, which is secreted by B cells, germinal center effector Th cells, and healthy effector Th cells and B cells forming germinal centers, with a production rate of 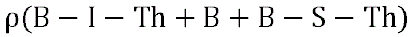. The ninth reaction is the regeneration process of B cells, and healthy effector Th cells and B cells forming germinal centers, with a production cells, which can be stimulated by germinal center effector Th cells and B cells forming germinal centers with infected effector Th cells and healthy effector Th cells, with a generation rate of 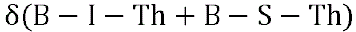. The tenth reaction represents the proliferation process of healthy effector Th cells, which can be stimulated by cytokines secreted by germinal centers formed by healthy effector Th cells and B cells, with a rate of δ(B – S-Th). The eleventh reaction represents the proliferation process of infected effector Th cells, which can be stimulated by cytokines secreted by germinal centers formed by infected effector Th cells and B cells, with a rate of δ(I – S – Th). The twelfth reaction is the infection process of susceptible cells, in HIV infection, it is the infection rate of other CD4+ cells. The thirteenth reaction is the ADCC clearance process of infected effector Th cells. The fourteenth reaction is the lysis process of infected effector Th cells under the action of the virus, with a rate of γI. Reaction 15 represents the natural degradation rate of antibody A, which is μ_1_A. Reaction 16 represents the natural degradation rate of B cells, which is μ_1_B, and to maintain the previous equilibrium state, we assume that B cells can be constantly replenished, with a replenishment rate of π_0_. Reaction 17 represents the natural degradation rate of effector Th cells, which is μ_2_S, with a self-supply constant of π_1_. Reaction 18 represents the natural degradation rate of other total Th cells, which is μ_1_(S-Th), with a self-supply constant of π_2_. Specific model description and parameter set can be referred to supplementary materials (model 2.1 to model 2.4).

Main reactions in this model are represented in Table 1.

The reaction rate is represented in Table 2.

The set of ordinary differential equations can be expressed as follows:

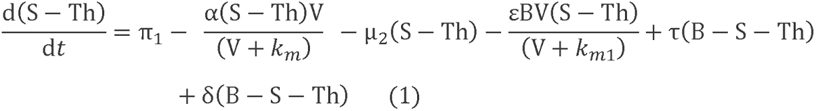

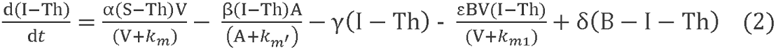

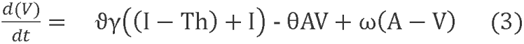

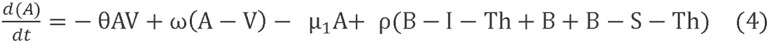

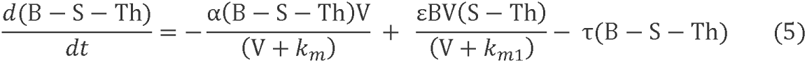

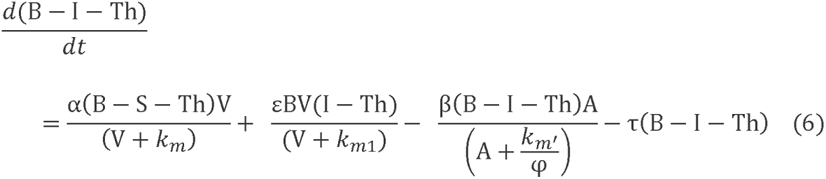

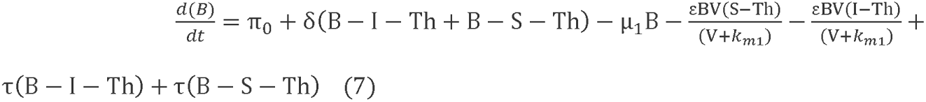

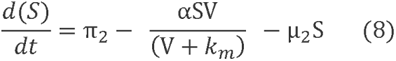

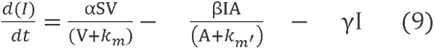

## 3 Results

### 3.1 The impact of T cell immunogenicity of antigens on immune responses

In our previous study [29], we analyzed how the T cell immunogenicity of antigens determines the fate of corresponding B cells, whether it leads to clonal deletion, massive proliferation, or the path towards clonal redemption.

This study further develops our previous model, from which we observe that antibody proliferation, or rather B cell proliferation, is mainly influenced by two factors: the rate of clearance of antigen-antibody complexes (k3) and the feedback regulation coefficient (k4) of antigen-antibody complexes on antibody regeneration. Generally, due to the presence of immune phagocytic cells such as natural killer (NK) cells in the body, antigen-antibody complexes are specifically recognized, leading to a clearance rate much higher than the decay rate of the antigen or antibody itself. The value of K4 depends on the T cell immunogenicity of the antigen molecule, that is, the primary sequence of the antigen molecule. After the antigen is phagocytosed and processed by antigen-presenting cells, it is degraded into short peptides and presented to the corresponding effector T cells. In viral infections, B cells mainly act as antigen-presenting cells, so effector T cells couple with B cells that secrete specific antibodies, forming germinal centers [28]. Stimulated effector T cells secrete cytokines to mediate their own division and proliferation, as well as the division and proliferation of B cells in germinal centers, thereby achieving rapid proliferation of high-affinity antibodies and screening out high-affinity antibodies within a short period of time. However, this process is premised on the antigenic substance having significant T cell immunogenicity. Otherwise, even if B cells can couple with effector T cells, the stimulation of these short peptides on effector T cells is not strong enough to induce sufficient secretion of cytokines and promote the proliferation of germinal centers. From the model’s perspective, when the value of k4 is significantly higher than that of k3, antigenic substances will lead to massive proliferation of specific antibodies. When k4 is smaller than k3, antigenic substances will induce rapid apoptosis of high-affinity antibodies, that is, clonal deletion. Specifically, as shown in Figure 3a, when exogenous pathogens invade (k4 = 1, k1 = 0.1, k3 = 0.4), high-affinity antibodies (k2 = 1e-6, represented by the red solid line) undergo rapid proliferation, while low-affinity antibodies (k2 = 0.5e-6, represented by the yellow solid line) have limited proliferation, and antibody proliferation effectively eliminates the virus (represented by the blue solid line). Through this mechanism, the immune system achieves enrichment and selection of specific neutralizing antibodies. In contrast, in Figure 3b, when self-antigens are present in the body (k4 = 0.2, k1 = 0, k3 = 0.4), due to the low T cell immunogenicity of self-antigens, the regenerative capacity of antibodies stimulated by germinal centers is significantly smaller than the clearance rate of antigen-antibody complexes. This leads to a sharp decrease in antibody quantity. Moreover, the stronger the affinity of the antibody, the faster its clearance rate, resulting in a drastic decline of high-affinity antibodies (k2 = 1e-6, represented by the blue solid line), while the decline of low-affinity antibodies (k2 = 0.5e-6, represented by the red solid line) is relatively slower, ultimately leading to clonal deletion. Figure 3 is generated using model 1.1. The detailed description and parameter set can be referred to supplementary materials (model 1.1).

**Figure 3a:**
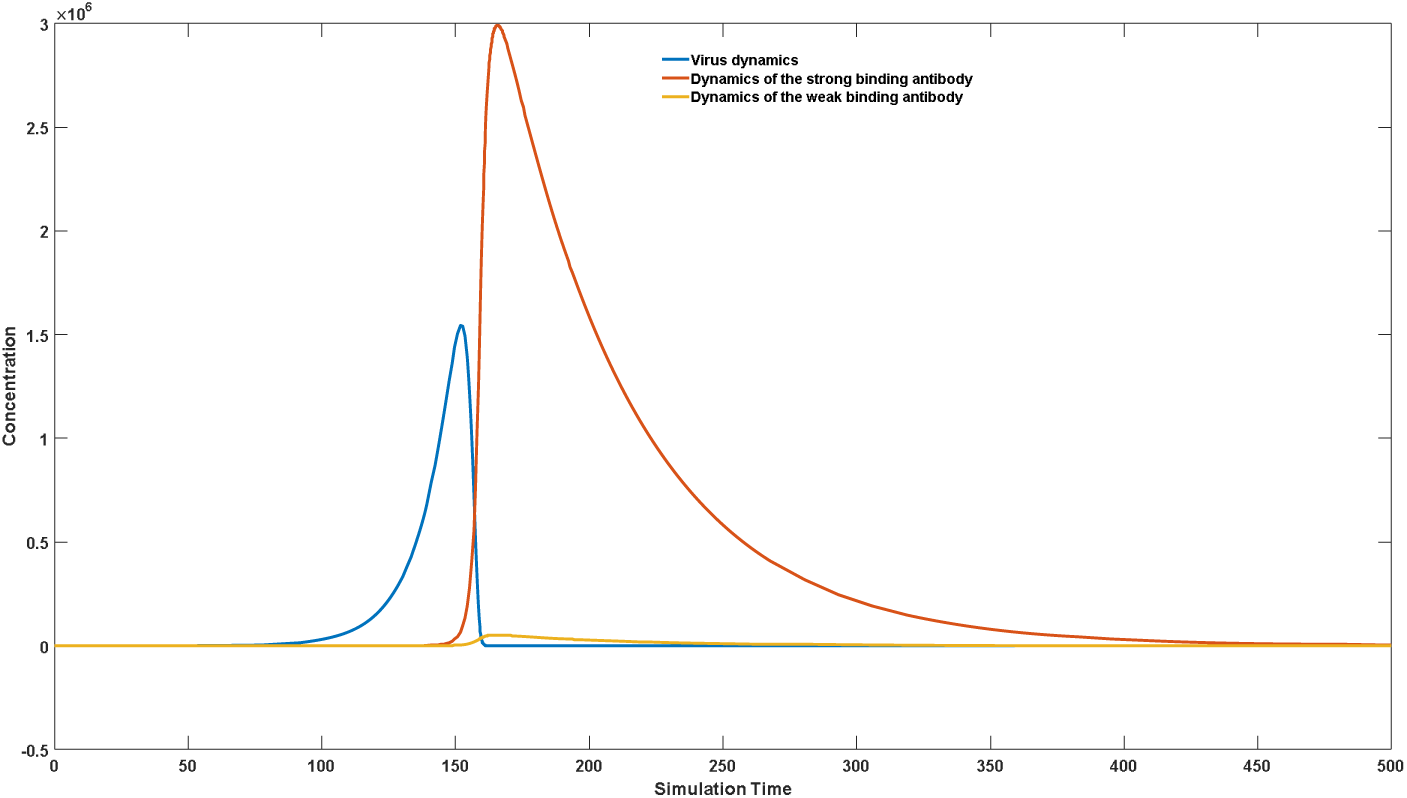
An illustration of how immune system screen for strong binding antibody against infectious diseases.

**Figure 3b:**
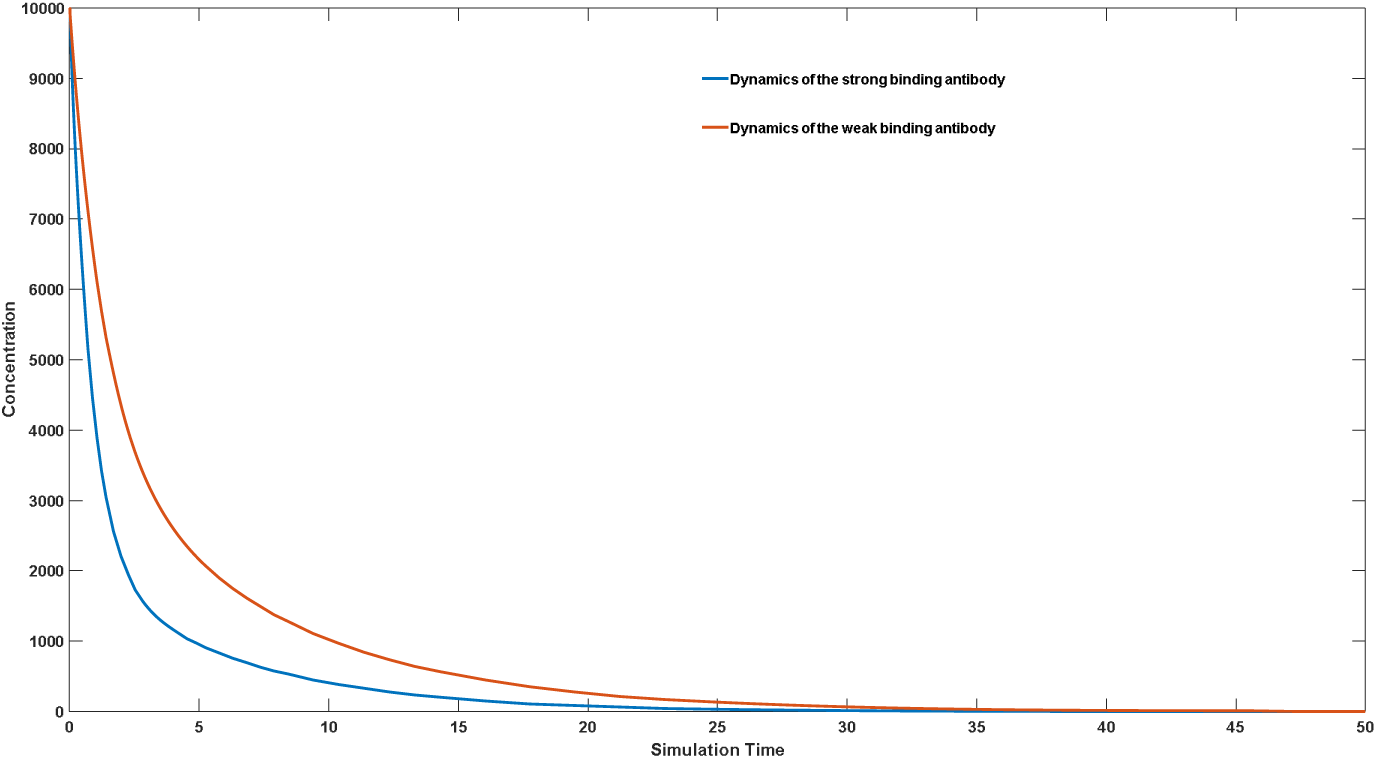
An illustration of clonal deletion

It is worth noting that not all self-antigenic substances have extremely low Th cell immunogenicity. Maintaining a low level of Th cell immunogenicity of self-antigens plays a critical role in the immune system’s ability to recognize “self” from “non-self.” The development of many autoimmune diseases is associated with increased Th cell immunogenicity. Nonetheless, over the course of evolution, we have not reduced or completely lost Th cell immunogenicity of self-antigens. We speculate that there are two reasons for this phenomenon. Firstly, the maintenance of Th cell immunogenicity of self-antigens is closely related to the maintenance of memory B cells, as explained in our earlier article [19]. In short, without the presence of self-antigens or with self-antigens having an extremely low level of Th cell immunogenicity, the level of memory B cells or IgG will exhibit an exponential decrease over time after losing pathogen stimulation. This leads to the loss of specific immune memory and frequent occurrence of secondary infections, which is detrimental to individual survival and evolution. Many experimental results also support this theory [30–32]. Secondly, maintaining a certain level of Th cell immunogenicity of self-antigens can effectively control the occurrence of cancer. Although some experiments have shown that the occurrence of cancer is related to mutations in protein sequences within the body [33–35], more evidence indicates that gene or protein mutations lead to abnormal metabolism and function of cells, affecting transcriptional and proteomic changes, which ultimately affect cell cycle regulation and lead to the occurrence of various types of cancer [36–38]. Therefore, for the vast majority of cancer cells, high levels of mutated proteins are not expressed. Rather, they express certain self-antigens that should not be highly expressed. Due to the loss of cell cycle regulation, this high expression becomes exponential. Maintaining a high level of Th cell immunogenicity of self-antigens can help the immune system better recognize and control the proliferation of cancer cells, as shown in Figure 4a (k3=0.4, k4=0.5). Under normal circumstances, self-antigenic substances do not proliferate, with a self-proliferation coefficient of k1=0. Therefore, self-antigenic substances (blue solid line) are maintained at a stable low level under the action of antibodies (red solid line). When cancer occurs, the self-proliferation coefficient changes to k1=0.1. However, due to the maintenance of relatively high Th cell immunogenicity of self-antigens, this self-antigenic substance (blue dashed line) fluctuates at a low level under the action of antibodies (red dashed line), allowing the immune system to better recognize and control the proliferation of cancer cells. However, when the Th cell immunogenicity of self-antigens decreases, the situation may change, as shown in Figure 4b. When k4 decreases to 0.46, although self-antigenic substances are controllable as a whole, they exhibit large fluctuations and high peak concentrations, leading to a significant decrease in the immune system’s ability to detect and kill cancer cells. When the Th cell immunogenicity of self-antigens further decreases, the situation may become uncontrollable, as shown in Figure 4c. When k4 further decreases to 0.42, the proliferation of self-antigenic substances becomes uncontrollable, indicating that the immune system has completely lost the ability to inhibit the early development of cancer. As age increases, the activity of Th cells significantly decreases, which may cause the Th cell stimulation ability of self-antigens to decrease, thereby increasing the risk of cancer. It also results in a decrease in the Th cell immunogenicity of exogenous antigens, which increases the risk of death from infectious diseases. In recent years, numerous experiments have shown that antibodies and B cells are directly involved in cancer immunity [39–42], which to a certain extent validates our simulation results. Thus, maintaining the sensitivity of the immune system may become an important factor in controlling the onset of cancer. Figure 4 is generated using model 1.2. The detailed description and parameter set can be referred in supplementary materials (model 1.2).

**Figure 4a:**
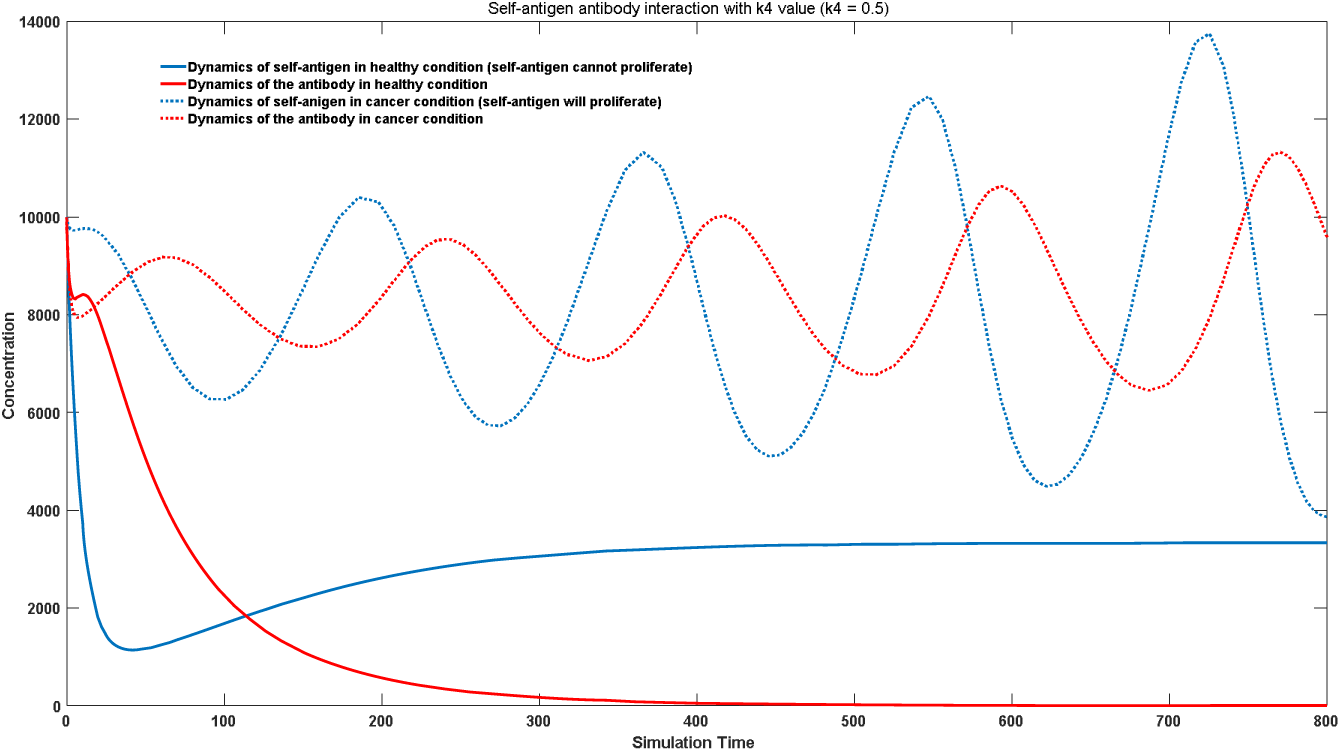
An illustration of how immune system prevent cancer well with strong Th cell immunogenicity. (strong feedback signal)

**Figure 4b:**
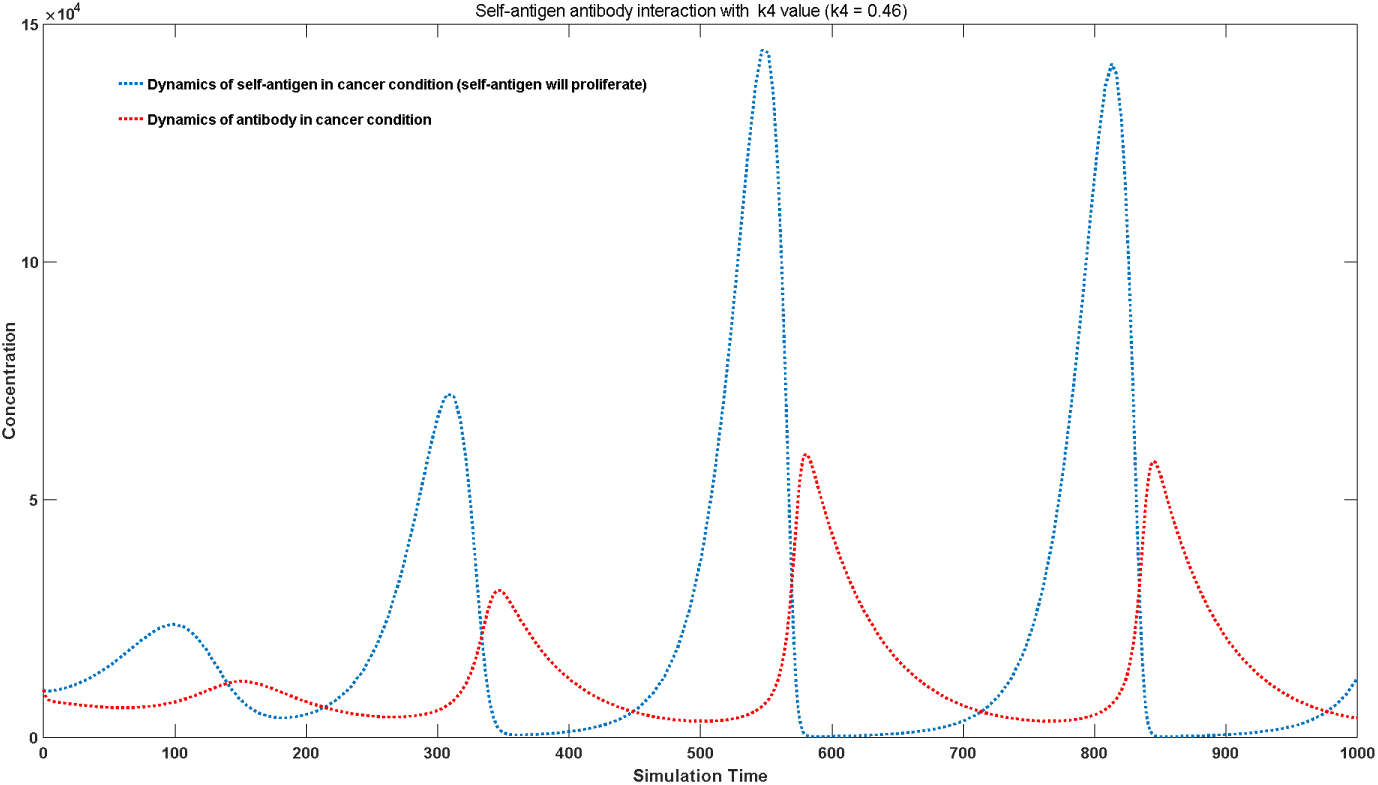
Impaired Th Cell Immunogenicity Leads to Decreased Anti-Cancer Immune Control. (decreased feedback signal)

**Figure 4c:**
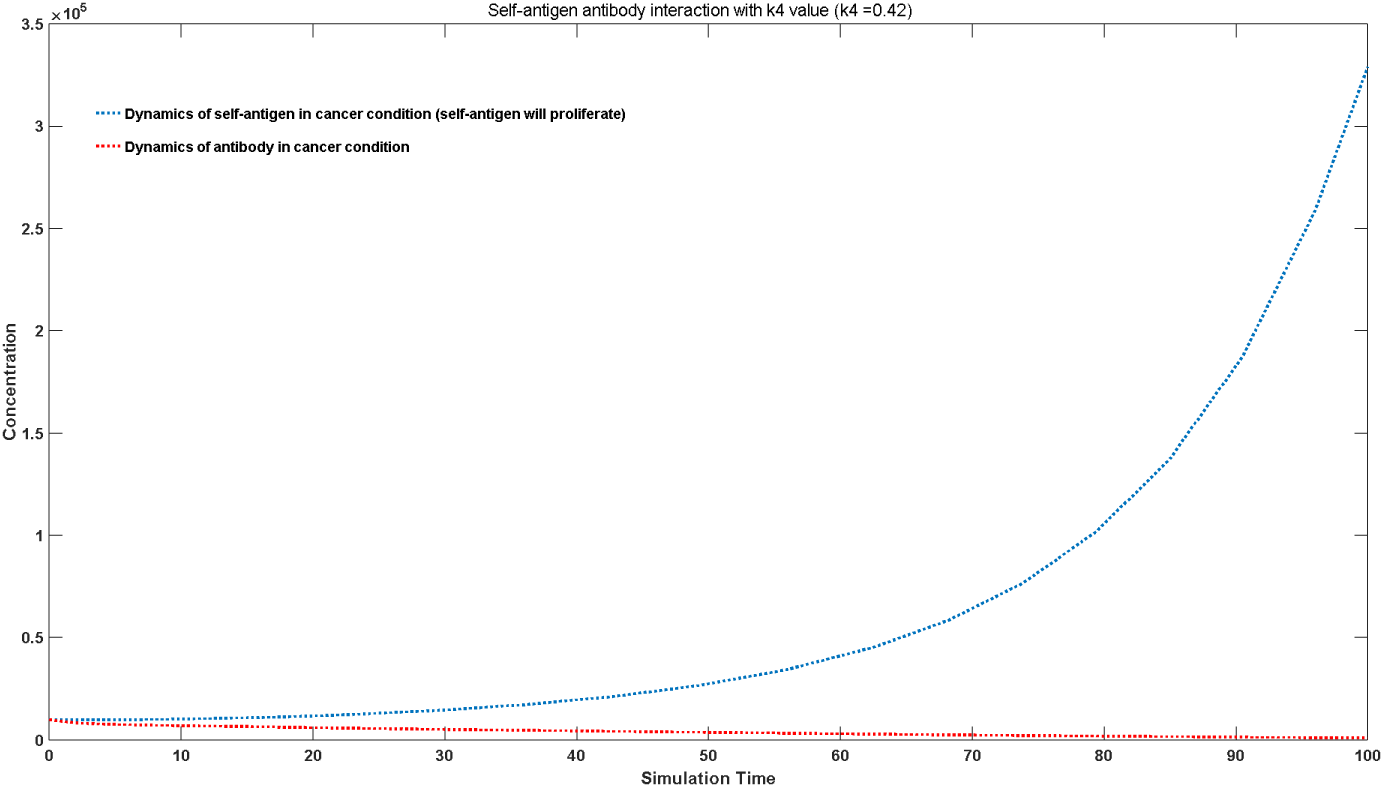
Low Th Cell Immunogenicity Leads to the complete failure of Anti-Cancer Immune Control.

### 3.2 The B cells-targeted ADCC effect

Through the dynamic data analysis of the virus-antibody interaction in different dengue fever patients, we have discovered an intriguing phenomenon. Despite having nearly identical peak antibody concentrations, there are significant differences in peak viral loads among these patients, which also correspond to noticeable discrepancies in symptomatology [43]. To investigate which parameter contributes to this phenomenon, we conducted a sensitivity analysis of various parameters and found that the variation in the clearance rate of antigen-antibody complexes, represented by the k3 value, is responsible for this occurrence. The clearance of antigen-antibody complexes involves the activity of NK cells. Additionally, we propose the hypothesis that different antibodies or antigenic substances may lead to variations in the clearance rate of these complexes, primarily through differential ADCC effects on B cells. As depicted in Figure 5, when a monomeric antigen molecule binds to a membrane-bound antibody on the surface of B cells, the antigenic determinant cluster is occupied, preventing binding with free antibodies. Consequently, the Fc epitope of the antibody remains hidden, thus failing to initiate ADCC-mediated B cell death. However, when the antigen molecule is a homodimer or multimer, or in the case of symmetrical viruses, the phenomenon of free antibody binding to non-B cell binding sites on the antigen occurs. This results in the extensive formation of Fc antibody epitopes on the surface of B cells, inducing ADCC-mediated B cell death, as manifested by an increased clearance rate of antigen-antibody complexes (k3) in our model.

**Figure 5:**
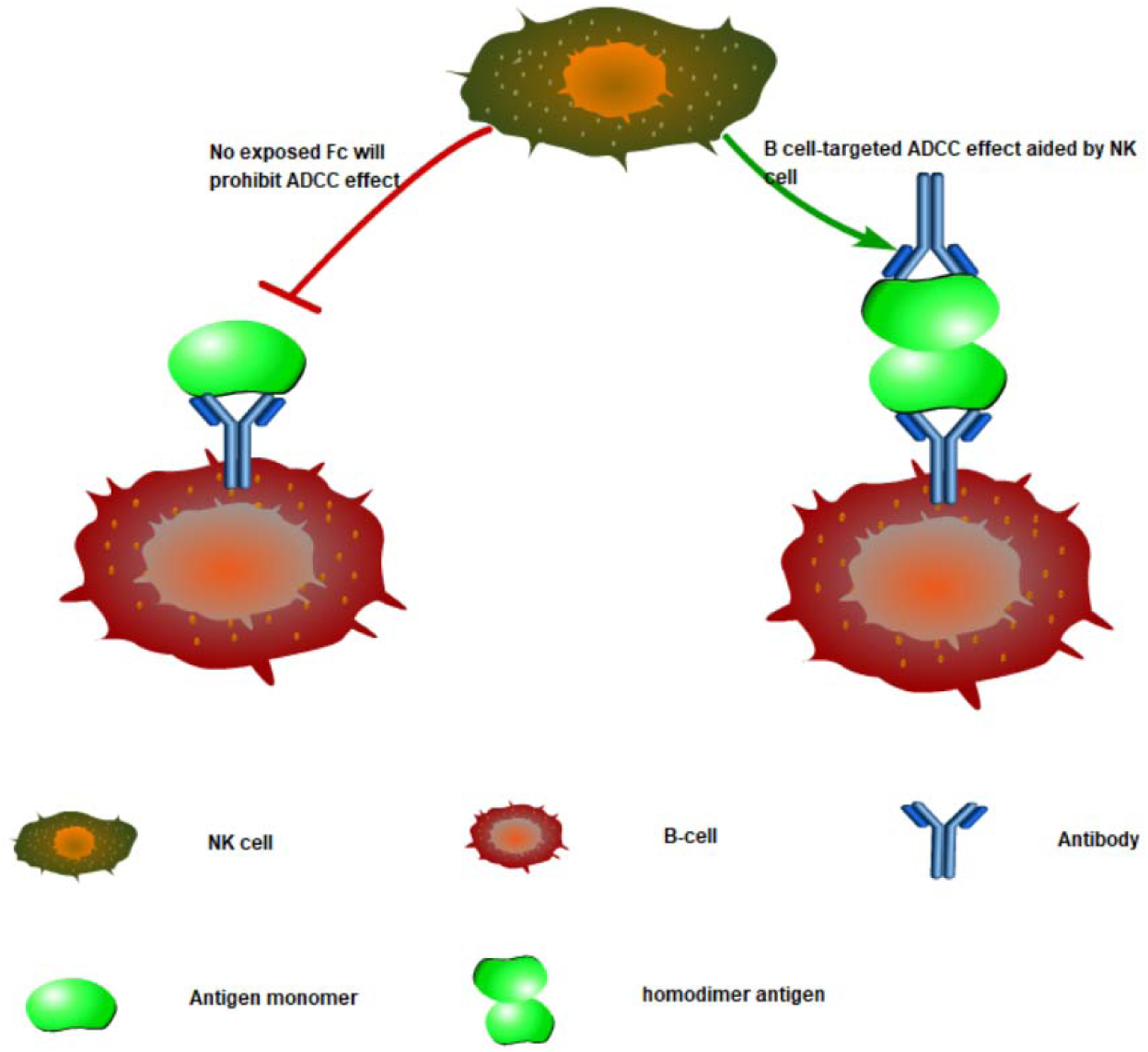
Proposed mechanism of B-cell targeted ADCC effect

As shown in Figure 6, the presence of ADCC effects on B cells significantly elevates the peak concentration of the virus (solid purple line) compared to the absence of ADCC effects (dotted purple line). Simultaneously, the concentration of antigen-antibody complexes (solid blue line) is considerably higher when ADCC effects are present (compared to the absence of ADCC effects, dotted blue line). Although the antibody concentration is lower in the presence of ADCC effects than in the control group without ADCC effects, this decrease is not as substantial as the reduction in viral load. This partially explains why viruses tend to evolve into symmetrical structures such as spherical or polyhedral particles, as these symmetrical structures facilitate ADCC effects on B cells and thereby maintain higher viral loads during infection, leading to enhanced transmission capabilities. Understanding the ADCC effects on B cells not only elucidates the significant differences in peak viral loads among patients during viral infections and the consequent variations in the risk of severe illness but also contributes to the development of macromolecular drugs for autoimmune diseases. Numerous studies in materials science have demonstrated that for chimeric nanomaterials with a linear framework, bivalent or low-valency antigens often exhibit the ability to eliminate self-reactive B cells or antibodies, whereas high-valency antigens usually exhibit less efficacy [44–47]. We hypothesize that this is because both high-valency and bivalent antigens can trigger the same ADCC effects on B cells, but high-valency antigens induce a more robust Th-cell immunogenicity due to the higher peptide fragment concentrations presented during antigen presentation, resulting in an increase in the k4 value. Therefore, the use of bivalent antigens yields the best results. Moreover, employing molecular biology techniques to design low Th-cell immunogenicity homodimers may potentially achieve superior therapeutic effects. This B cell-mediated ADCC effect may originate from the complement system or the cytotoxicity of NK cells. Currently, abundant research supports the dominant role of NK cells in the ADCC-mediated killing of B cells [48–50]. Figure 6 is generated using model 1.3. The detailed description and parameter set can be referred in supplementary materials (model 1.3).

**Figure 6:**
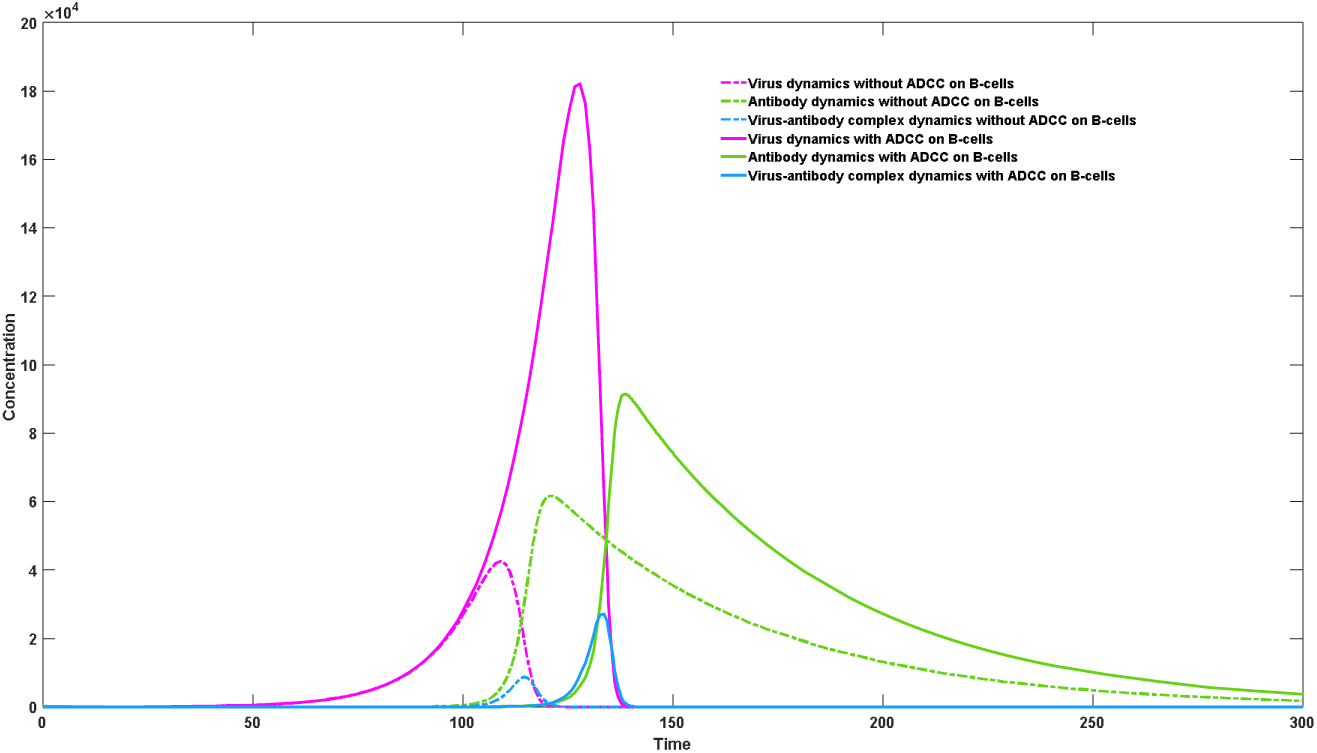
The impact of B cell-targeted ADCC on virus-antibody dynamics

### 3.3 ADCC Effect Targeting Th Cells

The ADCC effect targeting Th cells can influence the survival of Th cells in the germinal center, thereby affecting antibody production. The binding of antigens to Th cells can also lead to the destruction of the germinal center and hinder antibody generation. In 2017, Bogers et al. reported that the use of HIV antigen gp140, which had its CD4 binding domain removed, could rapidly stimulate the generation of specific antibodies in monkeys [51]. They suggested that the binding of gp140 to CD4 molecules could affect the activity of Th cells in the germinal center, resulting in reduced antibody production. Although this viewpoint has some validity, we cannot rule out the possibility that the observed phenomenon is due to ADCC-induced apoptosis of Th cells in the germinal center. This ADCC effect is likely mediated to a large extent by the complement system, as NK cells have limited access to Th cells at the core of the germinal center. Simulation results from a simple model tend to support the theory that Th cell apoptosis through ADCC occurs. Although both theories can hinder antibody production, the difference lies in the fact that antigen-mediated functional impairment of Th cells will proportionally reduce the rate of production for all types of antibodies, while the ADCC apoptosis theory of Th cells will pose greater obstacles to the production of high-affinity antibodies and lesser obstacles to the production of low-affinity antibodies. This leads to the observation that even after long-term chronic infection, most individuals are unable to effectively stimulate specific neutralizing antibodies. This phenomenon is widespread in HIV infections, with only approximately 10%-25% of infected individuals able to produce high-affinity specific neutralizing antibodies [52]. This suggests that even in the presence of antigen molecules that functionally suppress Th cells, the Th cell-targeted ADCC effect still plays a significant role. From Figure 7, it can be observed that in the absence of Th cell-mediated ADCC effects, high-affinity antibodies (solid red line, k2 = 1e-5) proliferate faster than low-affinity antibodies (solid green line, k2 = 9e-6), and the viral concentration (solid blue line) is better controlled compared to when ADCC effects are present (dashed blue line). Most importantly, in the absence of Th cell-mediated ADCC effects, the body can achieve complete clearance of the virus, as indicated by the absence of a secondary increase in viral concentration (solid blue line). In contrast, when Th cell-mediated ADCC effects are present, high-affinity antibodies (dashed red line) do not exhibit a more significant proliferative advantage compared to low-affinity antibodies (dashed green line). Although both show a significant decline compared to the control group, the decline in high-affinity antibodies is more pronounced, making it difficult for high-affinity antibodies to be selected by the immune system based on proliferative advantage. At the same time, this situation hinders viral clearance and leads to the occurrence of long-term chronic infections, with the virus undergoing secondary or repeated proliferation (dashed blue line). Figure 7 is generated using model 1.4. The detailed description and parameter set can be referred in supplementary materials (model 1.4).

**Figure 7:**
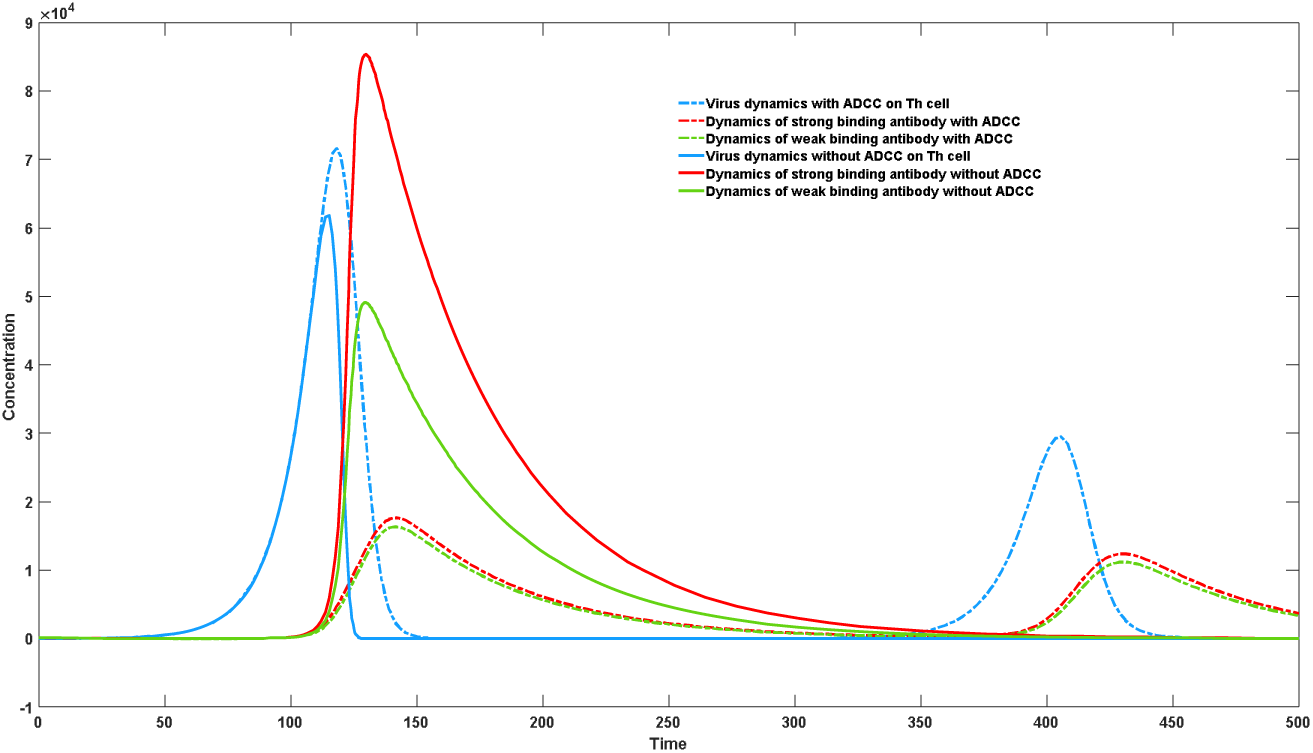
The impact of Th cell-targeted ADCC on virus-antibody dynamics

### 3.4 Quantitative Study of HIV Infection Process using Complex Models

After utilizing a simple coarse-graining model to investigate the ADCC effect of B cells and Th cells, we attempted to employ a more sophisticated model for a more precise simulation of the HIV infection process. The basic framework of this complex model is described in the Methods section and provides a better understanding of the consequences resulting from the ADCC effect of Th cells. With a more detailed characterization of cell types, this model allows us not only to observe the dynamic changes in virus and antibody kinetics but also to monitor and describe the population changes in various major cell types, including B cells, effector Th cells, total Th cells, infected effector Th cells, and total infected Th cells. By integrating viral changes with the dynamics of total infected Th cells, we can comprehensively assess the severity of the infection in patients. A detailed description and parameter settings for the dual-antibody competition model are provided in the supplementary materials. Figure 8 illustrates the characteristics of different compartments during infection in the presence of effector Th cells, with an initial total susceptible cell count of 10^8^ and an initial total effector Th cell count of 10^6^. Figure 9 depicts the infection in the absence of effector Th cells. In Figure 8a, it can be observed that in the presence of effector Th cells during infection, the total count of effector Th cells proliferates, but a majority of them become infected (indicated by the solid red line), while the remaining uninfected effector Th cells decrease to a smaller count from the initial state (indicated by the solid blue line). Conversely, in the absence of effector Th cells during infection (Figure 9a), the total count of effector Th cells proliferates (indicated by the solid blue line), and since the total infected susceptible cells do not include effector Th cells, there are no infected Th cells (indicated by the solid red line). In Figure 8b, it can be observed that in the presence of effector Th cells during infection, the host-virus interaction does not completely eliminate the virus. The viral count (indicated by the solid blue line) initially proliferates significantly, then decreases to a certain level and stabilizes, resulting in a long-term chronic infection scenario. Conversely, in the absence of effector Th cells during infection (Figure 9b), the host-virus interaction leads to substantial generation of antibodies, effectively eliminating the virus (indicated by the solid blue line), thus preventing the occurrence of long-term chronic infection. In Figure 8c, it can be observed that in the presence of effector Th cells during infection, the production of high-affinity antibodies (affinity = 1e-5) is suppressed due to the ADCC effect of Th cells (indicated by the solid blue line), with the production rate even lower than that of low-affinity antibodies (affinity = 1e-6) (indicated by the solid red line). The host is unable to achieve complete clearance of the virus, nor can it generate a substantial amount of antibodies or establish a proliferation advantage for high-affinity antibodies. Conversely, in the absence of effector Th cells during infection (Figure 9c), high-affinity antibodies (indicated by the solid blue line) experience significant proliferation and demonstrate a proliferation advantage over low-affinity antibodies (indicated by the solid red line). In Figure 8d, it can be observed that in the presence of effector Th cells during infection, the host-virus interaction does not completely eliminate infected cells, causing a decline in healthy cells to a lower level (indicated by the solid blue line), while infected cells can be maintained at a higher level in the long term (indicated by the solid red line). Conversely, in the absence of effector Th cells during infection (Figure 9d), the host-virus interaction can eliminate all infected cells (indicated by the solid red line), and healthy cells gradually recover to normal levels after complete viral clearance (indicated by the solid blue line). Figure 8 and Figure 9 are generated using model 2.1, The detailed description and parameter set can be referred in supplementary materials (model 2.1).

**Figure 8a:**
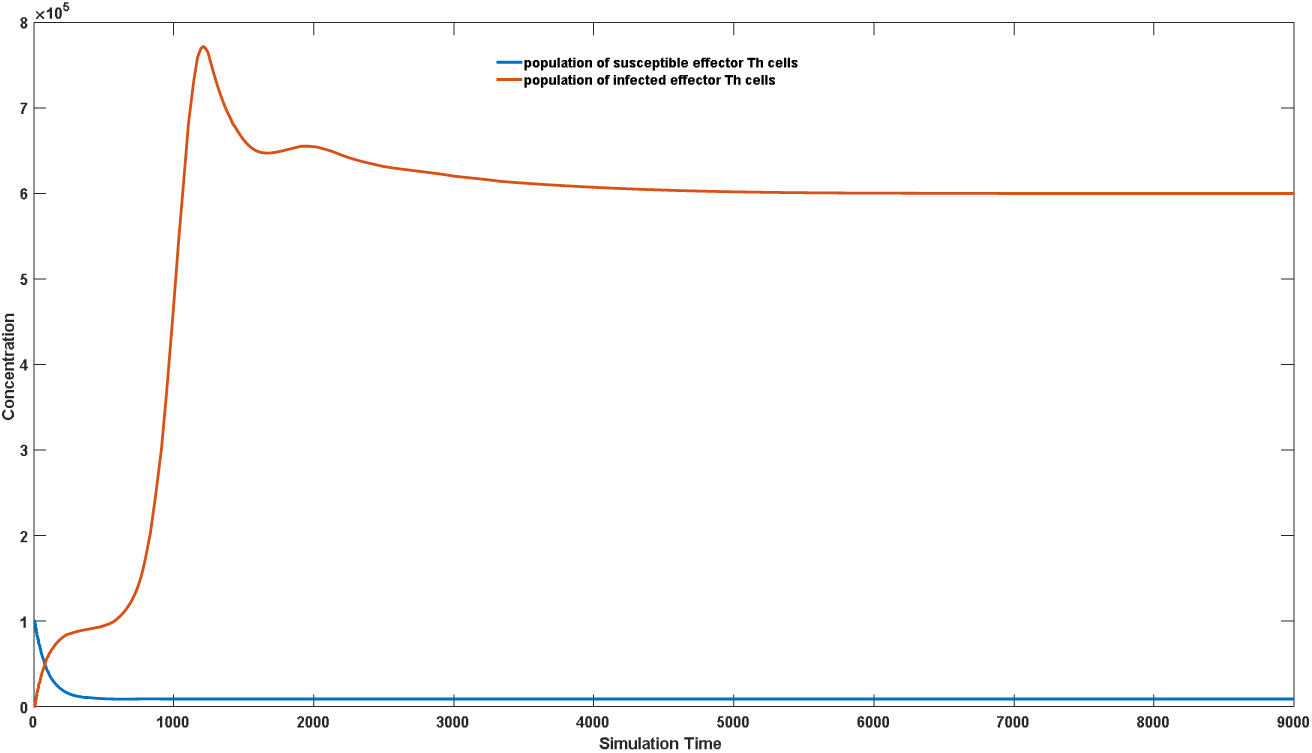
Dynamics of susceptible and infected effector Th cell considering Th-cell targeted ADCC effect.

**Figure 8b:**
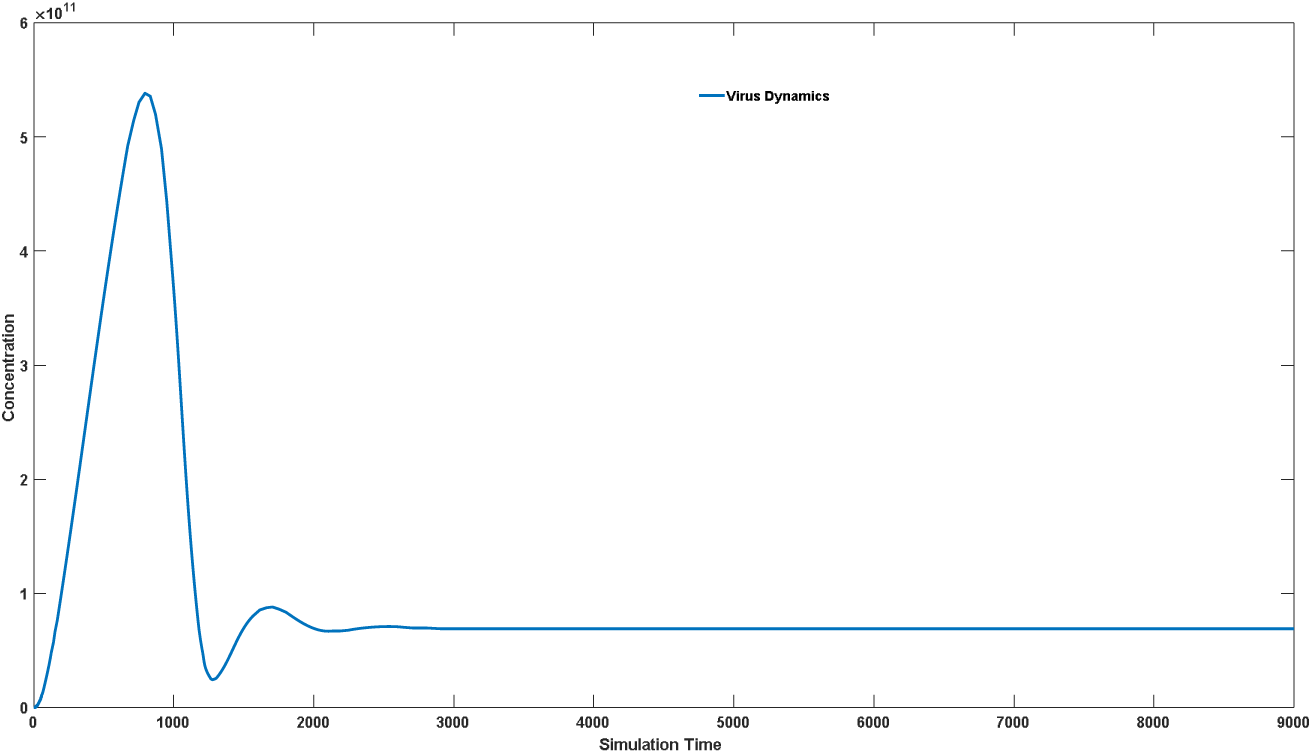
Dynamics of virus considering Th-cell targeted ADCC effect.

**Figure 8c:**
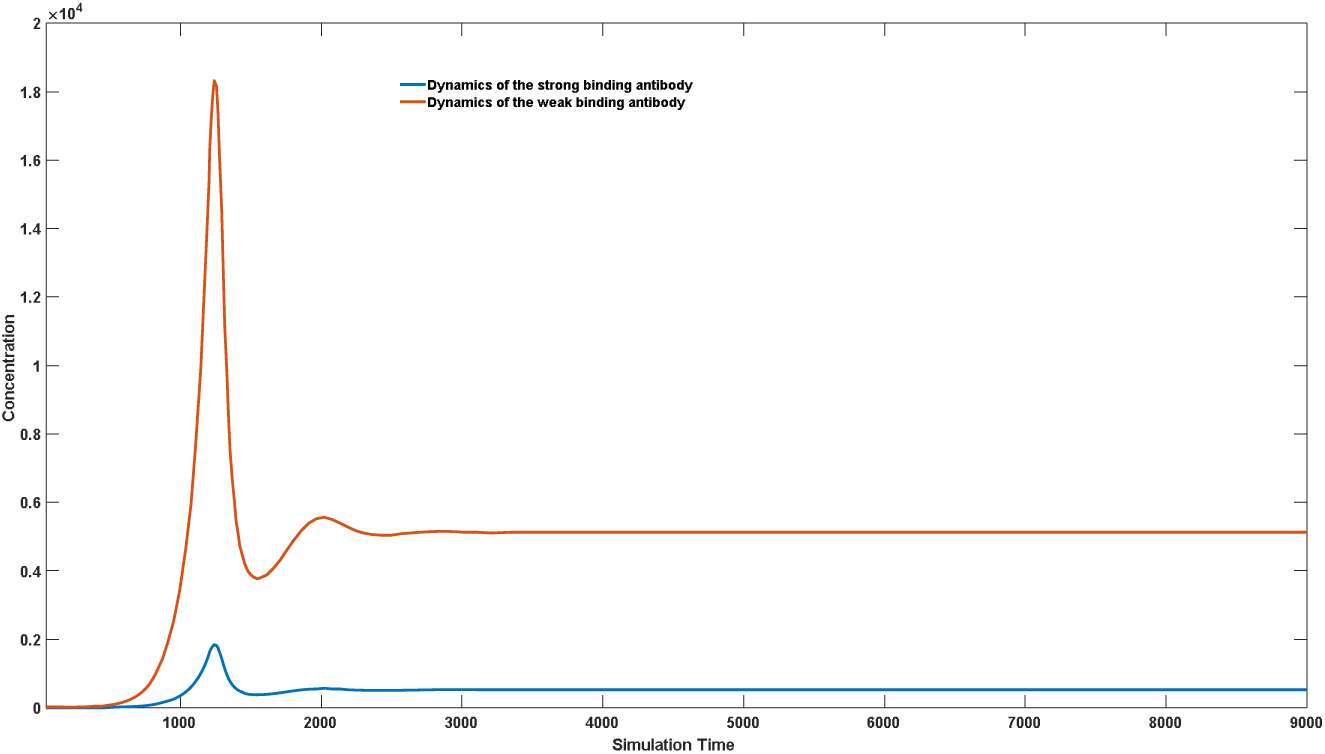
Dynamics of antibodies with different binding affinity considering Th-cell targeted ADCC effect.

**Figure 8d:**
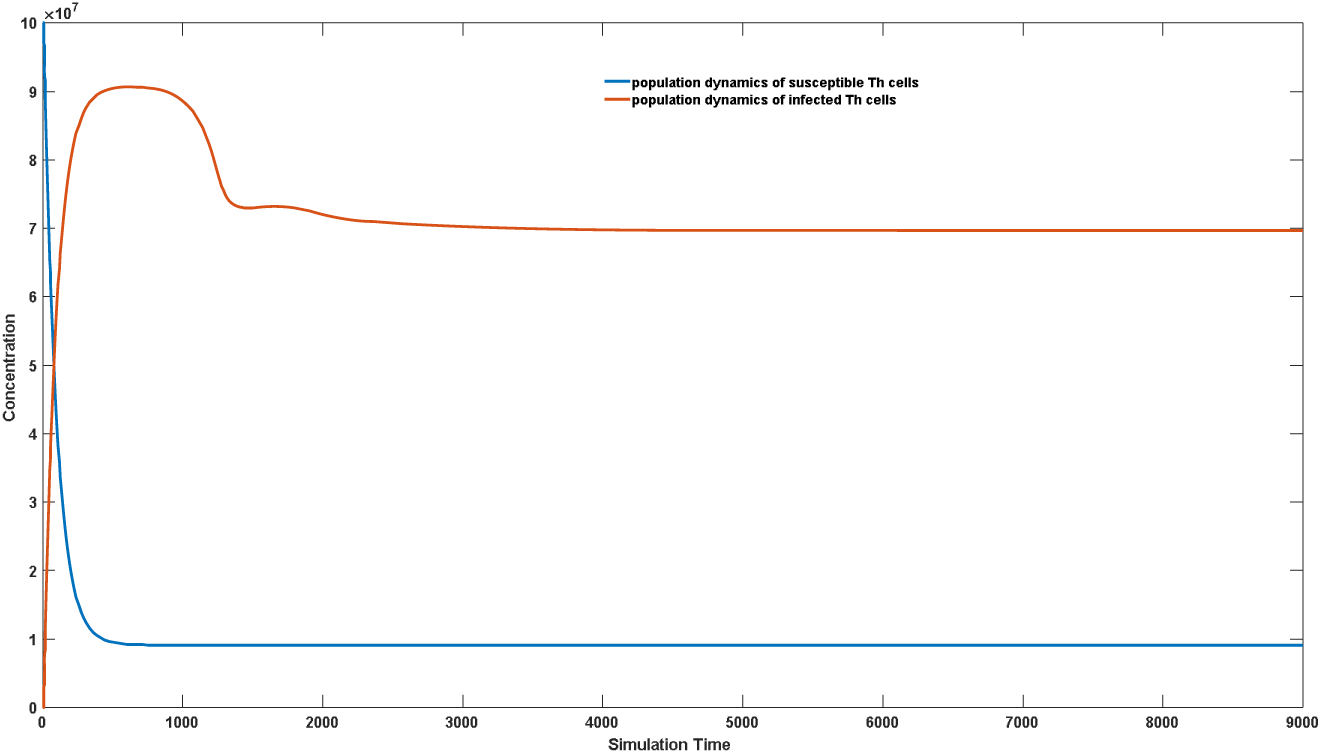
Dynamics of overall susceptible and infected CD4+ cells considering Th-cell targeted ADCC effect.

**Figure 9a:**
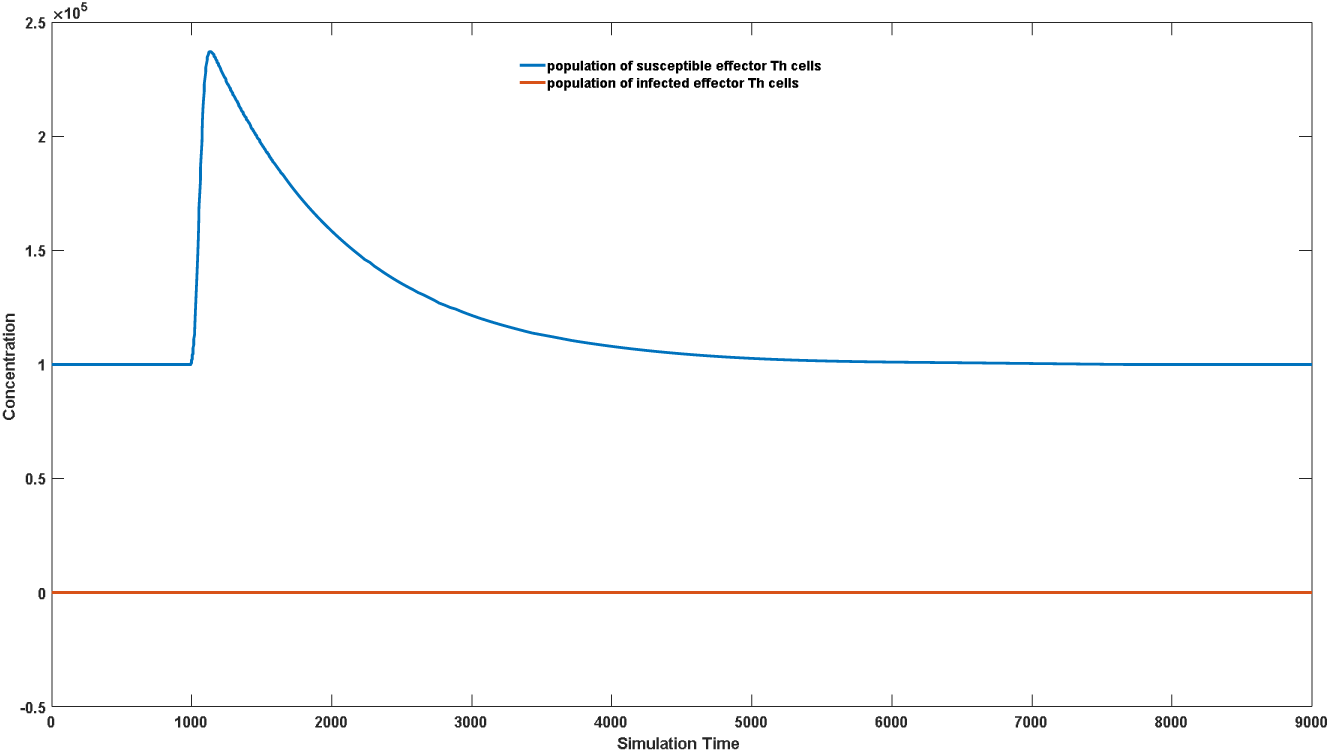
Dynamics of susceptible and infected effector Th cell without Th-cell targeted ADCC effect.

**Figure 9b:**
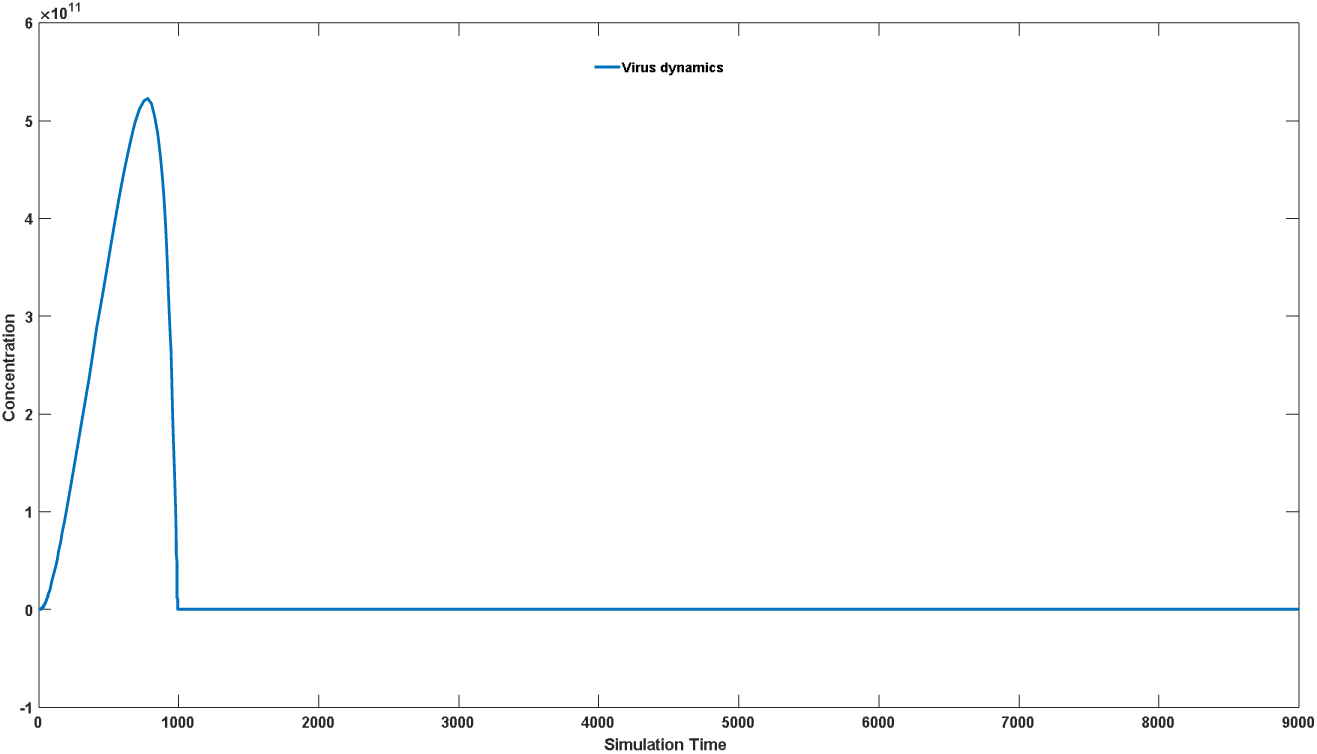
Dynamics of virus without Th-cell targeted ADCC effect.

**Figure 9c:**
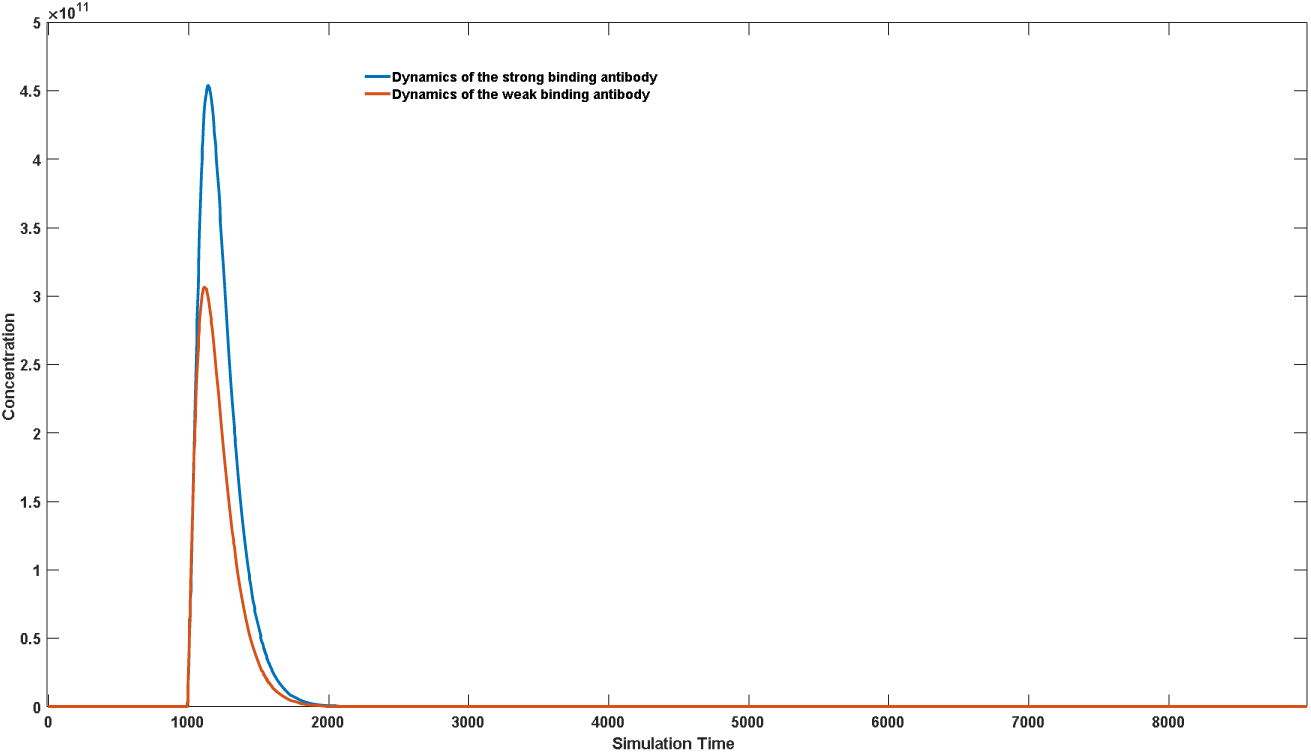
Dynamics of antibodies with different binding affinity without Th-cell targeted ADCC effect.

**Figure 9d:**
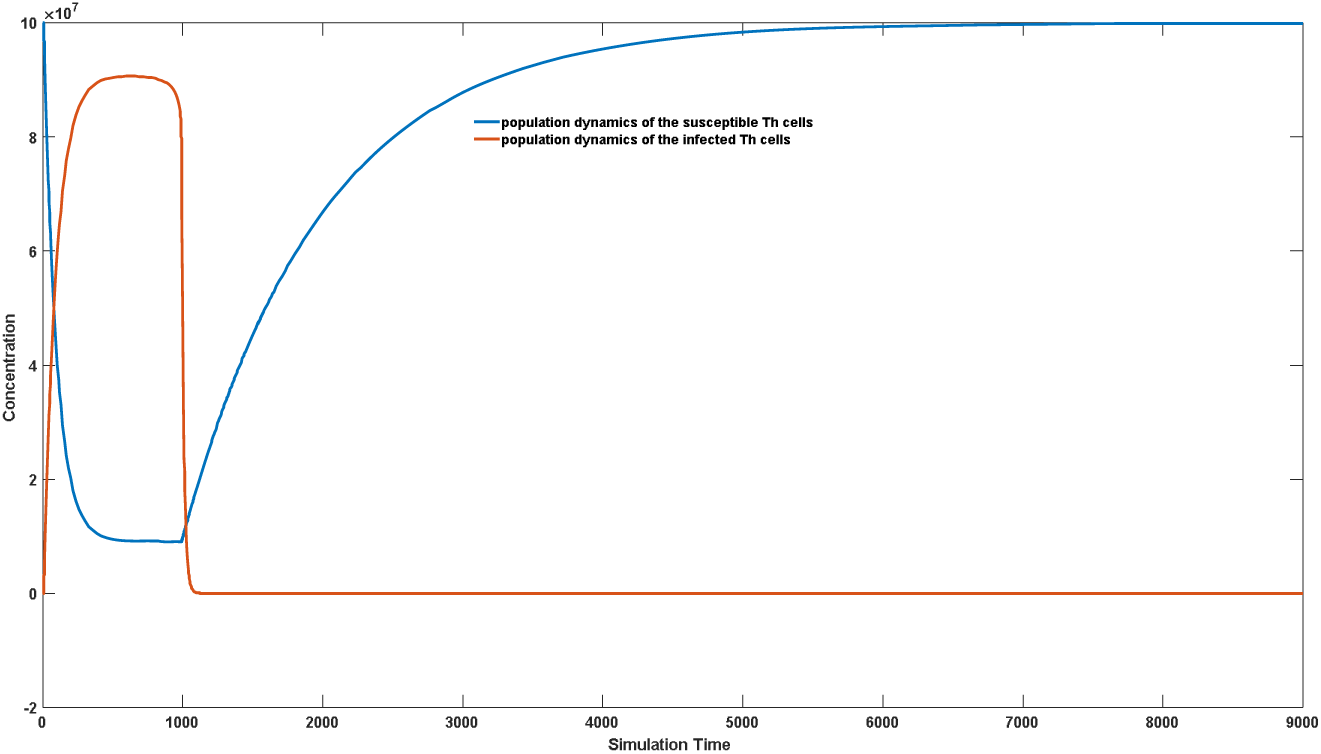
Dynamics of overall susceptible and infected CD4+ cells without Th-cell targeted ADCC effect.

### 3.5 Specific Neutralizing Antibodies are More Effective than Non-neutralizing Antibodies

We further employed sophisticated models to investigate why specific neutralizing antibodies exhibit better viral inhibition than conventional antibodies. For HIV infection, it has been recognized that not all high-affinity antibodies contribute to effective protection [53–55], despite their equal ability to bind to the virus and induce complex clearance. This is because specific neutralizing antibodies not only induce immune system clearance of the virus but also specifically bind to target cell recognition sites, thereby blocking further viral invasion of healthy cells. Whether it is HIV infection or other viral infections such as COVID-19, neutralizing antibodies targeting surface binding proteins often exhibit superior viral inhibitory effects compared to high-affinity antibodies targeting other sites. The specific models and parameter settings are provided in the supplementary materials. As shown in Figure 10a, if non-specific neutralizing antibodies are induced in the host, the virus is not well controlled. After the initial peak of infection, the virus experiences a decline, followed by secondary reinfection and maintains a certain concentration level (solid blue line). Meanwhile, these non-specific neutralizing antibodies are also induced but gradually decrease as the initial virus level declines, maintaining a very low level during subsequent chronic infection stages (solid red line). In contrast, if specific neutralizing antibodies are induced in the host, the virus is well controlled. After the initial peak of infection, the virus declines and remains at an extremely low level due to the presence of specific neutralizing antibodies (solid blue line). Similar to non-specific neutralizing antibodies, the levels of specific neutralizing antibodies decrease as the initial virus level declines and remain at a lower level during subsequent stages (solid red line). Figure 10 is generated using model 2.2. The detailed description and parameter set can be referred in supplementary materials (model 2.2).

**Figure 10a:**
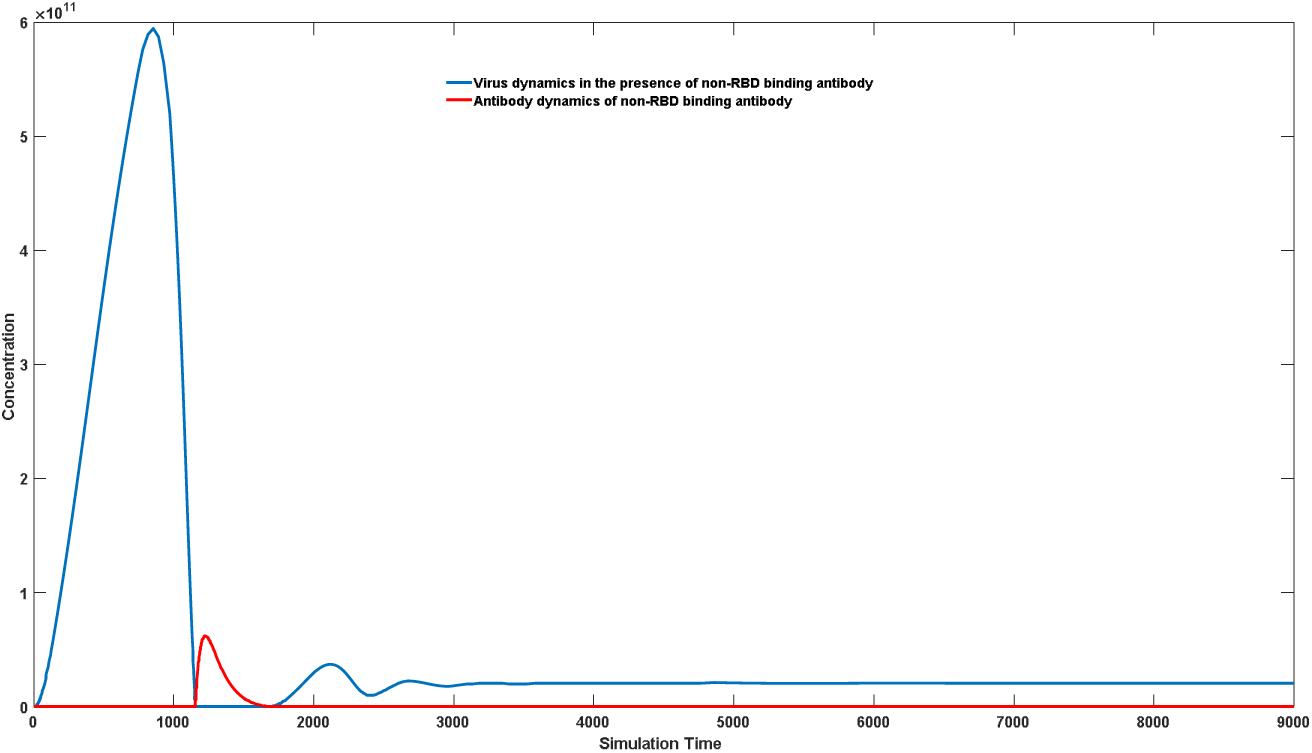
effect of non-RBD antibody on virus dynamics

**Figure 10b:**
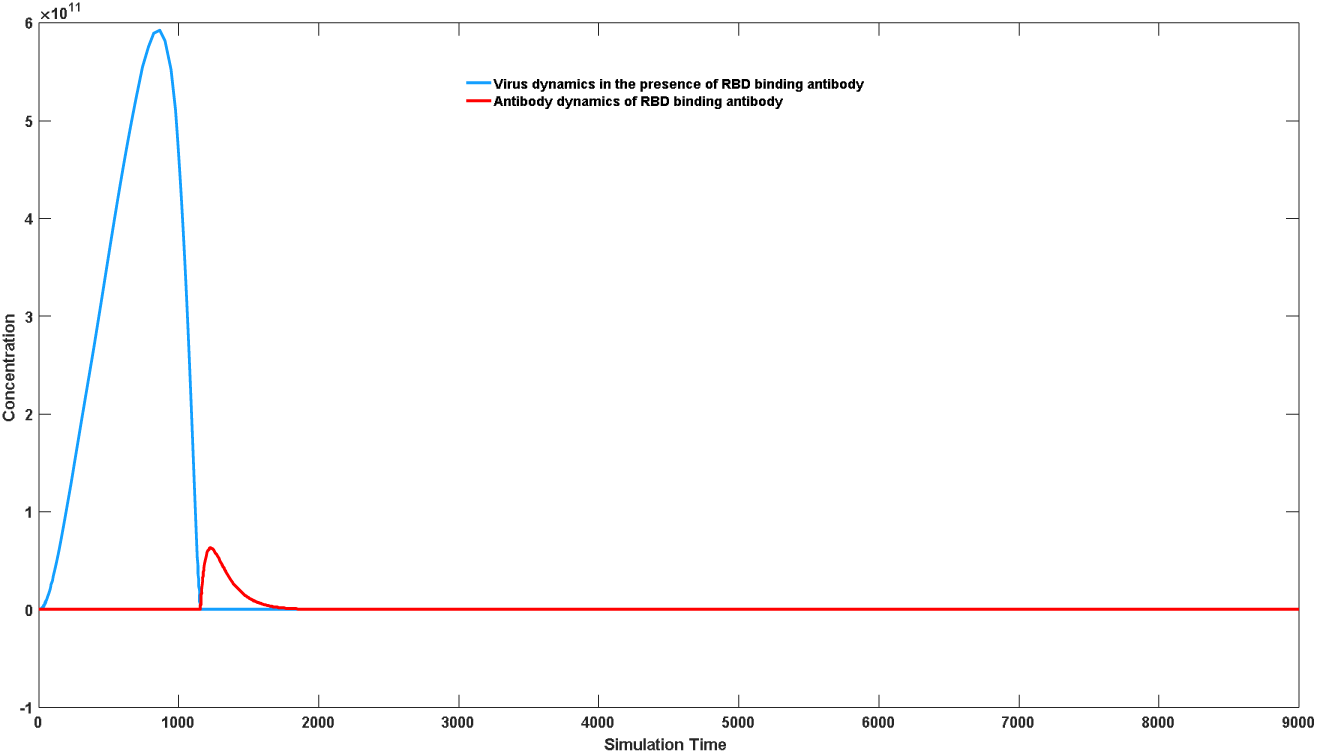
effect of RBD antibody on virus dynamics

This can explain the differences in symptoms among different HIV-infected individuals, as a small number of elite controller can achieve self-control of the virus without the need for drug intervention [56–57], which may be attributed to the induction of specific neutralizing antibodies. Additionally, our simulations revealed that the strength of Th cell-targeted ADCC effects also influences the host’s ability to control the virus. This may be one of the reasons why a small number of patients do not require drug treatment. The specific model and parameter settings are provided in the supplementary materials. As shown in Figure 11, when strong ADCC effects are present, the virus is less likely to be controlled and will decline to a relatively low steady state after the initial peak of viral infection (dashed blue line in Figure 11a), while antibodies are difficult to be successfully induced, resulting in consistently low antibody levels (dashed purple line in Figure 11a). Meanwhile, the total number of infected cells remains at a high level (indicated by dashed purple line in Figure 11b), while the total number of healthy cells remains at a very low level (indicated by dashed yellow line in Figure 11b). In such cases, severe chronic infection occurs, and patients often require reverse transcriptase inhibitors. In contrast, when the host’s Th cell-mediated ADCC effects are weak, the virus is more easily controlled. Although the virus may undergo repeated cycles (solid blue line in Figure 11a), each time it is effectively cleared with the help of antibodies (solid red line in Figure 11a). At the same time, the total number of infected cells fluctuates significantly (solid red line in Figure 11b), rather than remaining consistently high, and the total number of healthy cells (solid blue line in Figure 11b) also fluctuates, showing an upward trend, rather than remaining consistently low. These manifestations reduce the destructive effects of the HIV virus on the immune system. Figure 11 is generated using model 2.3. The detailed description and parameter set can be referred in supplementary materials (model 2.3).

**Figure 11a:**
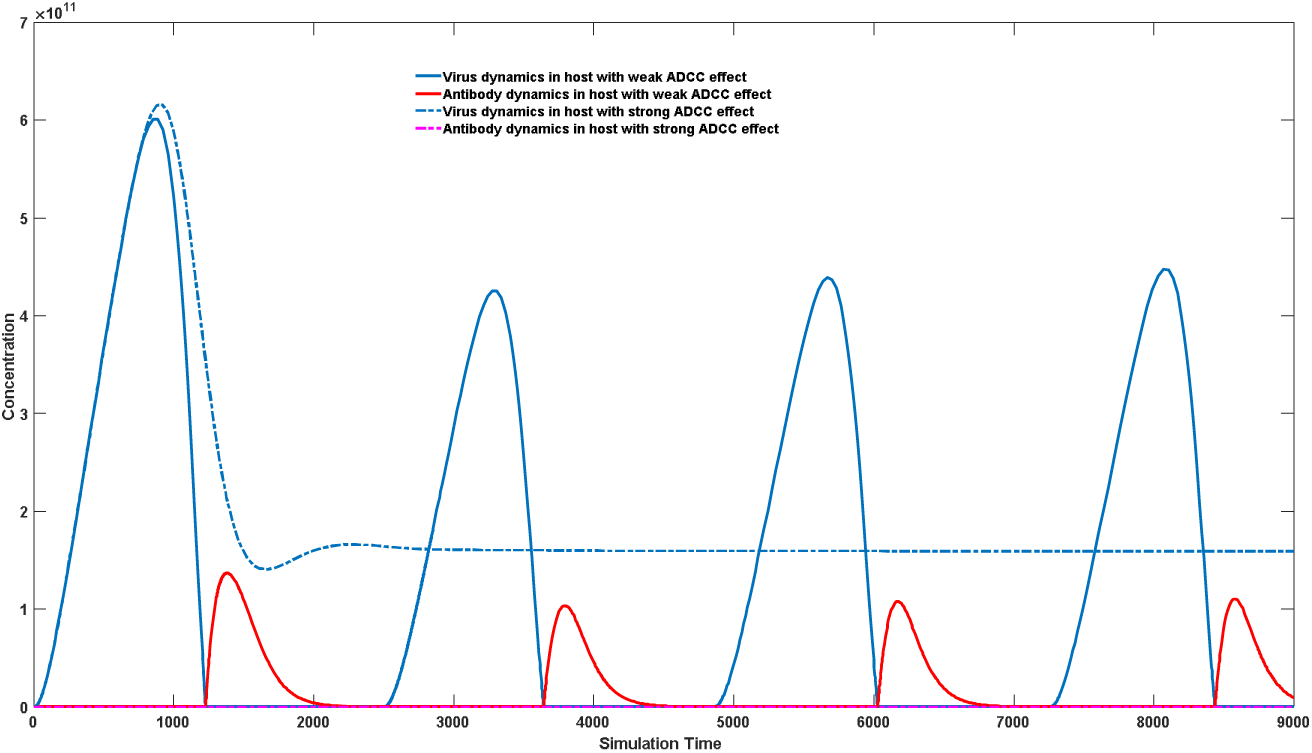
Antibody-virus dynamics in host with different Th-cell targeted ADCC effect

**Figure 11b:**
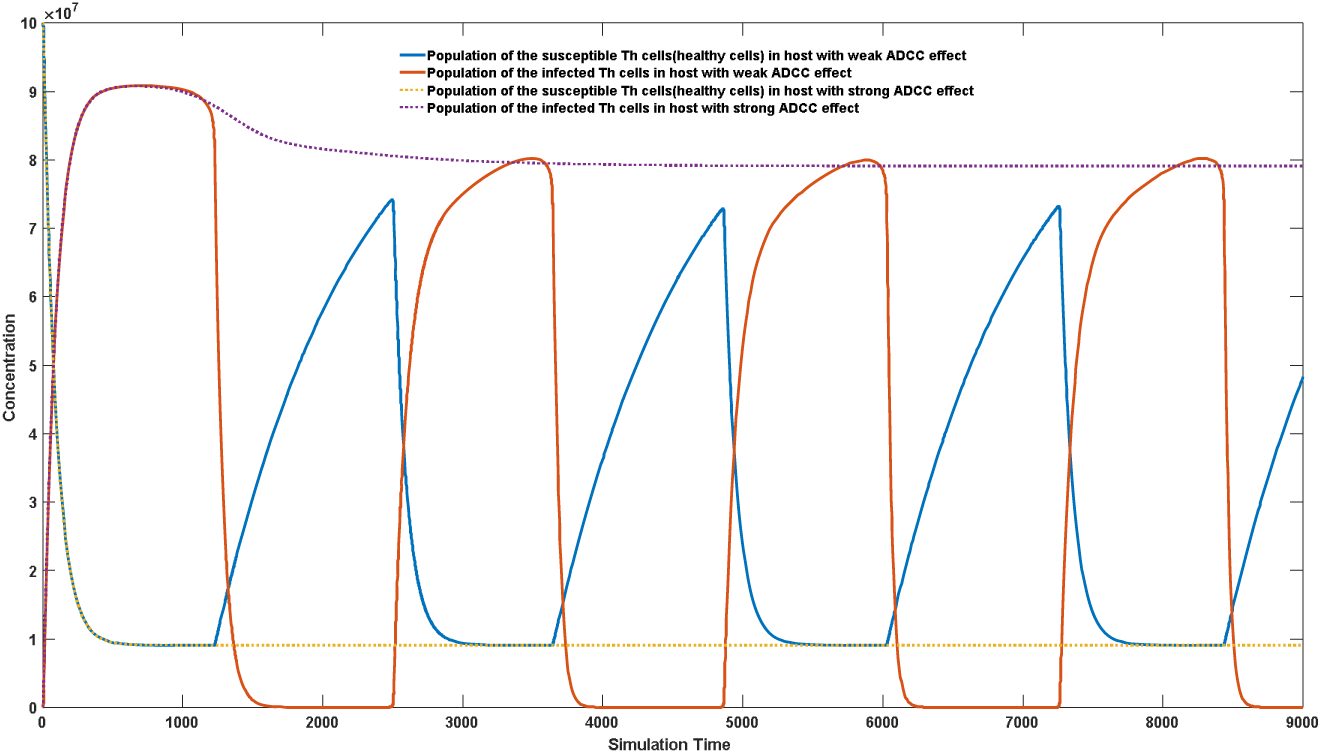
Th cell dynamics in host with different Th-cell targeted ADCC effect

### 3.6 Antibody treatment can effectively inhibit virus replication and even achieve virus clearance

With the discovery of broadly neutralizing antibodies, the use of this treatment modality for HIV/AIDS has become a trend [5–14]. It has been shown that the use of broadly neutralizing antibodies against specific viral strains can effectively control HIV replication in vivo, with inhibitory effects even superior to those of traditional reverse transcriptase inhibitors [58–60]. Herein, we demonstrate through simulation the effectiveness of antibody therapy, which, when used without reverse transcriptase inhibitors, may potentially cure HIV from its roots, as the CD4+ stem cells are not susceptible to HIV invasion. The body only needs to completely eliminate susceptible Th cells while relying on high concentrations of external antibodies to prevent the virus from invading new Th cells, gradually achieving complete virus clearance. A major problem with the use of reverse transcriptase inhibitors is that it inhibits virus replication, resulting in many infected Th cells being in a dormant state, rendering ADCC ineffective, and thus unable to cure HIV from its roots. The model’s specific parameter settings are detailed in the supplementary materials. As shown in Figure 12, due to the Th cell-targeted ADCC effect, patients are unable to achieve complete virus clearance in their natural state (as demonstrated by the blue solid line in Figure 12a), and the concentration of neutralizing antibodies is not successfully stimulated (as demonstrated by the red solid line in Figure 12a). However, upon the addition of high concentrations of specific neutralizing antibodies, the antibody concentration instantaneously increases to a high level (5*10^11^), resulting in a significant decrease in virus concentration (as demonstrated by the blue dashed line in Figure 12a), which can be sustained for a long time. Moreover, long-term use of specific neutralizing antibodies may lead to a sustained decrease in infected Th cell levels, thereby achieving a complete cure for HIV. Figure 12b shows the trend of changes in healthy and infected Th cell levels before and after antibody treatment, with healthy Th cell concentrations sharply decreasing before treatment (as demonstrated by the blue solid line in Figure 12b) but gradually returning to normal after treatment (as demonstrated by the blue dashed line in Figure 12b). Meanwhile, the infected Th cell level remains high before treatment (as demonstrated by the red solid line in Figure 12b), but rapidly decreases after treatment and can be maintained at a very low level for a certain period of time (as demonstrated by the red dashed line in Figure 12b). The latest reported antibody intervention studies have only lasted for several months [58–60]. Our simulation results show that with sufficient time and concentration of antibody treatment, achieving complete cure for HIV is highly feasible. Figure 12 is generated using model 2.4. The detailed description and parameter set can be referred in supplementary materials (model 2.4).

**Figure 12a:**
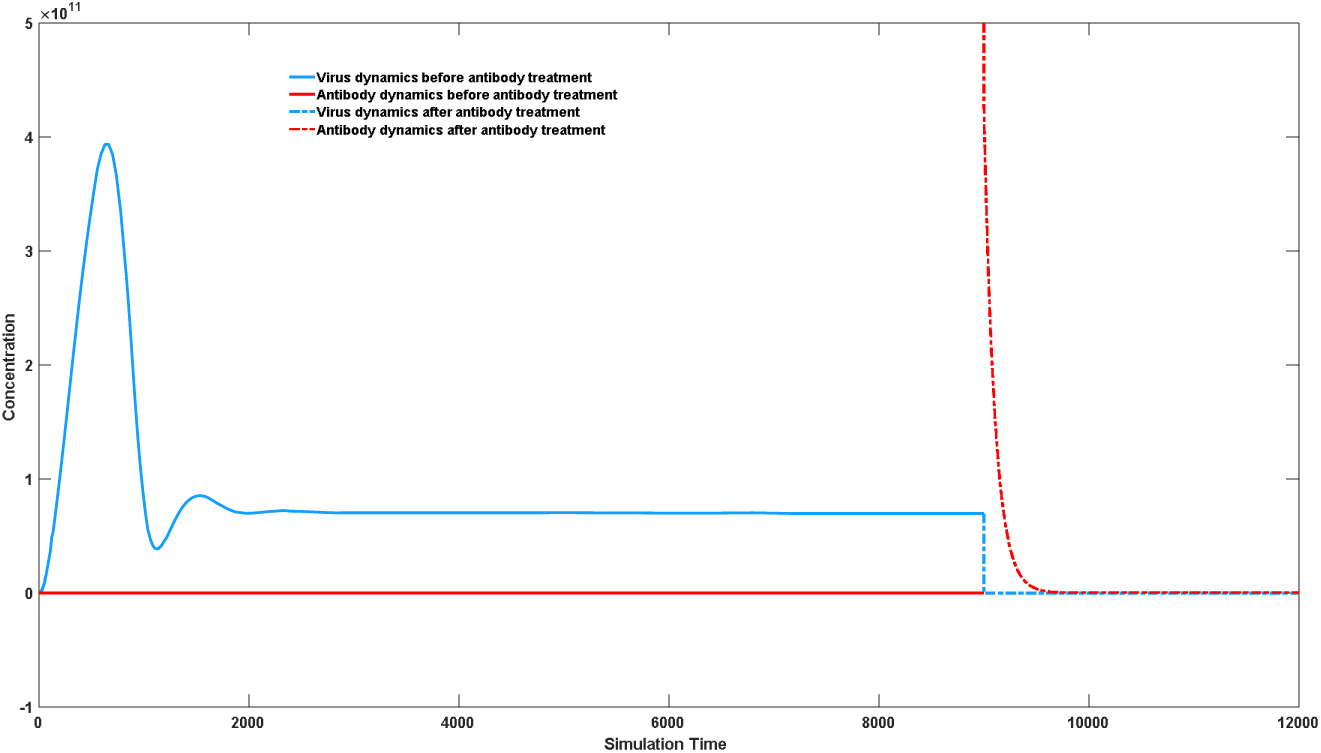
Virus dynamics after antibody treatment

**Figure 12b:**
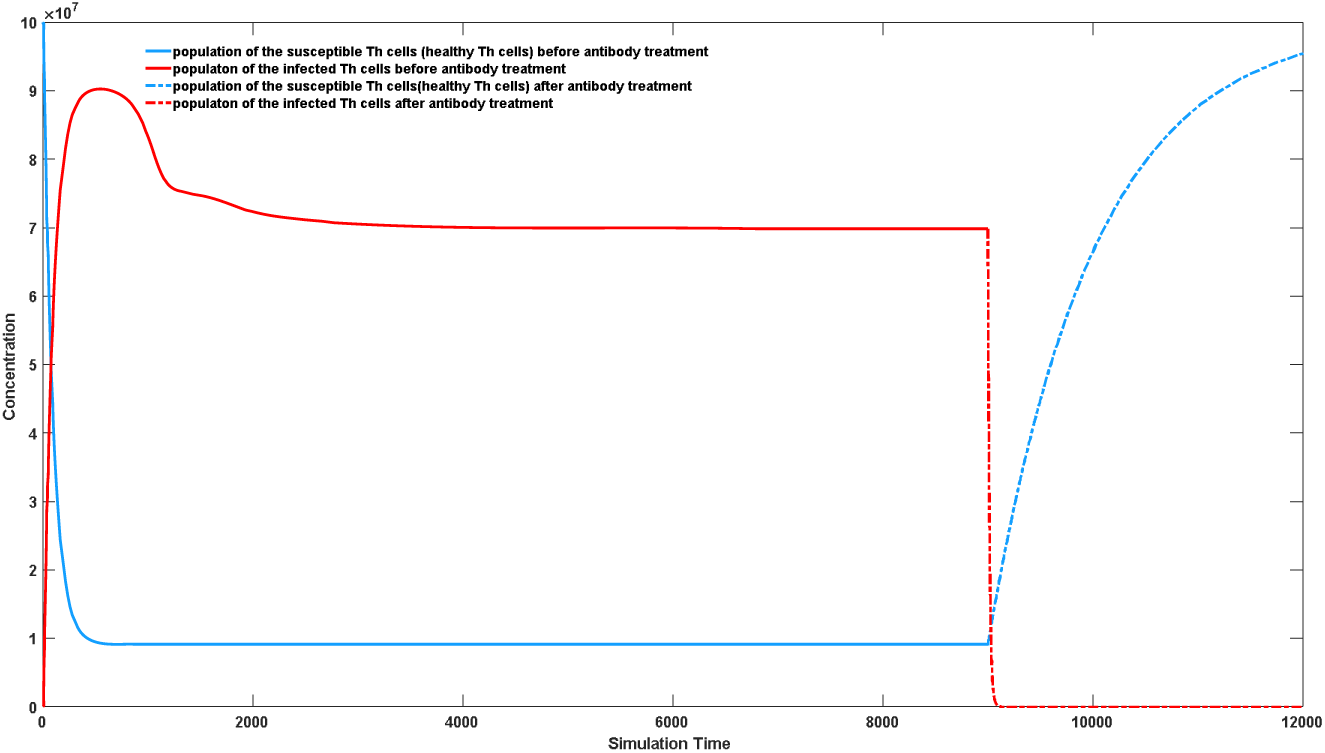
Th cell dynamics after antibody treatment

### 3.7 Inducing HIV-Specific Antibodies Using Bispecific Antibody

Our mathematical model reveals the Th cell-targeted Antibody-Dependent Cellular Cytotoxicity (ADCC) effect, which prevents complete clearance of HIV by the immune system during natural infection, leading to long-term chronic infection. This poses a challenge in vaccine development. If viral antigen substances, such as gp140, that can bind to CD4 molecules are used, it inevitably leads to the binding of the antigen substance to effector Th cells. The ADCC effect against Th cells generated during antibody induction will result in the apoptosis of germinal centers, thus preventing the generation of specific neutralizing antibodies. This may explain why many vaccine experiments have shown promising results in animals but often disappoint in clinical trials due to the non-specific binding of the antigen molecule to CD4+ cells in mice and other animals. Alternatively, if other antigens that do not bind to CD4 molecules are used to induce sufficient antibodies, these antibodies will target non-CD4 binding epitopes of HIV. Although they can accelerate viral clearance, they cannot prevent the virus from entering Th cells, thus failing to provide adequate protection. Our simulations confirm this finding.

How can we resolve this dilemma in HIV vaccine development? We propose the use of bispecific antibodies, as illustrated in Figure 13a. Bispecific antibodies are engineered antibodies that can simultaneously bind two different targets. They possess unique structures that allow recognition and interaction with different antigens or receptors. By targeting multiple molecules simultaneously, bispecific antibodies have diverse applications in various therapeutic and diagnostic settings. The mechanism of action of bispecific antibodies involves binding to two different targets, including tumor cell surface antigens and immune cells associated with disease occurrence. This ability to bridge different cell types or molecules enables bispecific antibodies to facilitate immune responses and enhance treatment efficacy. Common formats of bispecific antibodies include dual-variable domain, bispecific T-cell engagers (BiTEs), and dual-targeting antibodies. Currently, bispecific antibodies are predominantly used in cancer immunotherapy to achieve targeted recognition of specific cancer cells by immune cells [61–62]. However, there are no reports on the use of bispecific antibodies in vaccine development.

**Figure 13a:**
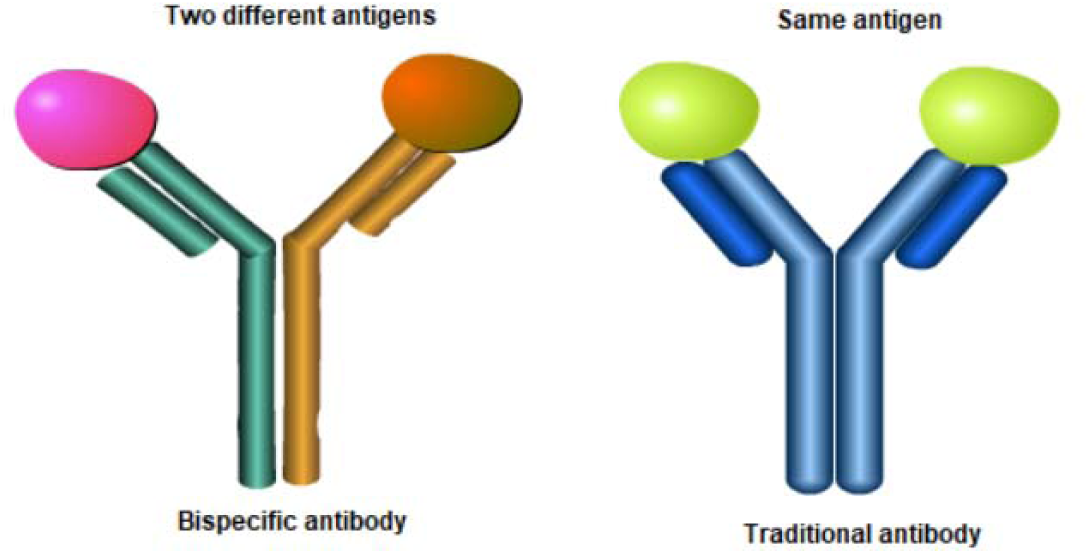
An illustration of bispecific antibody

The principle of inducing HIV-specific antibodies using bispecific antibodies is illustrated in Figure 13b. We employ a gene sequence targeting a broad-spectrum antibody against HIV and fuse it with another gene sequence specific to other pathogenic microorganisms or antigens. This gives rise to an antibody with two distinct properties. Unlike traditional antibodies, it transforms into a heterodimer, referred to as a bispecific antibody. One end of the bispecific antibody’s antigen recognition region can recognize antigens of other pathogenic microorganisms, such as the antibody sequence targeting the spike protein of SARS-CoV-2, while the other end can specifically recognize the CD4 binding region of gp120. We can use gene editing [63] or viral vector approaches [64–65] to insert this bispecific antibody into the B cells of patients, which can then be reintroduced into their bodies. Simultaneously, we can use another microbial antigen, such as the spike protein, as a vaccine to achieve in vivo amplification of the bispecific antibodies. This approach can elevate the levels of broad-spectrum neutralizing antibodies against HIV without directly administering HIV antigens. This strategy effectively circumvents the Th cell ADCC effect induced by HIV antigens. Moreover, utilizing an antigen to induce antibodies against other antigens in vaccine development represents a significant leap forward. Another reason for employing the method of inserting bispecific antibodies is that although traditional vaccination methods may be relatively safe, the number of B cells in an individual’s antibody repertoire that specifically target the gp120 binding region is very limited. High-affinity specific neutralizing antibodies isolated from patients often undergo extensive somatic hypermutation. Therefore, traditional vaccination methods may not be able to induce large-scale proliferation of this type of neutralizing antibodies within a short timeframe. Considering this, in recent years, James Voss and Adi Barzel [63–65] have attempted to directly insert these specific neutralizing antibodies using the principles of immunology. By stimulating the massive proliferation of these specific neutralizing antibodies through secondary antigen stimulation, they aim to prevent HIV infection. However, even with targeted insertion of antibody genes, due to the Th cell-targeted ADCC effect in the human body, it is possible that this method may fail to induce significant antibody proliferation. Therefore, we propose the use of bispecific antibodies for targeted insertion.

**Figure 13b:**
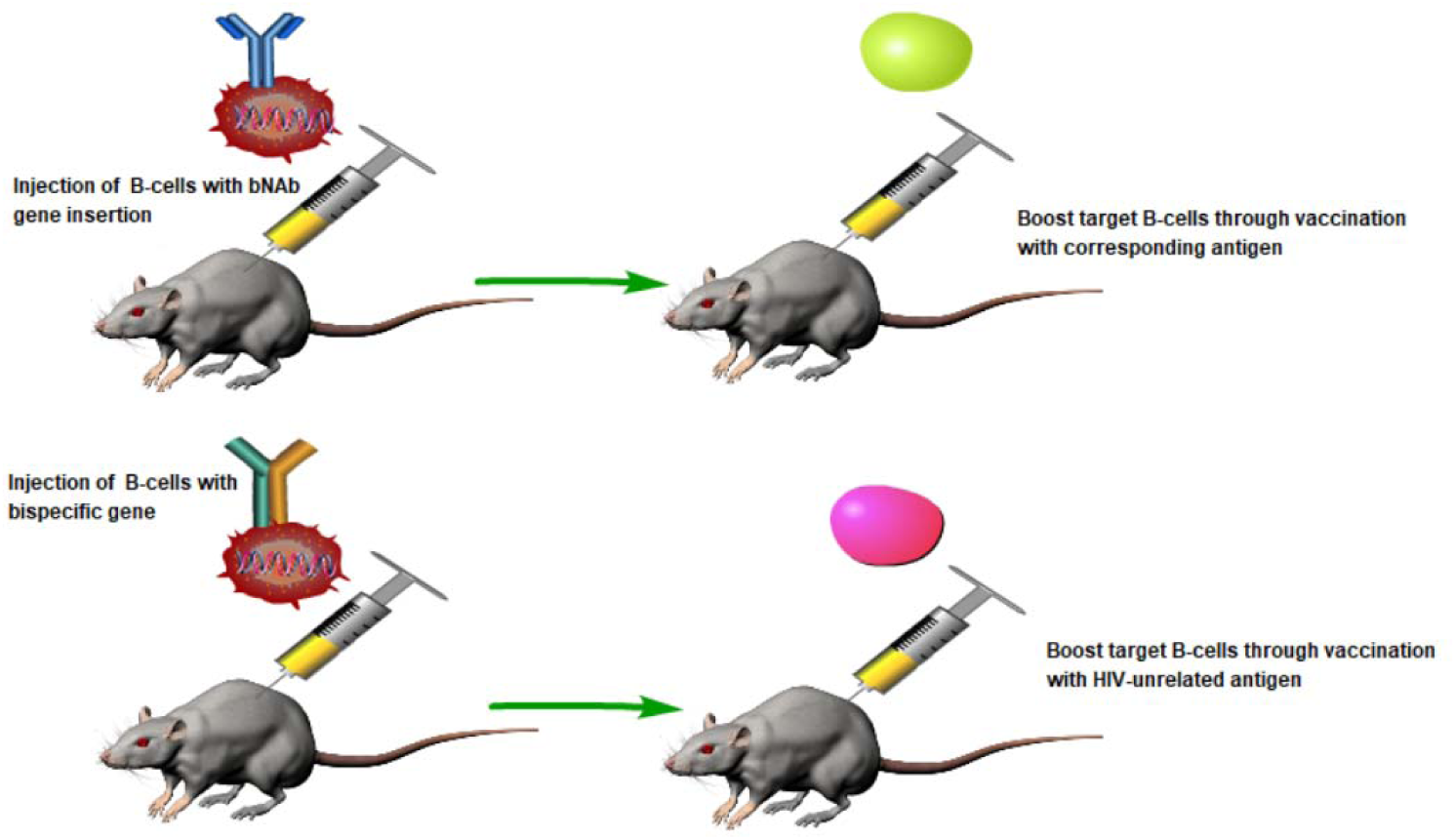
An illustration of HIV-bispecific antibody vaccination strategy

## 4 Discussion

Establishing a scientific mathematical model can help solve many experimental problems that are difficult to address otherwise. In this study, we inherited and developed our antibody kinetics model, focusing mainly on the effects of ADCC (antibody-dependent cell-mediated cytotoxicity) against B cells and Th cells on the specificity of immune responses. Specifically, we examined the Th cell-targeted ADCC effects, which we believe to have a direct impact on the interaction between HIV and the human immune system.

Firstly, we introduced how the Th cell immunogenicity of antigenic substances affects the fate of antibody production. For self-antigens, a relatively low Th cell immunogenicity is crucial for maintaining self-immune balance. Excessive Th cell immunogenicity or activity towards self-antigen substances can lead to autoimmune diseases, while low Th cell immunogenicity or activity can reduce the sensitivity and effectiveness of the immune system in monitoring cancers. Interestingly, relatively damaged Th cell activity can effectively control excessive proliferation of self-antigen substances but also generates a large amount of autoantibodies, leading to chronic inflammation. Therefore, the occurrence of many chronic inflammations is related to the decline in Th cell activity.

Regarding the effects of ADCC against B cells, our model successfully demonstrated that B cell-targeted ADCC effects can significantly increase the viral peak load, as well as the concentration of antibody-virus complex peaks, leading to more severe infection symptoms. Meanwhile, we explained why the formation of symmetrical spherical or polyhedral structures during virus assembly helps enhance transmission efficiency, thereby providing evolutionary advantages. We also proposed the idea of using molecular biology methods to directly use homologous oligomers of antigenic substances to eliminate autoantibodies.

We focused on Th cell-targeted ADCC effects, which, although traditional views emphasize the rapid mutation rate of HIV and its lengthy latency period, our simulation results suggest that the fundamental reason why HIV is difficult to clear lies in the ADCC effects against Th cells generated during HIV infection. Unlike B cell-targeted ADCC effects, Th cell-targeted ADCC effects are exacerbated due to the spatial proximity between Th cells and germinal center B cells. This leads to a peculiar phenomenon where high-affinity antibodies are not easily stimulated and may be surpassed by low-affinity antibodies in terms of proliferation rate, which is why most patients find it difficult to produce specific neutralizing antibodies. We also explained why some populations can achieve better virus control without using HIV drugs, which is related to the properties of antibodies. Combined with the specificity of neutralizing antibodies against the CD4 binding region of the HIV virus, better virus control effects can be achieved because they cannot only kill viruses but also significantly hinder the virus from entering new Th cells. We further studied the differences in individual ADCC effects that can lead to differences in virus control abilities, and strong Th cell ADCC effects can reduce the immune system’s suppression of the HIV virus.

We further simulated the effect of antibody therapy on HIV, and the results showed that periodic injection of exogenous broad-spectrum neutralizing antibodies can effectively inhibit virus proliferation while efficiently clearing infected Th cells. At the same time, our simulation also showed that using a sufficient dose of specific neutralizing antibodies to treat HIV, giving sufficient treatment time, may achieve the complete clearance of the virus, that is, achieving a cure.

Finally, we proposed a possible vaccine scheme that could address the Th cell-targeted ADCC effects of HIV infection, namely, inducing the proliferation of bispecific antibodies. Although bispecific antibodies have been widely used in cancer immunotherapy, there are currently no reports of their application in vaccine design. Using gene editing techniques to insert bispecific antibody genes into the recipient’s body, followed by immune stimulation with non-HIV antigens, according to immunological principles, can increase the concentration of both antibody substances and HIV-specific neutralizing antibodies, thus effectively avoiding the occurrence of Th cell-targeted ADCC effects and achieving efficient induction of antibody production.

We must acknowledge the limitations of our work: many parameter settings in this study lack strong experimental evidence and can only stay at the level of pattern analysis. This, to some extent, affects the credibility of our work. Moreover, although the simple model considering the virus-antibody interaction is simple, it may overlook many biological mechanisms. The simple model without cell typing does not consider the role of Th cells in antigen-antibody interactions. Although complex models consider multiple mechanisms, they inevitably bring too many parameters, and although some phenomena such as the growth rate of germinal centers can be qualitatively observed, quantitative data fitting is difficult to carry out. Therefore, the setting of these parameters inevitably introduces randomness, which also affects the reliability of the model. Our theory is just a hypothesis, although it can explain many macroscopic phenomena in immunology, more accurate conclusions require more experimental data support and verification. We hope that more experimental work can verify the Th cell-targeted ADCC effects and the bispecific antibody theory in HIV vaccine design.

## Codes availability

Matlab source codes are available at https://github.com/zhaobinxu23/ADCC_HIV_modelling

## Author Contributions

Conceptualization, Z.X.; methodology, Z.X. and D.Y; writing—original draft preparation, Z.X.; writing—review and editing, D.W., J.D. and H.Z.; funding acquisition, Z.X. All authors have read and agreed to the published version of the manuscript.

## Funding

This research was funded by DeZhou University, grant number 30101418.

## Acknowledgments

We thank Dr Zuyi Huang from Villanova University. Yushan Zhu from Tsinghua University, Jian Song, Peiyan Guan, XiangYong Li, and Zhenlin Wei from Dezhou University for helpful conversations, comments, and clarifications.

## Conflicts of Interest

The authors declare no conflict of interest.

## Notes

### Competing Interest Statement

The authors have declared no competing interest.

## References

1. Tatt I D, Barlow K L, Nicoll A, et al. The public health significance of HIV-1 subtypes[J]. Aids, 2001, 15: S59–S71.

2. Hemelaar J, Elangovan R, Yun J, et al. Global and regional molecular epidemiology of HIV-1, 1990–2015: a systematic review, global survey, and trend analysis[J]. The Lancet infectious diseases, 2019, 19(2): 143–155.

3. Williams A, Menon S, Crowe M, et al. Geographic and Population Distributions of Human Immunodeficiency Virus (HIV)–1 and HIV-2 Circulating Subtypes: A Systematic Literature Review and Meta-analysis (2010–2021)[J]. The Journal of Infectious Diseases, 2023, 228(11): 1583–1591.

4. Giovanetti M, Ciccozzi M, Parolin C, et al. Molecular epidemiology of HIV-1 in African countries: a comprehensive overview[J]. Pathogens, 2020, 9(12): 1072.

5. Sok D, Burton D R. Recent progress in broadly neutralizing antibodies to HIV[J]. Nature immunology, 2018, 19(11): 1179–1188.

6. Burton D R, Hangartner L. Broadly neutralizing antibodies to HIV and their role in vaccine design[J]. Annual review of immunology, 2016, 34: 635–659.

7. Stephenson K E, Barouch D H. Broadly neutralizing antibodies for HIV eradication[J]. Current Hiv/aids Reports, 2016, 13: 31–37.

8. McLellan J S, Pancera M, Carrico C, et al. Structure of HIV-1 gp120 V1/V2 domain with broadly neutralizing antibody PG9[J]. Nature, 2011, 480(7377): 336–343.

9. Gorman J, Soto C, Yang M M, et al. Structures of HIV-1 Env V1V2 with broadly neutralizing antibodies reveal commonalities that enable vaccine design[J]. Nature structural & molecular biology, 2016, 23(1): 81–90.

10. Scharf L, West Jr A P, Gao H, et al. Structural basis for HIV-1 gp120 recognition by a germ-line version of a broadly neutralizing antibody[J]. Proceedings of the National Academy of Sciences, 2013, 110(15): 6049–6054.

11. McCoy L E, Burton D R. Identification and specificity of broadly neutralizing antibodies against HIV[J]. Immunological reviews, 2017, 275(1): 11–20.

12. Matz J, Kessler P, Bouchet J, et al. Straightforward selection of broadly neutralizing single-domain antibodies targeting the conserved CD4 and coreceptor binding sites of HIV-1 gp120[J]. Journal of virology, 2013, 87(2): 1137–1149.

13. Kwong P D, Mascola J R. Human antibodies that neutralize HIV-1: identification, structures, and B cell ontogenies[J]. Immunity, 2012, 37(3): 412–425.

14. West A P, Scharf L, Scheid J F, et al. Structural insights on the role of antibodies in HIV-1 vaccine and therapy[J]. Cell, 2014, 156(4): 633–648.

15. Ng’uni T, Chasara C, Ndhlovu Z M. Major scientific hurdles in HIV vaccine development: historical perspective and future directions[J]. Frontiers in immunology, 2020, 11: 590780.

16. Kim J, Vasan S, Kim J H, et al. Current approaches to HIV vaccine development: a narrative review[J]. Journal of the International AIDS Society, 2021, 24: e25793.

17. A Day T, G Kublin J. Lessons learned from HIV vaccine clinical efficacy trials[J]. Current HIV research, 2013, 11(6): 441–449.

18. Watkins D I, Burton D R, Kallas E G, et al. Nonhuman primate models and the failure of the Merck HIV-1 vaccine in humans[J]. Nature medicine, 2008, 14(6): 617–621.

19. Xu Z, Wei D, Zhang H, et al. A Novel Mathematical Model That Predicts the Protection Time of SARS-CoV-2 Antibodies[J]. Viruses, 2023, 15(2): 586.

20. Hong S, Zhang Z, Liu H, et al. B cells are the dominant antigen-presenting cells that activate naive CD4+ T cells upon immunization with a virus-derived nanoparticle antigen[J]. Immunity, 2018, 49(4): 695–708. e4.

21. Rastogi I, Jeon D, Moseman J E, et al. Role of B cells as antigen presenting cells[J]. Frontiers in Immunology, 2022, 13: 954936.

22. Hua Z, Hou B. The role of B cell antigen presentation in the initiation of CD4+ T cell response[J]. Immunological reviews, 2020, 296(1): 24–35.

23. Mujal A M, Delconte R B, Sun J C. Natural killer cells: From innate to adaptive features[J]. Annual review of immunology, 2021, 39: 417–447.

24. Li G, Iyer B, Prasath V B S, et al. DeepImmuno: deep learning-empowered prediction and generation of immunogenic peptides for T-cell immunity[J]. Briefings in bioinformatics, 2021, 22(6): bbab160.

25. Mei S, Li F, Leier A, et al. A comprehensive review and performance evaluation of bioinformatics tools for HLA class I peptide-binding prediction[J]. Briefings in bioinformatics, 2020, 21(4): 1119–1135.

26. Liu Y J, Arpin C. Germinal center development[J]. Immunological reviews, 1997, 156(1): 111–126.

27. Wang Y, Shi J, Yan J, et al. Germinal-center development of memory B cells driven by IL-9 from follicular helper T cells[J]. Nature immunology, 2017, 18(8): 921–930.

28. Heesters B A, van der Poel C E, Das A, et al. Antigen presentation to B cells[J]. Trends in immunology, 2016, 37(12): 844–854.

29. Xu Z, Peng Q, Liu W, et al. Antibody Dynamics Simulation—A Mathematical Exploration of Clonal Deletion and Somatic Hypermutation[J]. Biomedicines, 2023, 11(7): 2048.

30. Gaudin E, Rosado M, Agenes F, et al. B-cell homeostasis, competition, resources, and positive selection by self-antigens[J]. Immunological reviews, 2004, 197(1): 102–115.

31. Maruyama M, Lam K P, Rajewsky K. Memory B-cell persistence is independent of persisting immunizing antigen[J]. Nature, 2000, 407(6804): 636–642.

32. Gray D, Skarvall H. B–cell memory is short-lived in the absence of antigen[J]. Nature, 1988, 336(6194): 70–73.

33. Martincorena I, Campbell P J. Somatic mutation in cancer and normal cells[J]. Science, 2015, 349(6255): 1483–1489.

34. Milholland B, Auton A, Suh Y, et al. Age-related somatic mutations in the cancer genome[J]. Oncotarget, 2015, 6(28): 24627.

35. Abascal F, Harvey L M R, Mitchell E, et al. Somatic mutation landscapes at single-molecule resolution[J]. Nature, 2021, 593(7859): 405–410.

36. Rhodes D R, Chinnaiyan A M. Integrative analysis of the cancer transcriptome[J]. Nature genetics, 2005, 37(Suppl 6): S31–S37.

37. Uhlen M, Zhang C, Lee S, et al. A pathology atlas of the human cancer transcriptome[J]. Science, 2017, 357(6352): eaan2507.

38. Rhodes D R, Kalyana-Sundaram S, Mahavisno V, et al. Mining for regulatory programs in the cancer transcriptome[J]. Nature genetics, 2005, 37(6): 579–583.

39. Tokunaga R, Naseem M, Lo J H, et al. B cell and B cell-related pathways for novel cancer treatments[J]. Cancer treatment reviews, 2019, 73: 10–19.

40. Tsou P, Katayama H, Ostrin E J, et al. The emerging role of B cells in tumor immunity[J]. Cancer research, 2016, 76(19): 5597–5601.

41. Coronella-Wood J A, Hersh E M. Naturally occurring B-cell responses to breast cancer[J]. Cancer Immunology, Immunotherapy, 2003, 52: 715–738.

42. Schwartz M, Zhang Y, Rosenblatt J D. B cell regulation of the anti-tumor response and role in carcinogenesis[J]. Journal for immunotherapy of cancer, 2016, 4: 1–15.

43. Zhaobin Xu, 。。。。。。The Mathematical Modeling of the Host–Virus Interaction in Dengue Virus Infection: A Quantitative Study[J]. MDPI Viruses, 2024, 4: 1–15.

44. Johnson S N, Brucks S D, Apley K D, et al. Multivalent Scaffolds to Promote B cell Tolerance[J]. Molecular Pharmaceutics, 2023, 20(8): 3741–3756.

45. Griffin J D, Leon M A, Salash J R, et al. Acute B-cell inhibition by soluble antigen arrays is valency-dependent and predicts immunomodulation in splenocytes[J]. Biomacromolecules, 2019, 20(5): 2115–2122.

46. Hartwell B L, Pickens C J, Leon M, et al. Soluble antigen arrays disarm antigen-specific B cells to promote lasting immune tolerance in experimental autoimmune encephalomyelitis[J]. Journal of autoimmunity, 2018, 93: 76–88.

47. Tostanoski L H, Chiu Y C, Andorko J I, et al. Design of polyelectrolyte multilayers to promote immunological tolerance[J]. ACS nano, 2016, 10(10): 9334–9345.

48. Rydyznski C, Daniels K A, Karmele E P, et al. Generation of cellular immune memory and B-cell immunity is impaired by natural killer cells[J]. Nature communications, 2015, 6(1): 6375.

49. Cook, K.D.; Kline, H.C.; Whitmire, J.K. NK cells inhibit humoral immunity by reducing the abundance of CD4+ T follicular helper cells during a chronic virus infection. J. Leucoc. Biol. 2015, 98, 153–162

50. O’Leary J G, Goodarzi M, Drayton D L, et al. T cell–and B cell–independent adaptive immunity mediated by natural killer cells[J]. Nature immunology, 2006, 7(5): 507–516.

51. Bogers W M J M, Barnett S W, Oostermeijer H, et al. Increased, durable B-cell and ADCC responses associated with T-helper cell responses to HIV-1 envelope in macaques vaccinated with gp140 occluded at the CD4 receptor binding site[J]. Journal of Virology, 2017, 91(19): 10.1128/jvi.00811-17.

52. Hraber P, Seaman M S, Bailer R T, et al. Prevalence of broadly neutralizing antibody responses during chronic HIV-1 infection[J]. AIDS (London, England), 2014, 28(2): 163.

53. Robinson H L. Non-neutralizing antibodies in prevention of HIV infection[J]. Expert opinion on biological therapy, 2013, 13(2): 197–207.

54. Srivastava I K, Ulmer J B, Barnett S W. Role of neutralizing antibodies in protective immunity against HIV[J]. Human vaccines, 2005, 1(2): 45–60.

55. Parsons M S, Le Grand R, Kent S J. Neutralizing antibody-based prevention of cell-associated HIV-1 infection[J]. Viruses, 2018, 10(6): 333.

56. Kumar P. Long term non-progressor (LTNP) HIV infection[J]. The Indian journal of medical research, 2013, 138(3): 291.

57. Navarrete-Muñoz M A, Restrepo C, Benito J M, et al. Elite controllers: A heterogeneous group of HIV-infected patients[J]. Virulence, 2020, 11(1): 889–897.

58. Mendoza P, Gruell H, Nogueira L, et al. Combination therapy with anti-HIV-1 antibodies maintains viral suppression[J]. Nature, 2018, 561(7724): 479–484.

59. Sneller M C, Blazkova J, Justement J S, et al. Combination anti-HIV antibodies provide sustained virological suppression[J]. Nature, 2022, 606(7913): 375–381.

60. Caskey M, Klein F, Nussenzweig M C. Broadly neutralizing anti-HIV-1 monoclonal antibodies in the clinic[J]. Nature medicine, 2019, 25(4): 547–553.

61. Ma J, Mo Y, Tang M, et al. Bispecific antibodies: from research to clinical application[J]. Frontiers in Immunology, 2021, 12: 1555.

62. Brinkmann U, Kontermann R E. Bispecific antibodies[J]. Science, 2021, 372(6545): 916–917.

63. Voss J E, Gonzalez-Martin A, Andrabi R, et al. Reprogramming the antigen specificity of B cells using genome-editing technologies[J]. Elife, 2019, 8: e42995.

64. Huang D, Tran J T, Olson A, et al. Vaccine elicitation of HIV broadly neutralizing antibodies from engineered B cells[J]. Nature communications, 2020, 11(1): 5850.

65. Nahmad A D, Lazzarotto C R, Zelikson N, et al. In vivo engineered B cells secrete high titers of broadly neutralizing anti-HIV antibodies in mice[J]. Nature biotechnology, 2022, 40(8): 1241–1249.

