## Supplemental materials for "Mathematical Modelling Indicates Th-cell Targeted Antibody-Dependent Cellular Cytotoxic Is a Crucial Obstacle Hurdling HIV Vaccine Development"

Supplementary materials

Model 1.1

Model 1.1 is used to simulate the impact of antigenic Th immunogenicity on acquired immunity, generating Figure 3. It consists of two antibodies with different affinity constants: antibody 1 with a high binding affinity has a binding constant of k2 = 1e-6, while antibody 2 with a lower binding affinity has a binding constant of k7 = 0.5e-6. When the antigen is an exogenous antigen, such as a virus, it can undergo self-proliferation, hence k1 = 0.1. In the case of strong Th cell immunogenicity of the antigen, k3 = k8 = 1 (Figure 3a). Conversely, when the antigen possesses weak Th cell immunogenicity, k3 = k8 = 0.2 (Figure 3b).

| Reaction index | Reaction |
| --- | --- |
| 1 | Antigen Antigen |
| 2 | Antibody-1 + Antigen Antibody-1-Antigen Complex |
| 3 | Antibody-1-Antigen Complex Antibody-1 |
| 4 | Antibody-1-Antigen Complex Cleared by immune system |
| 5 | Antigen Degradation |
| 6 | Antibody-1 Degradation |
| 7 | Antibody-2 + Antigen Antibody-2-Antigen Complex |
| 8 | Antibody-2-Antigen Complex Antibody-2 |
| 9 | Antibody-2-Antigen Complex Cleared by immune system |
| 10 | Antibody-2 Degradation |

Table S1: Reaction index and the name of each reaction in model 1.1

| Parameter Name | Description | Value | Range, refs |
| --- | --- | --- | --- |
| k_1_ | Antigen proliferation rate | 0.1 in Figure3a | [0.1~1], ref [1-3] |
| k_2_ | Forward binding constant between antibody-1 and antigen | 1e-6 | [1e-8~1e-5], ref [1-3] |
| k_-2_ | Dissociation constant of antibody-1-antigen complex | 1e-14 | 1e-14, ref [1-3] |
| k_3_ | Feedback constant of antibody-1-antigen complex on the regeneration of antibody-1 | 1 in Figure 3a  0.2 in Figure 3b | [0.1~2], ref [1-3] |
| k_4_ | Clearance rate of antibody-1-antigen complex | 0.4 | [0.1~1], ref [1-3] |
| k_5_ | Antigen degradation rate | 0.02 | [0.01~0.02], ref [1-3] |
| k_6_ | Antibody-1 degradation rate | 0.02 | [0.01~0.02], ref [1-3] |
| k_7_ | Forward binding constant between antibody-2 and antigen | 0.5e-6 | [1e-8~1e-5], ref [1-3] |
| k_-7_ | Dissociation constant of antibody-2-antigen complex | 1e-14 | 1e-14, ref [1-3] |
| k_8_ | Feedback constant of antibody-2-antigen complex on the regeneration of antibody-2 | 1 in Figure 3a  0.2 in Figure 3b | [0.1~2], ref [1-3] |
| k_9_ | Clearance rate of antibody-2-antigen complex | 0.4 | [0.1~1], ref [1-3] |
| k_10_ | Antibody-2 degradation rate | 0.02 | [0.01~0.02], ref [1-3] |

Table S2：Estimates of parameters in model 1.1.

| Variables | Meaning | Initial value |
| --- | --- | --- |
| $x1$ | Antigen | 10 in figure 3a  10^6^ in figure 3b |
| $x2$ | Antibody-1 | 100  10000 in figure 3b |
| $x3$ | Antibody-1-antigen complex | 0 |
| $x4$ | Antibody-2 | 100  10000 in figure 3b |
| $x5$ | Antibody-2-antigen complex | 0 |

Table S3：Time-dependent variables of model 1.1

The set of differential equations of model 1.1 is represented below:

$\frac{d(x1)}{dt}=k1*x1-k5*x1-k2*x1*x2+k_{-2}*x3-k7*x1*x4+k_{-7}*x5; (Figure 3a)$

$\frac{d\left( x1 \right)}{dt}=0; (Figure 3b)$

$\frac{d\left( x2 \right)}{dt}=-k2*x1*x2+k_{-2}*x3-k6*x2+k3*x3;$

$$\frac{d\left( x3 \right)}{dt}=k2*x1*x2-k_{-2}*x3-k4*x3;$$

$$\frac{d(x4)}{dt}=-k7*x1*x4+k_{-7}*x5-k9*x4+k8*x5;$$

$$\frac{d(x5)}{dt}=k7*x1*x4-k_{-7}*x5-k9*x5;$$

Model 1.2

Model 1.2 is utilized to elucidate how the Th cell immunogenicity of endogenous antigens can effectively control the occurrence of cancer. We have selected three sets of different gradient values for k3, specifically 0.5, 0.46, and 0.42, to represent strong Th cell immunogenicity, decreasing Th immunogenicity, and low Th cell immunogenicity, respectively.

| Reaction index | Reaction |
| --- | --- |
| 1 | Antigen Antigen |
| 2 | Antibody + Antigen Antibody-Antigen Complex |
| 3 | Antibody-Antigen Complex Antibody |
| 4 | Antibody-Antigen Complex Clearance by immune system |
| 5 | Antigen Degradation |
| 6 | Antibody Degradation |
| 7 | Constant replenishment from environment Antigen |

Table S4: Reaction index and the name of each reaction in model 1.2

| Parameter Name | Description | Value |
| --- | --- | --- |
| k_1_ | Antigen proliferation rate | 0 for healthy condition  0.1 for cancer cell |
| k_2_ | Forward binding constant between antibody and antigen | 1e-5 |
| k_-2_ | Dissociation constant of antibody-antigen complex | 1e-14 |
| k_3_ | Feedback constant of antibody-antigen complex on the regeneration of antibody | 0.50 for figure4a;  0.46 for figure 4b;  0.42 for figure 4c. |
| k_4_ | Clearance rate of antibody-antigen complex | 0.4 |
| k_5_ | Antigen degradation rate | 0.02 |
| k_6_ | Antibody degradation rate | 0.02 |
| c | Replenish constant of antigen | 200 |

Table S5：Estimates of parameters in model 1.2.

| Variables | Meaning | Initial value |
| --- | --- | --- |
| $x1$ | Antigen | 10000 |
| $x2$ | Antibody | 10000 |
| $x3$ | Antibody-antigen complex | 0 |

Table S6：Time-dependent variables of model 1.2

The set of differential equations of model 1.2 is represented below:

$\frac{d(x1)}{dt}=c+k1*x1-k5*x1-k2*x1*x2+k_{-2}*x3$

$\frac{d\left( x2 \right)}{dt}=-k2*x1*x2+k_{-2}*x3-k6*x2+k3*x3;$

$$\frac{d\left( x3 \right)}{dt}=k2*x1*x2-k_{-2}*x3-k4*x3;$$

Model 1.3

Model 1.3 is employed to simulate the impact of B cell-targeted Antibody-Dependent Cellular Cytotoxicity (ADCC) on antibody-virus interactions. In the presence of B cell-targeted ADCC effect, the clearance rate of antigen-antibody complexes is accelerated, with the k4 value increasing from 1 to 1.5.

| Reaction index | Reaction |
| --- | --- |
| 1 | Antigen Antigen |
| 2 | Antibody + Antigen Antibody-Antigen Complex |
| 3 | Antibody-Antigen Complex Antibody |
| 4 | Antibody-Antigen Complex Clearance by immune system |
| 5 | Antigen Degradation |
| 6 | Antibody Degradation |

Table S7: Reaction index and the name of each reaction in model 1.3

| Parameter Name | Description | Value |
| --- | --- | --- |
| k_1_ | Antigen proliferation rate | 0.1 |
| k_2_ | Forward binding constant between antibody and antigen | 1.5e-5 |
| k_-2_ | Dissociation constant of antibody-antigen complex | 1e-14 |
| k_3_ | Feedback constant of antibody-antigen complex on the regeneration of antibody | 2 |
| k_4_ | Clearance rate of antibody-antigen complex | 1.5 for B-cell targeted ADCC;  1 for control group |
| k_5_ | Antigen degradation rate | 0.02 |
| k_6_ | Antibody degradation rate | 0.02 |

Table S8：Estimates of parameters in model 1.3.

| Variables | Meaning | Initial value |
| --- | --- | --- |
| $x1$ | Antigen | 10 |
| $x2$ | Antibody | 100 |
| $x3$ | Antibody-antigen complex | 0 |

Table S9：Time-dependent variables of model 1.3

The set of differential equations of model 1.3 is represented below:

$\frac{d(x1)}{dt}=k1*x1-k5*x1-k2*x1*x2+k_{-2}*x3$

$\frac{d\left( x2 \right)}{dt}=-k2*x1*x2+k_{-2}*x3-k6*x2+k3*x3;$

$$\frac{d\left( x3 \right)}{dt}=k2*x1*x2-k_{-2}*x3-k4*x3;$$

Model 1.4

Model 1.4 is utilized to simulate the impact of Th cell-targeted Antibody-Dependent Cellular Cytotoxicity (ADCC) on antibody activation. Similarly, this model involves two types of antibodies: antibody 1 with high binding affinity (k2 = 1e-5) and antibody 2 with low binding affinity (k7 = 0.9e-5). For the Th cell-mediated ADCC effect, k11 = 1. In the control group without ADCC effect, k11 = 0. The Th cell-targeted ADCC effect is expressed in a form similar to the Michaelis-Menten equation, as k11*antibody concentration/(antibody concentration+ K_m_).

| Reaction index | Reaction |
| --- | --- |
| 1 | Antigen Antigen |
| 2 | Antibody-1 + Antigen Antibody-1-Antigen Complex |
| 3 | Antibody-1-Antigen Complex Antibody-1 |
| 4 | Antibody-1-Antigen Complex Clearance by immune system |
| 5 | Antigen Degradation |
| 6 | Antibody-1 Degradation |
| 7 | Antibody-2 + Antigen Antibody-2-Antigen Complex |
| 8 | Antibody-2-Antigen Complex Antibody-2 |
| 9 | Antibody-2-Antigen Complex Clearance by immune system |
| 10 | Antibody-2 Degradation |

Table S10: Reaction index and the name of each reaction in model 1.4

| Parameter Name | Description | Value |
| --- | --- | --- |
| k_1_ | Antigen proliferation rate | 0.1 |
| k_2_ | Forward binding constant between antibody-1 and antigen | 1e-5 |
| k_-2_ | Dissociation constant of antibody-1-antigen complex | 1e-14 |
| k_3_ | Feedback constant of antibody-1-antigen complex on the regeneration of antibody-1 | 1 |
| k_4_ | Clearance rate of antibody-1-antigen complex | 0.4 |
| k_5_ | Antigen degradation rate | 0.02 |
| k_6_ | Antibody-1 degradation rate | 0.02 |
| k_7_ | Forward binding constant between antibody-2 and antigen | 0.9e-5 |
| k_-7_ | Dissociation constant of antibody-2-antigen complex | 1e-14 |
| k_8_ | Feedback constant of antibody-2-antigen complex on the regeneration of antibody-2 | 1 |
| k_9_ | Clearance rate of antibody-2-antigen complex | 0.4 |
| k_10_ | Antibody-2 degradation rate | 0.02 |
| k_11_ | Maximal inhibition effect of antibody | 1 for Th-cell targeted ADCC; 0 for control group |
| K_m_ | Concentration of antibody with 50% maximal inhibition effect | 1e4 |

Table S11：Estimates of parameters in model 1.4.

| Variables | Meaning | Initial value |
| --- | --- | --- |
| $x1$ | Antigen | 10 |
| $x2$ | Antibody-1 | 100 |
| $x3$ | Antibody-1-antigen complex | 0 |
| $x4$ | Antibody-2 | 100 |
| $x5$ | Antibody-2-antigen complex | 0 |

Table S12：Time-dependent variables of model 1.4

The set of differential equations of model 1.4 is represented below:

$\frac{d(x1)}{dt}=k1*x1-k5*x1-k2*x1*x2+k_{-2}*x3 -k7*x1*x4+k_{-7}*x5;$

$\frac{d\left( x2 \right)}{dt}=-k2*x1*x2+k_{-2}*x3-k6*x2+(k3-k11*x2/(x2+Km)*x3;$

$$\frac{d\left( x3 \right)}{dt}=k2*x1*x2-k_{-2}*x3-k4*x3;$$

$$\frac{d\left( x4 \right)}{dt}=-k7*x1*x4+k_{-7}*x5-k10*x4+(k8-k11*x4/(x4+Km)*x5;$$

$$\frac{d\left( x5 \right)}{dt}=k7*x1*x4-k_{-7}*x5-k9*x5;$$

Model 2.1

Model 2.1 employs a cell-compartment model to simulate the impact of Th cell-targeted ADCC effect on antibody generation. For the control group of non-HIV virus-induced infections, the effect Th cells are not infected, and $\alpha_{1}$ = 0.

| Reaction Index | Reaction |
| --- | --- |
| 1 | S-Th + V  I-Th |
| 2 | I-Th +A1 Removed by immune system |
| 3 | I-Th +A2 Removed by immune system |
| 4 | B1 + V + S-Th B1-S-Th |
| 5 | B1 + V + I-Th B1-I-Th |
| 6 | B2 + V + S-Th B2-S-Th |
| 7 | B2 + V + I-Th B2-I-Th |
| 8 | A1 + V A1-V |
| 9 | A2 + V A2-V |
| 10 | I-Th Lysis and release viruses into body fluids |
| 11 | B1-I-Th +A1 Removed by immune system |
| 12 | B2-I-Th +A2 Removed by immune system |
| 13 | B1, B1-S-Th and B1-I-Th A1 |
| 14 | B2, B2-S-Th and B2-I-Th A2 |
| 15 | B1-S-Th and B1-I-Th B1 |
| 16 | B2-S-Th and B2-I-Th B2 |
| 17 | B1-S-Th S-Th |
| 18 | B2-S-Th S-Th |
| 19 | B1-I-Th I-Th |
| 20 | B2-I-Th I-Th |
| 21 | S + V I |
| 22 | I + A1 Removed by immune system |
| 23 | I + A2 Removed by immune system |
| 24 | I Lysis and release viruses into body fluids |
| 25 | A1 Decay |
| 26 | A2 Decay |
| 27 | B1 Decay |
| 28 | B2 Decay |
| 29 | S-Th Decay |
| 30 | S Decay |

Table S13: Reaction index and the name of each reaction in model 2.1

| Reaction index | Reaction rate |
| --- | --- |
| 1 | $\frac{\alpha_{1}\left( S-Th \right)V}{(V+k_{m})}$ |
| 2 | $\frac{\beta\left( I-Th \right)A1}{(A1+k_{m'})}$ |
| 3 | $\frac{\beta\left( I-Th \right)A2}{(A2+k_{m''})}$ |
| 4 | Forward reaction rate: $\frac{\varepsilon B1V(S-Th)}{(V+k_{m1})}$  Reverse reaction rate: $\tau(B1-S-Th)$ |
| 5 | Forward reaction rate: $\frac{\varepsilon B1V(I-Th)}{(V+k_{m1})}$  Reverse reaction rate: $\tau(B1-I-Th)$ |
| 6 | Forward reaction rate: $\frac{\varepsilon B2V(S-Th)}{(V+k_{m2})}$  Reverse reaction rate: $\tau(B2-S-Th)$ |
| 7 | Forward reaction rate: $\frac{\varepsilon B2V(I-Th)}{(V+k_{m2})}$  Reverse reaction rate: $\tau(B2-I-Th)$ |
| 8 | Forward reaction rate: $\theta_{1}A1V$  Reverse reaction rate: $\omega(A1-V)$ |
| 9 | Forward reaction rate: $\theta_{2}A2V$  Reverse reaction rate: $\omega(A2-V)$ |
| 10 | $\gamma(I-Th)$ |
| 11 | $\frac{\beta\left( B1-I-Th \right)A1}{(A1+k_{m^{'}}/\varphi)}$ |
| 12 | $\frac{\beta\left( B2-I-Th \right)A2}{(A2+k_{m''}/\varphi)}$ |
| 13 | $\rho(B1-I-Th+B1+B1-S-Th)$ |
| 14 | $\rho(B2-I-Th+B2+B2-S-Th)$ |
| 15 | $\delta(B1-I-Th+B1-S-Th)$ |
| 16 | $\delta(B2-I-Th+B2-S-Th)$ |
| 17 | $\delta(B1-S-Th)$ |
| 18 | $\delta(B2-S-Th)$ |
| 19 | $\delta(B1-I-Th)$ |
| 20 | $\delta(B2-I-Th)$ |
| 21 | $\frac{\alpha_{2}\mathrm{SV}}{(V+k_{m})}$ |
| 22 | $\frac{\beta IA1}{(A1+k_{m'})}$ |
| 23 | $\frac{\beta IA2}{(A2+k_{m''})}$ |
| 24 | $\gamma I$ |
| 25 | $\mu_{1}A1$ |
| 26 | $\mu_{1}A2$ |
| 27 | $\mu_{1}B1$ |
| 28 | $\mu_{1}B2$ |
| 29 | $\mu_{2}(S-Th)$ |
| 30 | $\mu_{2}S$ |

Table S14: Reaction rate of each reaction in model 2.1

| Parameter Name | Description | Value | Range |
| --- | --- | --- | --- |
| $\alpha_{1}$ | Maximal infection rate toward effect Th-cells | 1e-2 for effector Th-cell infection; 0 for control group | [1e-3~1e-2] |
| $\alpha_{2}$ | Maximal infection rate toward overall susceptible cells | 1e-2 | [1e-3~1e-2] |
| $\beta$ | Maximal clearance rate of Th-cell targeted ADCC effect | 3e-2 | [1e-2~5e-2] |
| $k_{m}$ | The virus concentration in 50% of maximal infection rate toward susceptible cells | 1e6 | [1e5~1e7] |
| $k_{m'}$ | The antibody-1 concentration in 50% of maximal ADCC effect | 1e5 | Is equal to 1/$\theta_{1}$ |
| $k_{m''}$ | The antibody-2 concentration in 50% of maximal ADCC effect | 1e6 | Is equal to 1/$\theta_{2}$ |
| $\varphi$ | The fold increase in antibody concentration in the germinal center compared to that in the bloodstream. | 1e2 | [1e1~1e3] |
| $\varepsilon$ | Maximal formation rate of germinal center | 2e-2 | [1e-3~5e-2] |
| $k_{m1}$ | The antibody-1 concentration in 50% of maximal ADCC effect in germinal center | 1e5 | Is equal to 1/$\theta_{1}$ |
| $k_{m2}$ | The antibody-2 concentration in 50% of maximal ADCC effect in germinal center | 1e6 | Is equal to 1/$\theta_{2}$ |
| $\tau$ | The dissociation constant of germinal center | 0.1 | [0.01~0.1] |
| $\theta_{1}$ | Forward virus binding constant of antibody-1 | 1e-5 | [1e-5~1e-8] |
| $\theta_{2}$ | Forward virus binding constant of antibody-2 | 1e-6 | [1e-5~1e-8] |
| $\omega$ | Dissociation constant of antibody-virus complex | 1e-14 | 1e-14 |
| $\gamma$ | Lysis rate of infected cell | 1e-3 | [1e-3~1e-2] |
| $\rho$ | Antibody production rate per single B-cell | 1e5 | [1e4~1e6] |
| $\delta$ | B-cell or Th-cell regeneration rate in germinal center | 1e-2 | [1e-3~5e-2] |
| $\mu_{1}$ | Degradation rate of antibody and B-cell | 1e-2 | [1e-3~5e-2] |
| $\mu_{2}$ | Degradation rate of susceptible cells | 1e-3 | [1e-3~1e-2] |
| $\pi_{0}$ | Replenish constant of B-cell | 1e-2 | $\pi_{0}=B_{0}*\mu_{1}$ |
| $\pi_{1}$ | Replenish constant of effect Th-cell | 1e2 | $\pi_{1}={S-Th}_{0}*\mu_{2}$ |
| $\pi_{2}$ | Replenish constant of overall susceptible cells | 1e5 | $\pi_{2}=S_{0}*\mu_{2}$ |
| $\vartheta$ | Clearance rate of antibody-virus complex | 0.1 | [0.01~0.5] |

Table S15: Estimates of parameters in model 2.1.

| Variables | Meaning | Initial value |
| --- | --- | --- |
| S-Th | Susceptible effect Th-cell | 1e5 |
| I-Th | Infected effect Th-cell | 0 |
| V | Virus | 1 |
| A1 | Antibody-1 | 100 |
| A2 | Antibody-2 | 100 |
| B1-S-Th | GC formed by susceptible effect Th-cell and type 1 B-cell | 0 |
| B2-S-Th | GC formed by susceptible effect Th-cell and type 2 B-cell | 0 |
| B1-I-Th | GC formed by infected effect Th-cell and type 1 B-cell | 0 |
| B2-I-Th | GC formed by infected effect Th-cell and type 2 B-cell | 0 |
| B1 | Type 1 B-cell | 1 |
| B2 | Type 2 B-cell | 1 |
| S | Overall susceptible cells | 1e8 |
| I | Overall infected cells | 0 |
| A1-V | Virus-antibody-1 complex | 0 |
| A2-V | Virus-antibody-2 complex | 0 |

Table S16: Time-dependent variables of model 2.1.

The set of ordinary differential equations is displayed below:

$\frac{d\left( S-Th \right)}{dt}=\pi_{1}- \frac{\alpha_{1}\left( S-Th \right)V}{\left( V+k_{m} \right)} -\mu_{2}\left( S-Th \right)-\frac{\varepsilon B1V\left( S-Th \right)}{\left( V+k_{m1} \right)}-\frac{\varepsilon B2V\left( S-Th \right)}{\left( V+k_{m2} \right)}+\tau\left( B1-S-Th \right)+\delta\left( B1-S-Th \right) +\tau\left( B2-S-Th \right)+\delta\left( B2-S-Th \right)$

$$\frac{d\left( I-Th \right)}{dt}=\frac{\alpha_{1}\left( S-Th \right)V}{\left( V+k_{m} \right)}- \frac{\beta\left( I-Th \right)A1}{\left( A1+k_{m^{'}} \right)}- \frac{\beta\left( I-Th \right)A2}{\left( A2+k_{m^{''}} \right)}-\gamma\left( I-Th \right) -\frac{\varepsilon B1V\left( I-Th \right)}{\left( V+k_{m1} \right)}+\delta\left( B1-I-Th \right)+\frac{\varepsilon B2V(I-Th)}{(V+k_{m2})}+\delta\left( B2-I-Th \right)$$

$\frac{d\left( V \right)}{dt}= \vartheta\gamma\left( \left( I-Th \right)+I \right)$-$\theta_{1}A1V+\omega\left( A1-V \right)- \theta_{2}A2V+\omega\left( A2-V \right)$

$\frac{d(A1)}{dt}=- \theta_{1}A1V+\omega\left( A1-V \right)- \mu_{1}A1$+ $\rho(B1-I-Th+B1+B1-S-Th)$

$\frac{d(A2)}{dt}=- \theta_{2}A2V+\omega\left( A2-V \right)- \mu_{1}A2$+ $\rho(B2-I-Th+B2+B2-S-Th)$

$$\frac{d(B1-S-Th)}{dt}=-\frac{\alpha_{1}\left( B1-S-Th \right)V}{\left( V+k_{m} \right)} + \frac{\varepsilon B1V\left( S-Th \right)}{\left( V+k_{m1} \right)}- \tau(B1-S-Th)$$

$$\frac{d(B2-S-Th)}{dt}=-\frac{\alpha_{1}\left( B2-S-Th \right)V}{\left( V+k_{m} \right)} + \frac{\varepsilon B2V\left( S-Th \right)}{\left( V+k_{m2} \right)}- \tau(B2-S-Th)$$

$$\frac{d(B1-I-Th)}{dt}=\frac{\alpha_{1}\left( B1-S-Th \right)V}{\left( V+k_{m} \right)}+ \frac{\varepsilon B1V(I-Th)}{(V+k_{m1})}- \frac{\beta\left( B1-I-Th \right)A1}{\left( A1+\frac{k_{m^{'}}}{\varphi} \right)}-\tau\left( B1-I-Th \right)$$

$$\frac{d(B2-I-Th)}{dt}=\frac{\alpha_{1}\left( B2-S-Th \right)V}{\left( V+k_{m} \right)}+ \frac{\varepsilon B2V(I-Th)}{(V+k_{m2})}- \frac{\beta\left( B2-I-Th \right)A2}{\left( A2+\frac{k_{m^{''}}}{\varphi} \right)}-\tau\left( B2-I-Th \right)$$

$\frac{d(B1)}{dt}=\pi_{0}+\delta\left( B1-I-Th+B1-S-Th \right)-\mu_{1}B1-\frac{\varepsilon B1V\left( S-Th \right)}{\left( V+k_{m1} \right)}-\frac{\varepsilon B1V\left( I-Th \right)}{\left( V+k_{m1} \right)}+\tau\left( B1-I-Th \right)+\tau\left( B1-S-Th \right)$

$\frac{d(B2)}{dt}=\pi_{0}+\delta\left( B2-I-Th+B2-S-Th \right)-\mu_{1}B2-\frac{\varepsilon B2V\left( S-Th \right)}{\left( V+k_{m2} \right)}-\frac{\varepsilon B2V\left( I-Th \right)}{\left( V+k_{m2} \right)}+\tau\left( B2-I-Th \right)+\tau\left( B2-S-Th \right)$

$$\frac{d(S)}{dt}=\pi_{2}- \frac{\alpha_{2}\mathrm{SV}}{\left( V+k_{m} \right)} -\mu_{2}S$$

$\frac{d(I)}{dt}=\frac{\alpha_{2}\mathrm{SV}}{\left( V+k_{m} \right)}- \frac{\beta IA1}{\left( A1+k_{m^{'}} \right)} - \frac{\beta IA2}{\left( A2+k_{m''} \right)} - \gamma I$

$$\frac{d\left( A1-V \right)}{dt}=\theta_{1}A1V-\omega\left( A1-V \right)- \vartheta(A1-V)$$

$$\frac{d\left( A2-V \right)}{dt}=\theta_{2}A2V-\omega\left( A2-V \right)- \vartheta(A2-V)$$

Model 2.2

Model 2.2 is designed to simulate the differences in HIV inhibitory effects of antibodies with different specificities. Specifically, RBD-specific neutralizing antibodies targeting the virus CD4 binding site can render the virus incapable of infecting cells after binding. Non-RBD antibodies, on the other hand, lack this capability.

| Reaction Index | Reaction |
| --- | --- |
| 1 | S-Th + V I-Th |
| 2 | I-Th +A Removed by immune system |
| 3 | B + V + S-Th B-S-Th |
| 4 | B + V + I-Th B-I-Th |
| 5 | A + V A-V |
| 6 | I-Th Lysis and release viruses into body fluids |
| 7 | B-I-Th +A Removed by immune system |
| 8 | B, B-S-Th and B-I-Th A |
| 9 | B-S-Th and B-I-Th B |
| 10 | B-S-Th S-Th |
| 11 | B-I-Th I-Th |
| 12 | S + V I |
| 13 | I + A Removed by immune system |
| 14 | I Lysis and release viruses into body fluids |
| 15 | A Decay |
| 16 | B Decay |
| 17 | S-Th Decay |
| 18 | S Decay |

Table S17: Reaction index and the name of each reaction in model 2.2

| Reaction index | Reaction rate |
| --- | --- |
| 1 | $\frac{\alpha\left( S-Th \right)V}{(V+k_{m})}$ |
| 2 | $\frac{\beta\left( I-Th \right)A}{(A+k_{m'})}$ |
| 3 | Forward reaction rate: $\frac{\varepsilon BV(S-Th)}{(V+k_{m1})}$  Reverse reaction rate: $\tau(B-S-Th)$ |
| 4 | Forward reaction rate: $\frac{\varepsilon BV(I-Th)}{(V+k_{m1})}$  Reverse reaction rate: $\tau(B-I-Th)$ |
| 5 | Forward reaction rate: $\theta AV$  Reverse reaction rate: $\omega(A-V)$ |
| 6 | $\gamma(I-Th)$ |
| 7 | $\frac{\beta\left( B-I-Th \right)A}{(A+k_{m^{'}}/\varphi)}$ |
| 8 | $\rho(B-I-Th+B+B-S-Th)$ |
| 9 | $\delta(B-I-Th+B-S-Th)$ |
| 10 | $\delta(B-S-Th)$ |
| 11 | $\delta(B-I-Th)$ |
| 12 | $\frac{\alpha SV}{(V+k_{m})}$ |
| 13 | $\frac{\beta IA}{(A+k_{m'})}$ |
| 14 | $\gamma I$ |
| 15 | $\mu_{1}A$ |
| 16 | $\mu_{1}B$ |
| 17 | $\mu_{2}(S-Th)$ |
| 18 | $\mu_{2}S$ |

Table S18: Reaction rate of each reaction in model 2.2

| Parameter Name | Description | Value | Range |
| --- | --- | --- | --- |
| $\alpha$ | Maximal infection rate toward overall susceptible cells | 1e-2 | [1e-3~1e-2] |
| $\beta$ | Maximal clearance rate of Th-cell targeted ADCC effect | 3e-2 | [1e-2~5e-2] |
| $k_{m}$ | The virus concentration in 50% of maximal infection rate toward susceptible cells | 1e6 | [1e5~1e7] |
| $k_{m'}$ | The antibody concentration in 50% of maximal ADCC effect | 1e5 | Is equal to 1/$\theta$ |
| $\varphi$ | The fold increase in antibody concentration in the germinal center compared to that in the bloodstream. | 1e2 | [1e1~1e3] |
| $\varepsilon$ | Maximal formation rate of germinal center | 2e-2 | [1e-3~5e-2] |
| $k_{m1}$ | The antibody concentration in 50% of maximal ADCC effect in germinal center | 1e5 | Is equal to 1/$\theta$ |
| $\tau$ | The dissociation constant of germinal center | 0.1 | [0.01~0.1] |
| $\theta$ | Forward virus binding constant of antibody | 1e-5 | [1e-5~1e-8] |
| $\omega$ | Dissociation constant of antibody-virus complex | 1e-14 | 1e-14 |
| $\gamma$ | Lysis rate of infected cell | 1e-3 | [1e-3~1e-2] |
| $\rho$ | Antibody production rate per single B-cell | 1e5 | [1e4~1e6] |
| $\delta$ | B-cell or Th-cell regeneration rate in germinal center | 1e-2 | [1e-3~5e-2] |
| $\mu_{1}$ | Degradation rate of antibody and B-cell | 1e-2 | [1e-3~5e-2] |
| $\mu_{2}$ | Degradation rate of susceptible cells | 1e-3 | [1e-3~1e-2] |
| $\pi_{0}$ | Replenish constant of B-cell | 1e-2 | $\pi_{0}=B_{0}*\mu_{1}$ |
| $\pi_{1}$ | Replenish constant of effect Th-cell | 1e2 | $\pi_{1}={S-Th}_{0}*\mu_{2}$ |
| $\pi_{2}$ | Replenish constant of overall susceptible cells | 1e5 | $\pi_{2}=S_{0}*\mu_{2}$ |
| $\vartheta$ | Clearance rate of antibody-virus complex | 0.1 | [0.01~0.5] |

Table S19: Estimates of parameters in model 2.2.

| Variables | Meaning | Initial value |
| --- | --- | --- |
| S-Th | Susceptible effect Th-cell | 1e4 |
| I-Th | Infected effect Th-cell | 0 |
| V | Virus | 1 |
| A | Antibody | 100 |
| B-S-Th | GC formed by susceptible effect Th-cell and B-cell | 0 |
| B-I-Th | GC formed by infected effect Th-cell and B-cell | 0 |
| B | B-cell | 1 |
| S | Overall susceptible cells | 1e8 |
| I | Overall infected cells | 0 |
| A-V | Virus-antibody complex | 0 |

Table S20: Time-dependent variables of model 2.2.

The ordinary differential equations of RBD-neutralizing antibody are represented below:

$$\frac{d\left( S-Th \right)}{dt}=\pi_{1}- \frac{\alpha\left( S-Th \right)V}{\left( V+k_{m} \right)} -\mu_{2}\left( S-Th \right)-\frac{\varepsilon BV\left( S-Th \right)}{\left( V+k_{m1} \right)}+\tau\left( B-S-Th \right)+\delta\left( B-S-Th \right)$$

$\frac{d\left( I-Th \right)}{dt}=\frac{\alpha\left( S-Th \right)V}{\left( V+k_{m} \right)}- \frac{\beta\left( I-Th \right)A}{\left( A+k_{m^{'}} \right)}-\gamma\left( I-Th \right)-\frac{\varepsilon BV(I-Th)}{(V+k_{m1})}+\delta\left( B-I-Th \right)$

$$\frac{d\left( V \right)}{dt}= \vartheta\gamma\left( \left( I-Th \right)+I \right)- \theta AV+\omega\left( A-V \right)$$

$\frac{d(A)}{dt}=- \theta AV+\omega\left( A-V \right)- \mu_{1}A+\rho(B-I-Th+B+B-S-Th)$

$$\frac{d(B-S-Th)}{dt}=-\frac{\alpha\left( B-S-Th \right)V}{\left( V+k_{m} \right)} + \frac{\varepsilon BV\left( S-Th \right)}{\left( V+k_{m1} \right)}- \tau(B-S-Th)$$

$$\frac{d(B-I-Th)}{dt}=\frac{\alpha\left( B-S-Th \right)V}{\left( V+k_{m} \right)}+ \frac{\varepsilon BV(I-Th)}{(V+k_{m1})}- \frac{\beta\left( B-I-Th \right)A}{\left( A+\frac{k_{m^{'}}}{\varphi} \right)}-\tau\left( B-I-Th \right)$$

$\frac{d(B)}{dt}=\pi_{0}+\delta\left( B-I-Th+B-S-Th \right)-\mu_{1}B-\frac{\varepsilon BV\left( S-Th \right)}{\left( V+k_{m1} \right)}-\frac{\varepsilon BV\left( I-Th \right)}{\left( V+k_{m1} \right)}+\tau\left( B-I-Th \right)+\tau\left( B-S-Th \right)$

$$\frac{d(S)}{dt}=\pi_{2}- \frac{\alpha SV}{\left( V+k_{m} \right)} -\mu_{2}S$$

$\frac{d(I)}{dt}=\frac{\alpha SV}{\left( V+k_{m} \right)}- \frac{\beta IA}{\left( A+k_{m^{'}} \right)} - \gamma I$

The ordinary differential equations of non-RBD-neutralizing antibody are represented below:

$$\frac{d\left( S-Th \right)}{dt}=\pi_{1}- \frac{\alpha\left( S-Th \right)(V+(A-V))}{\left( V+(A-V)+k_{m} \right)} -\mu_{2}\left( S-Th \right)-\frac{\varepsilon BV\left( S-Th \right)}{\left( V+k_{m1} \right)}+\tau\left( B-S-Th \right)+\delta\left( B-S-Th \right)$$

$\frac{d\left( I-Th \right)}{dt}=\frac{\alpha\left( S-Th \right)(V+(A-V))}{\left( (V+(A-V))+k_{m} \right)}- \frac{\beta\left( I-Th \right)A}{\left( A+k_{m^{'}} \right)}-\gamma\left( I-Th \right)-\frac{\varepsilon BV(I-Th)}{(V+k_{m1})}+\delta\left( B-I-Th \right)$

$$\frac{d\left( V \right)}{dt}= \vartheta\gamma\left( \left( I-Th \right)+I \right)- \theta AV+\omega\left( A-V \right)$$

$\frac{d(A)}{dt}=- \theta AV+\omega\left( A-V \right)- \mu_{1}A+\rho(B-I-Th+B+B-S-Th)$

$$\frac{d(B-S-Th)}{dt}=-\frac{\alpha\left( B-S-Th \right)(V+(A-V))}{\left( (V+(A-V))+k_{m} \right)} + \frac{\varepsilon BV\left( S-Th \right)}{\left( V+k_{m1} \right)}- \tau(B-S-Th)$$

$$\frac{d(B-I-Th)}{dt}=\frac{\alpha\left( B-S-Th \right)(V+(A-V))}{\left( (V+(A-V))+k_{m} \right)}+ \frac{\varepsilon BV(I-Th)}{(V+k_{m1})}- \frac{\beta\left( B-I-Th \right)A}{\left( A+\frac{k_{m^{'}}}{\varphi} \right)}-\tau\left( B-I-Th \right)$$

$\frac{d(B)}{dt}=\pi_{0}+\delta\left( B-I-Th+B-S-Th \right)-\mu_{1}B-\frac{\varepsilon BV\left( S-Th \right)}{\left( V+k_{m1} \right)}-\frac{\varepsilon BV\left( I-Th \right)}{\left( V+k_{m1} \right)}+\tau\left( B-I-Th \right)+\tau\left( B-S-Th \right)$

$$\frac{d(S)}{dt}=\pi_{2}- \frac{\alpha S(V+(A-V))}{\left( (V+(A-V))+k_{m} \right)} -\mu_{2}S$$

$\frac{d(I)}{dt}=\frac{\alpha S(V+(A-V))}{\left( (V+(A-V))+k_{m} \right)}- \frac{\beta IA}{\left( A+k_{m^{'}} \right)} - \gamma I$

Model 2.3

Model 2.3 is utilized to simulate the impact of variations in the strength of ADCC effect on infection in a complex model. In this case, a strong ADCC effect is represented by a larger β value of 0.03, while the weak ADCC control group has a smaller β value of 0.015. Model 2.3 is employed for generating Figure 11. Interestingly, the simulation indicates that, for the same type of antibody, a weaker ADCC effect actually benefits virus control and the generation of elite controllers.

| Reaction Index | Reaction |
| --- | --- |
| 1 | S-Th + V I-Th |
| 2 | I-Th +A Removed by immune system |
| 3 | B + V + S-Th B-S-Th |
| 4 | B + V + I-Th B-I-Th |
| 5 | A + V A-V |
| 6 | I-Th Lysis and release viruses into body fluids |
| 7 | B-I-Th +A Removed by immune system |
| 8 | B, B-S-Th and B-I-Th A |
| 9 | B-S-Th and B-I-Th B |
| 10 | B-S-Th S-Th |
| 11 | B-I-Th I-Th |
| 12 | S + V I |
| 13 | I + A Removed by immune system |
| 14 | I Lysis and release viruses into body fluids |
| 15 | A Decay |
| 16 | B Decay |
| 17 | S-Th Decay |
| 18 | S Decay |

Table S21: Reaction index and the name of each reaction in model 2.3

| Reaction index | Reaction rate |
| --- | --- |
| 1 | $\frac{\alpha\left( S-Th \right)V}{(V+k_{m})}$ |
| 2 | $\frac{\beta\left( I-Th \right)A}{(A+k_{m'})}$ |
| 3 | Forward reaction rate: $\frac{\varepsilon BV(S-Th)}{(V+k_{m1})}$  Reverse reaction rate: $\tau(B-S-Th)$ |
| 4 | Forward reaction rate: $\frac{\varepsilon BV(I-Th)}{(V+k_{m1})}$  Reverse reaction rate: $\tau(B-I-Th)$ |
| 5 | Forward reaction rate: $\theta AV$  Reverse reaction rate: $\omega(A-V)$ |
| 6 | $\gamma(I-Th)$ |
| 7 | $\frac{\beta\left( B-I-Th \right)A}{(A+k_{m^{'}}/\varphi)}$ |
| 8 | $\rho(B-I-Th+B+B-S-Th)$ |
| 9 | $\delta(B-I-Th+B-S-Th)$ |
| 10 | $\delta(B-S-Th)$ |
| 11 | $\delta(B-I-Th)$ |
| 12 | $\frac{\alpha SV}{(V+k_{m})}$ |
| 13 | $\frac{\beta IA}{(A+k_{m'})}$ |
| 14 | $\gamma I$ |
| 15 | $\mu_{1}A$ |
| 16 | $\mu_{1}B$ |
| 17 | $\mu_{2}(S-Th)$ |
| 18 | $\mu_{2}S$ |

Table S22: Reaction rate of each reaction in model 2.3

| Parameter Name | Description | Value | Range |
| --- | --- | --- | --- |
| $\alpha$ | Maximal infection rate toward overall susceptible cells | 1e-2 | [1e-3~1e-2] |
| $\beta$ | Maximal clearance rate of Th-cell targeted ADCC effect | 3e-2 for strong ADCC; 1.5e-2 for weak ADCC | [1e-2~5e-2] |
| $k_{m}$ | The virus concentration in 50% of maximal infection rate toward susceptible cells | 1e6 | [1e5~1e7] |
| $k_{m'}$ | The antibody concentration in 50% of maximal ADCC effect | 1e5 | Is equal to 1/$\theta$ |
| $\varphi$ | The fold increase in antibody concentration in the germinal center compared to that in the bloodstream. | 1e2 | [1e1~1e3] |
| $\varepsilon$ | Maximal formation rate of germinal center | 2e-2 | [1e-3~5e-2] |
| $k_{m1}$ | The antibody concentration in 50% of maximal ADCC effect in germinal center | 1e5 | Is equal to 1/$\theta$ |
| $\tau$ | The dissociation constant of germinal center | 0.1 | [0.01~0.1] |
| $\theta$ | Forward virus binding constant of antibody | 1e-5 | [1e-5~1e-8] |
| $\omega$ | Dissociation constant of antibody-virus complex | 1e-14 | 1e-14 |
| $\gamma$ | Lysis rate of infected cell | 1e-3 | [1e-3~1e-2] |
| $\rho$ | Antibody production rate per single B-cell | 1e5 | [1e4~1e6] |
| $\delta$ | B-cell or Th-cell regeneration rate in germinal center | 1e-2 | [1e-3~5e-2] |
| $\mu_{1}$ | Degradation rate of antibody and B-cell | 1e-2 | [1e-3~5e-2] |
| $\mu_{2}$ | Degradation rate of susceptible cells | 1e-3 | [1e-3~1e-2] |
| $\pi_{0}$ | Replenish constant of B-cell | 1e-2 | $\pi_{0}=B_{0}*\mu_{1}$ |
| $\pi_{1}$ | Replenish constant of effect Th-cell | 1e1 | $\pi_{1}={S-Th}_{0}*\mu_{2}$ |
| $\pi_{2}$ | Replenish constant of overall susceptible cells | 1e5 | $\pi_{2}=S_{0}*\mu_{2}$ |
| $\vartheta$ | Clearance rate of antibody-virus complex | 0.1 | [0.01~0.5] |

Table S23: Estimates of parameters in model 2.3.

| Variables | Meaning | Initial value |
| --- | --- | --- |
| S-Th | Susceptible effect Th-cell | 1e4 |
| I-Th | Infected effect Th-cell | 0 |
| V | Virus | 1 |
| A | Antibody | 100 |
| B-S-Th | GC formed by susceptible effect Th-cell and B-cell | 0 |
| B-I-Th | GC formed by infected effect Th-cell and B-cell | 0 |
| B | B-cell | 1 |
| S | Overall susceptible cells | 1e8 |
| I | Overall infected cells | 0 |
| A-V | Virus-antibody complex | 0 |

Table S24: Time-dependent variables of model 2.3.

The ordinary differential equations of model 2.3 are represented below:

$$\frac{d\left( S-Th \right)}{dt}=\pi_{1}- \frac{\alpha\left( S-Th \right)V}{\left( V+k_{m} \right)} -\mu_{2}\left( S-Th \right)-\frac{\varepsilon BV\left( S-Th \right)}{\left( V+k_{m1} \right)}+\tau\left( B-S-Th \right)+\delta\left( B-S-Th \right)$$

$\frac{d\left( I-Th \right)}{dt}=\frac{\alpha\left( S-Th \right)V}{\left( V+k_{m} \right)}- \frac{\beta\left( I-Th \right)A}{\left( A+k_{m^{'}} \right)}-\gamma\left( I-Th \right)-\frac{\varepsilon BV(I-Th)}{(V+k_{m1})}+\delta\left( B-I-Th \right)$

$$\frac{d\left( V \right)}{dt}= \vartheta\gamma\left( \left( I-Th \right)+I \right)- \theta AV+\omega\left( A-V \right)$$

$\frac{d(A)}{dt}=- \theta AV+\omega\left( A-V \right)- \mu_{1}A+\rho(B-I-Th+B+B-S-Th)$

$$\frac{d(B-S-Th)}{dt}=-\frac{\alpha\left( B-S-Th \right)V}{\left( V+k_{m} \right)} + \frac{\varepsilon BV\left( S-Th \right)}{\left( V+k_{m1} \right)}- \tau(B-S-Th)$$

$$\frac{d(B-I-Th)}{dt}=\frac{\alpha\left( B-S-Th \right)V}{\left( V+k_{m} \right)}+ \frac{\varepsilon BV(I-Th)}{(V+k_{m1})}- \frac{\beta\left( B-I-Th \right)A}{\left( A+\frac{k_{m^{'}}}{\varphi} \right)}-\tau\left( B-I-Th \right)$$

$\frac{d(B)}{dt}=\pi_{0}+\delta\left( B-I-Th+B-S-Th \right)-\mu_{1}B-\frac{\varepsilon BV\left( S-Th \right)}{\left( V+k_{m1} \right)}-\frac{\varepsilon BV\left( I-Th \right)}{\left( V+k_{m1} \right)}+\tau\left( B-I-Th \right)+\tau\left( B-S-Th \right)$

$$\frac{d(S)}{dt}=\pi_{2}- \frac{\alpha SV}{\left( V+k_{m} \right)} -\mu_{2}S$$

$\frac{d(I)}{dt}=\frac{\alpha SV}{\left( V+k_{m} \right)}- \frac{\beta IA}{\left( A+k_{m^{'}} \right)} - \gamma I$

Model 2.4

Model 2.4 is employed to simulate the inhibitory and clearance effects of antibody therapy on viruses in a complex model. At a fixed time point, the amount of exogenous antibodies added is represented by *c*, which is the dose of antibody therapy received. The value of *c* is zero for all other time points. Model 2.4 can effectively simulate the changes in concentration of various components after the addition of antibodies, including fluctuations in the virus, antibodies, susceptible cells, and infected Th cells.

| Reaction Index | Reaction |
| --- | --- |
| 1 | S-Th + V I-Th |
| 2 | I-Th +A Removed by immune system |
| 3 | B + V + S-Th B-S-Th |
| 4 | B + V + I-Th B-I-Th |
| 5 | A + V A-V |
| 6 | I-Th Lysis and release viruses into body fluids |
| 7 | B-I-Th +A Removed by immune system |
| 8 | B, B-S-Th and B-I-Th A |
| 9 | B-S-Th and B-I-Th B |
| 10 | B-S-Th S-Th |
| 11 | B-I-Th I-Th |
| 12 | S + V I |
| 13 | I + A Removed by immune system |
| 14 | I Lysis and release viruses into body fluids |
| 15 | A Decay |
| 16 | B Decay |
| 17 | S-Th Decay |
| 18 | S Decay |

Table S25: Reaction index and the name of each reaction in model 2.4

| Reaction index | Reaction rate |
| --- | --- |
| 1 | $\frac{\alpha\left( S-Th \right)V}{(V+k_{m})}$ |
| 2 | $\frac{\beta\left( I-Th \right)A}{(A+k_{m'})}$ |
| 3 | Forward reaction rate: $\frac{\varepsilon BV(S-Th)}{(V+k_{m1})}$  Reverse reaction rate: $\tau(B-S-Th)$ |
| 4 | Forward reaction rate: $\frac{\varepsilon BV(I-Th)}{(V+k_{m1})}$  Reverse reaction rate: $\tau(B-I-Th)$ |
| 5 | Forward reaction rate: $\theta AV$  Reverse reaction rate: $\omega(A-V)$ |
| 6 | $\gamma(I-Th)$ |
| 7 | $\frac{\beta\left( B-I-Th \right)A}{(A+k_{m^{'}}/\varphi)}$ |
| 8 | $\rho(B-I-Th+B+B-S-Th)$ |
| 9 | $\delta(B-I-Th+B-S-Th)$ |
| 10 | $\delta(B-S-Th)$ |
| 11 | $\delta(B-I-Th)$ |
| 12 | $\frac{\alpha SV}{(V+k_{m})}$ |
| 13 | $\frac{\beta IA}{(A+k_{m'})}$ |
| 14 | $\gamma I$ |
| 15 | $\mu_{1}A$ |
| 16 | $\mu_{1}B$ |
| 17 | $\mu_{2}(S-Th)$ |
| 18 | $\mu_{2}S$ |

Table S26: Reaction rate of each reaction in model 2.4

| Parameter Name | Description | Value | Range |
| --- | --- | --- | --- |
| $\alpha$ | Maximal infection rate toward overall susceptible cells | 1e-2 | [1e-3~1e-2] |
| $\beta$ | Maximal clearance rate of Th-cell targeted ADCC effect | 3e-2 | [1e-2~5e-2] |
| $k_{m}$ | The virus concentration in 50% of maximal infection rate toward susceptible cells | 1e6 | [1e5~1e7] |
| $k_{m'}$ | The antibody concentration in 50% of maximal ADCC effect | 1e5 | Is equal to 1/$\theta$ |
| $\varphi$ | The fold increase in antibody concentration in the germinal center compared to that in the bloodstream. | 1e2 | [1e1~1e3] |
| $\varepsilon$ | Maximal formation rate of germinal center | 2e-2 | [1e-3~5e-2] |
| $k_{m1}$ | The antibody concentration in 50% of maximal ADCC effect in germinal center | 1e5 | Is equal to 1/$\theta$ |
| $\tau$ | The dissociation constant of germinal center | 0.1 | [0.01~0.1] |
| $\theta$ | Forward virus binding constant of antibody | 1e-5 | [1e-5~1e-8] |
| $\omega$ | Dissociation constant of antibody-virus complex | 1e-14 | 1e-14 |
| $\gamma$ | Lysis rate of infected cell | 1e-3 | [1e-3~1e-2] |
| $\rho$ | Antibody production rate per single B-cell | 1e5 | [1e4~1e6] |
| $\delta$ | B-cell or Th-cell regeneration rate in germinal center | 1e-2 | [1e-3~5e-2] |
| $\mu_{1}$ | Degradation rate of antibody and B-cell | 1e-2 | [1e-3~5e-2] |
| $\mu_{2}$ | Degradation rate of susceptible cells | 1e-3 | [1e-3~1e-2] |
| $\pi_{0}$ | Replenish constant of B-cell | 1e-2 | $\pi_{0}=B_{0}*\mu_{1}$ |
| $\pi_{1}$ | Replenish constant of effect Th-cell | 1e1 | $\pi_{1}={S-Th}_{0}*\mu_{2}$ |
| $\pi_{2}$ | Replenish constant of overall susceptible cells | 1e5 | $\pi_{2}=S_{0}*\mu_{2}$ |
| $\vartheta$ | Clearance rate of antibody-virus complex | 0.1 | [0.01~0.5] |
| c | Dose of antibody treatment | 5e11 | [1e10~1e12] |

Table S27: Estimates of parameters in model 2.4.

| Variables | Meaning | Initial value |
| --- | --- | --- |
| S-Th | Susceptible effect Th-cell | 1e4 |
| I-Th | Infected effect Th-cell | 0 |
| V | Virus | 1 |
| A | Antibody | 100 |
| B-S-Th | GC formed by susceptible effect Th-cell and B-cell | 0 |
| B-I-Th | GC formed by infected effect Th-cell and B-cell | 0 |
| B | B-cell | 1 |
| S | Overall susceptible cells | 1e8 |
| I | Overall infected cells | 0 |
| A-V | Virus-antibody complex | 0 |

Table S28: Time-dependent variables of model 2.4.

The ordinary differential equations of model 2.4 are represented below:

$$\frac{d\left( S-Th \right)}{dt}=\pi_{1}- \frac{\alpha\left( S-Th \right)V}{\left( V+k_{m} \right)} -\mu_{2}\left( S-Th \right)-\frac{\varepsilon BV\left( S-Th \right)}{\left( V+k_{m1} \right)}+\tau\left( B-S-Th \right)+\delta\left( B-S-Th \right)$$

$\frac{d\left( I-Th \right)}{dt}=\frac{\alpha\left( S-Th \right)V}{\left( V+k_{m} \right)}- \frac{\beta\left( I-Th \right)A}{\left( A+k_{m^{'}} \right)}-\gamma\left( I-Th \right)-\frac{\varepsilon BV(I-Th)}{(V+k_{m1})}+\delta\left( B-I-Th \right)$

$$\frac{d\left( V \right)}{dt}= \vartheta\gamma\left( \left( I-Th \right)+I \right)- \theta AV+\omega\left( A-V \right)$$

$\frac{d(A)}{dt}=c- \theta AV+\omega\left( A-V \right)- \mu_{1}A+\rho(B-I-Th+B+B-S-Th)$

$$\frac{d(B-S-Th)}{dt}=-\frac{\alpha\left( B-S-Th \right)V}{\left( V+k_{m} \right)} + \frac{\varepsilon BV\left( S-Th \right)}{\left( V+k_{m1} \right)}- \tau(B-S-Th)$$

$$\frac{d(B-I-Th)}{dt}=\frac{\alpha\left( B-S-Th \right)V}{\left( V+k_{m} \right)}+ \frac{\varepsilon BV(I-Th)}{(V+k_{m1})}- \frac{\beta\left( B-I-Th \right)A}{\left( A+\frac{k_{m^{'}}}{\varphi} \right)}-\tau\left( B-I-Th \right)$$

$\frac{d(B)}{dt}=\pi_{0}+\delta\left( B-I-Th+B-S-Th \right)-\mu_{1}B-\frac{\varepsilon BV\left( S-Th \right)}{\left( V+k_{m1} \right)}-\frac{\varepsilon BV\left( I-Th \right)}{\left( V+k_{m1} \right)}+\tau\left( B-I-Th \right)+\tau\left( B-S-Th \right)$

$$\frac{d(S)}{dt}=\pi_{2}- \frac{\alpha SV}{\left( V+k_{m} \right)} -\mu_{2}S$$

$\frac{d(I)}{dt}=\frac{\alpha SV}{\left( V+k_{m} \right)}- \frac{\beta IA}{\left( A+k_{m^{'}} \right)} - \gamma I$

References

1. Xu, Zhaobin, et al. "A Novel Mathematical Model That Predicts the Protection Time of SARS-CoV-2 Antibodies." Viruses 15.2 (2023): 586.
2. Xu, Zhaobin, et al. "More or less deadly? A mathematical model that predicts SARS-CoV-2 evolutionary direction." Computers in Biology and Medicine 153 (2023): 106510.
3. Xu, Zhaobin, et al. "An agent-based model with antibody dynamics information in COVID-19 epidemic simulation." Infectious Disease Modelling 8.4 (2023): 1151-1168.
